# Brain functional network connectivity interpolation characterizes the neuropsychiatric continuum and heterogeneity

**DOI:** 10.1101/2024.11.13.623318

**Authors:** Xinhui Li, Eloy Geenjaar, Zening Fu, Godfrey D. Pearlson, Vince D. Calhoun

## Abstract

Psychiatric and neurodevelopmental disorders such as schizophrenia (SZ) and autism spectrum disorder (ASD) are challenging to characterize in part due to their heterogeneous presentation in individuals, with symptoms now believed to exist on a continuum. Conventional diagnostic and neuroimaging analytical approaches rely on subjective assessment or group differences but typically ignore progression between groups or heterogeneity within a group. To estimate the neuropsychiatric continuum and heterogeneity, we propose a functional network connectivity (FNC) interpolation framework based on a variational autoencoder (VAE) using static FNC (sFNC) and dynamic FNC (dFNC) data from controls and patients with SZ or ASD. We demonstrate that VAEs significantly outperform a linear baseline and a semi-supervised counterpart. For both sFNC and dFNC interpolation, the generated results effectively capture representative and generalizable properties in the original data. The interpolated continua from controls to patients in both disorders reveal group-wise gradients characterized by reduced positive correlations within the auditory, sensorimotor, and visual networks, as well as between the subcortical and cerebellar domains. In contrast, anti-correlations weaken between the subcortical domain and sensory domains, and between the cerebellar domain and sensory regions. Finally, we show examples of how to generate continuous FNC data following pathological or state-based trajectories in the VAE latent space. The proposed framework offers added advantages over traditional methods, including data-driven discovery of hidden relationships, visualization of individual differences, imputation of missing values along a continuous spectrum, and estimation of the stage where an individual falls within the continuum.

## 1 Introduction

*“Happy individuals are all alike; every unhappy individual is unhappy in their own way*.*”*

*– adapted from Anna Karenina by Leo Tolstoy*

Psychiatric and neurodevelopmental disorders such as schizophrenia (SZ) and autism spectrum disorder (ASD) are recognized as leading causes of the global disease burden, affecting hundreds of millions of individuals worldwide (Collaborators et al., 2022). SZ is a lifelong mental disorder characterized by symptoms such as delusions, hallucinations, disorganized speech, catatonic behavior, and diminished emotional expression (McCutcheon et al., 2020). ASD is a developmental disability defined by social communication difficulties, and repetitive and restrictive behaviors (Lord et al., 2018). SZ and ASD show overlapping features, including genetic risk factors (O’Connell et al., 2018; The Autism Spectrum Disorders Working Group of The Psychiatric Genomics Consortium, 2017), symptom profiles (Barneveld et al., 2011; Konstantareas & Hewitt, 2001), and brain abnormalities (H. Chen et al., 2017; Du et al., 2021; Fu et al., 2021; Yoshihara et al., 2020). Moreover, both SZ and ASD are known to be highly heterogeneous across individuals (Masi et al., 2017; McCutcheon et al., 2020). Namely, individuals within the disorder group may differ in various aspects, such as genetic traits (Abrahams & Geschwind, 2008; Bill & Geschwind, 2009; Di Biase et al., 2022; Geschwind et al., 2015; Pulver et al., 2000), clinical symptoms (Ahmed et al., 2018; Georgiades et al., 2013; Kim et al., 2016; Mohr et al., 2004; Morales-Hidalgo et al., 2018), brain structure and function (Alnæs et al., 2019; Amaral et al., 2008; Brugger & Howes, 2017; Hahamy et al., 2015; Hong et al., 2018; Marín, 2012), cognitive function (Bora, 2016; Feczko et al., 2018; Joyce & Roiser, 2007), developmental trajectories (Carpenter Jr & Kirkpatrick, 1988; Fountain et al., 2012; Lord et al., 2015), and other relevant characteristics. In addition, psychiatric symptoms, such as hallucinations and delusions, may exist along a multidimensional continuum in the general population, varying in severity, frequency, and degree of conviction (DeRosse & Karlsgodt, 2015; Johns & Van Os, 2001; Morales-Hidalgo et al., 2018; Strauss, 1969; Van Os et al., 2000, 2009; Verdoux & van Os, 2002). Estimating such a continuous spectrum and imputing missing values on the continuum remains underexplored in the neuropsychiatric literature. Considering high prevalence and severity of both disorders, as well as their similarity and complexity, it is important to develop reliable diagnostic and analytical methods to characterize the continuum and heterogeneity of SZ and ASD.

However, there are several limitations in conventional evaluation approaches. First, the current diagnostic system based on the Diagnostic and Statistical Manual of Mental Disorders Fifth Edition (DSM-V) (American Psychiatric Association, 2013) relies on expert assessment or patient self-report data. Such subjective measures are insufficient to accurately capture biological deficits and symptom progression of psychiatric disorders, hindering the development of more effective treatments (Craddock & Mynors-Wallis, 2014; Cuthbert & Insel, 2013; Scott & Nelson, 2024; Su et al., 2020). Thus, it is critical to discover objective biomarkers for psychosis characterization. Recent neuroimaging studies have shown that SZ and ASD can be characterized by resting-state static and dynamic brain connectivity, derived from functional magnetic resonance imaging (fMRI) data (H. Chen et al., 2017; Damaraju et al., 2014; Du et al., 2021; Fu et al., 2021; Guo et al., 2019; Hahamy et al., 2015; Hull et al., 2017; Iraji et al., 2019, 2021; Yan et al., 2024; Yoshihara et al., 2020; Yu et al., 2012). Second, group average approaches have been widely used to analyze patient versus control functional connectivity patterns (Damaraju et al., 2014; Du et al., 2020; Y. He et al., 2020; Rabany et al., 2019). Yet, given the heterogeneous nature of SZ and ASD, group averages alone are not sufficient to characterize their multidimensional continua or multifaceted heterogeneity (Segal et al., 2024). Third, supervised learning methods, which utilize diagnostic labels, have been widely developed to predict disorder conditions from functional connectivity data (C. P. Chen et al., 2015; Rabany et al., 2019; Rashid et al., 2016; Zeng et al., 2018). As a result, the success of supervised learning models inherently depends on the validity of diagnostic labels. However, these diagnostic labels derived from current psychiatric nosologies, aside from their subjectivity, indicate only the current status of a disorder and cannot predict future disorder progression for early intervention (Scott & Nelson, 2024). Additionally, binary or multi-class supervised classification approaches alone neither explain individual differences within a group nor estimate psychotic continua between groups. To address these challenges, it is necessary to apply *unsupervised* learning approaches to brain imaging data to help characterize objective biomarkers, individual variability, and psychosis continua.

Variational autoencoders (VAEs) (Kingma & Welling, 2014) are one type of generative model that can learn data distributions in an *unsupervised* manner. A VAE consists of an encoder and a decoder. The encoder projects input data onto a latent space, and the decoder reconstructs the input data from the learned latent distributions. There are two key benefits of using a VAE. First, the encoder can perform *nonlinear* dimension reduction and approximate low-dimensional latent distributions from high-dimensional data. Second, the decoder can generate continuous synthetic data that closely resemble the observed data by sampling from the learned distributions. Recent studies have demonstrated that latent representations of resting-state fMRI data learned by a VAE can be used to identify SZ (Geenjaar et al., 2021) and ASD (F. Zhang et al., 2022), and characterize spatiotemporal dynamics (X. Zhang et al., 2021). Traditional dimension reduction approaches such as principal component analysis (PCA) (Jolliffe, 1986) and independent component analysis (ICA) (Hyvärinen & Oja, 2000) cannot naturally handle missing values, and thus cannot infer the continuous spectrum from the data. Probabilistic PCA (PPCA) has been developed to address the limitation of PCA by estimating a probabilistic model of observed data (Tipping & Bishop, 1999), but it cannot estimate complex nonlinear relationships due to its linear nature. Deep learning models such as autoencoders nonlinearly compress the data into latent features but do not provide a probabilistic estimate of the data continuum. Other deep generative models also have limitations in this specific task. For example, generative adversarial networks (GANs) (Goodfellow et al., 2020) are known for mode collapse (Che et al., 2016; Salimans et al., 2016) and instability issues (Barnett, 2018). Diffusion models (Ho et al., 2020) learn latent variables of the same dimensionality as the original data, rather than learning a low-dimensional, interpretable latent space. In addition, inference in diffusion models is computationally intensive, which typically requires hundreds of denoising steps. In contrast, the VAE provides a low-dimensional manifold that allows effective *interpretation* and efficient *interpolation* of latent features, making it a promising architecture for estimating psychotic continuum and characterizing individual variability.

Here, we propose an interpretable and generative unsupervised learning framework based on a VAE to interpolate within latent spaces learned from static functional network connectivity (sFNC) and dynamic functional network connectivity (dFNC) derived from resting-state fMRI. Interpolation – mathematically defined as the process of estimating an unknown value based on surrounding known values – can be applied to estimate an FNC matrix from its neighboring FNC matrices, assuming continuous changes between subjects. For sFNC interpolation, individual sFNC matrices are organized on a two-dimensional (2D) grid based on their similarity. A trained VAE is subsequently used to interpolate the sFNC matrices by sampling from the latent distributions along this 2D grid. For dFNC interpolation, we train the VAE using all windowed dFNC matrices from the training subjects, performed *k*-means clustering (Hartigan & Wong, 1979) on the VAE latent features, and then generated dynamic states based on the *k*-means clusters. For comparison, dynamic states are also identified by applying the *k*-means clustering algorithm on the dFNC matrices directly.

Our study highlights the benefits of maximizing data transparency to visualize individual differences within a group and continuous changes between groups. Crucially, sFNC interpolation facilitates examining individual differences and disorder continua, while dFNC interpolation captures generalizable group-specific dynamic states and intermediate state transition patterns, providing both static and dynamic comprehensive views of the psychiatric illnesses. Moreover, the VAE latent space offers the flexibility to interpolate FNC patterns between samples in the original dataset. We provide examples of how unsupervised generative models can be used to learn new insights by interpolating FNC across subjects, groups, and dynamic states.

## 2 Methods

### 2.1 Data

#### 2.1.1 Datasets

We used two resting-state fMRI datasets: the Functional Biomedical Informatics Research Network (FBIRN) dataset (Keator et al., 2016) and the initial release of the Autism Brain Imaging Data Exchange (ABIDE I) dataset (Di Martino et al., 2014).

The FBIRN fMRI scans were acquired using a standard gradient-echo echo-planar imaging (EPI) paradigm with repetition time (TR) = 2s, echo time (TE) = 30ms, flip angle (FA) = 77^◦^, 162 volumes, 32 sequential ascending axial slices of 4mm thickness and 1mm skip. Participants had their eyes closed during the scan. In addition to fMRI data, we utilized Positive and Negative Syndrome Scale (PANSS) scores (Kay et al., 1987), with positive and negative subscale scores summed separately. We also used six cognitive domain scores (speed of processing, attention/vigilance, working memory, verbal learning, visual learning, and reasoning/problem solving), as well as composite scores derived from the Computerized Multiphasic Interactive Neurocognitive System (CMINDS) (van Erp et al., 2015). All CMINDS cognitive scores were z-scored based on the mean and standard deviation of the control group. The original FBIRN dataset includes 311 subjects. 45 subjects were excluded because they had one or more missing or invalid diagnosis and cognitive scores. 266 subjects with valid subject measures were used in the subsequent analysis, including 125 subjects labeled as SZ (age mean *±* s.d.: 38.88 *±* 11.19 years; 95 males, 30 females) and 141 controls (CTR) (age mean *±* s.d.: 37.15 *±* 11.03 years; 103 males, 38 females).

The original ABIDE I dataset includes 869 subjects. We excluded 107 subjects without valid clinical assessment scores. Then we used 266 subjects with valid clinical assessment scores from three sites, including 133 subjects labeled as ASD (age mean *±* s.d.: 17.01 *±* 7.20 years; 119 males, 14 females) and 133 controls (age mean *±* s.d.: 16.05 *±* 5.67 years; 104 males, 29 females). The site-specific demographics and acquisition parameters of the ABIDE I participants are presented in Appendix A.

The statistics of subject measures in each dataset are shown in Appendix B. For each dataset, 85% of subjects were used as the training set (*N*_train_ = 225) and 15% of subjects with balanced labels were used as the holdout test set (*N*_test_ = 41). The detailed demographic information of the subjects used in this study is described in Table 1.

**Table 1:** Dataset demographic information. Demographics in the FBIRN and ABIDE I datasets are shown below. (*N*: number of total subjects; *N*_male_: number of male subjects; *N*_female_: number of female subjects; *N*_train_: number of subjects in the training set; *N*_test_: number of subjects in the test set; s.d.: standard deviation.)

| Dataset | Label | $N$ | $N_{\text{male}}$ | $N_{\text{female}}$ | $N_{\text{train}}$ | $N_{\text{test}}$ | Age mean ± s.d. (years) | Age range (years) |
| --- | --- | --- | --- | --- | --- | --- | --- | --- |
| FBIRN | SZ | 125 | 95 | 30 | 105 | 20 | $38.88 \pm 11.19$ | 18 – 62 |
| FBIRN | CTR | 141 | 103 | 38 | 120 | 21 | $37.15 \pm 11.03$ | 19 – 59 |
| ABIDE I | ASD | 133 | 119 | 14 | 113 | 20 | $17.01 \pm 7.20$ | 7 – 39 |
| ABIDE I | CTR | 133 | 104 | 29 | 112 | 21 | $16.05 \pm 5.67$ | 7 – 32 |

#### 2.1.2 Data preprocessing

The fMRI data were preprocessed using the statistical parametric mapping toolbox (SPM12, http://www.fil.ion.ucl.ac.uk/spm/) (Ashburner et al., 2014) in the MATLAB 2016 environment. We removed the first five scans for signal equilibrium and for participants’ adaptation to the scanner environment. We then performed rigid body motion correction using SPM, followed by slice timing correction. Subsequently, the fMRI data were registered to the standard Montreal Neurological Institute (MNI) space using an EPI template and resampled to 3 *×* 3 *×* 3mm^3^isotropic voxels. The resampled fMRI data were further smoothed using a Gaussian kernel with a full width at half maximum (FWHM) = 6mm.

#### 2.1.3 NeuroMark functional network connectivity

We applied the fully automated NeuroMark pipeline (Du et al., 2020) to the preprocessed fMRI data to extract subject-specific functional components and their corresponding time courses (TCs). NeuroMark, implemented in the group ICA of fMRI toolbox (GIFT; http://trendscenter.org/software/gift) leverages network templates (available in GIFT and at http://trendscenter.org/data) originally derived by separately computing group-level independent components (ICs) from two independent datasets of healthy individuals: the Human Connectome Project (HCP) (Van Essen et al., 2013) and the Brain Genomics Superstruct Project (GSP) (Holmes et al., 2015). ICs from the two datasets were matched based on spatial correlations, and 53 intrinsic connectivity networks (ICNs) with correlations greater than 0.4 selected as the network templates. The less noisy ICNs from the GSP dataset were selected for inclusion in the NeuroMark fMRI 1.0 template (Du et al., 2020). These 53 ICNs were then ordered and assigned to seven functional domains according to their anatomical and functional properties, including the subcortical (SC), auditory (AU), sensorimotor (SM), visual (VI), cognitive control (CC), default mode (DM), and cerebellar (CB) domains (Allen et al., 2014; Du et al., 2020). The components in this template were then used as spatial priors to perform spatially constrained ICA to compute ICNs and TCs for each subject for the FBIRN and ABIDE I datasets. The NeuroMark framework allows us to (1) estimate the ICA separately for each subject, thus preventing data leakage, (2) fully automate the ICA pipeline, and (3) estimate ICNs for each subject in the FBIRN and ABIDE I datasets which are aligned and comparable. Four postprocessing steps were performed on the extracted TCs, including detrending, removing head motion, despiking, and filtering with a 0.15Hz low-pass filter.

Static functional network connectivity (sFNC) was subsequently computed as the pairwise Pearson correlation between the TCs of 53 ICNs, leading to a 53*×*53 matrix. Dynamic functional network connectivity (dFNC) was estimated using a sliding window approach and a graphical LASSO method (Friedman et al., 2008). A tapered window to segment the TCs was created by convolving a rectangle (width = 20, TR = 2s, TRs = 40s) with a Gaussian kernel (*σ* = 3TRs). The window width covered 20 TRs with a step size of 1 TR, leading to 137 windows in the FBIRN dFNC data and 168 windows in the ABIDE I dFNC data. For further details on data processing, please refer to Fu et al., 2021.

### 2.2 Variational autoencoders

We used an unsupervised generative model, a variational autoencoder (VAE) (Kingma & Welling, 2014), as the backbone of the interpolation framework. We trained one VAE for each dataset (FBIRN or ABIDE I) and each data type (sFNC or dFNC). Here, we briefly explain the VAE objective function and provide a step-by-step derivation in Appendix C.

Let 𝒟_sFNC = {_x^(1)^, …, x^(*i*)^, …, x^(*N*)^} denote a dataset of *N* sFNC samples x^(*i*)^ ∈ ℝ^*V*^, where *N* is the number of subjects and *V* is the number of features in the sample. Each sample x^(*i*)^is the vectorized upper triangle of a symmetric sFNC matrix, thus the dimensionality of each sample can be calculated as 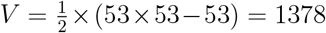. For dFNC data, we treat each windowed dFNC matrix as an independent sample, and denote the dFNC dataset as 𝒟_dFNC =_ x^(1,1)^, …, x^(*i,t*)^, …, x^(*N,T*)^. The number of total samples is the product of the number of subjects *N* and the number of windows *T* (*N × T*). Each sample x^(*i,t*)^ ∈ ℝ^*V*^is the flattened upper triangle of a windowed dFNC matrix from the *i*-th subject and the *t*-th window (*V* = 1378). For pedagogical purposes, we will explain the generative model using a single sample x ∈ ℝ^*V*^.

Let z ∈ ℝ^*D*^ denote a continuous latent variable which represents the underlying factors in the input data. We initially defined a two-dimensional latent space (i.e. *D* = 2) to better visualize the FNC matrices. Note that it is also possible to define a latent space with a different dimension. We assume that the latent variable z is sampled from a Gaussian prior distribution *p*(z) = 𝒩 (z|0, I) and the observed data variable x is sampled from the conditional likelihood distribution *p*_*θ*_(x|z) parameterized by *θ*, with the joint distribution *p*_*θ*_(x, z) = *p*_*θ*_(x|z)*p*(z). Our goal is to estimate the posterior distribution to perform inference on the latent variable. According to Bayes’ theorem, the posterior can be derived as:

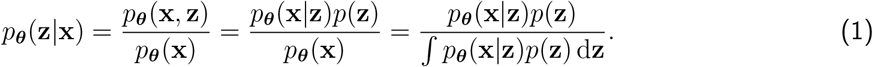

However, the computation of the posterior *p*_*θ*_(z|x) is analytically intractable because the integral of the _marginal likelihood ∫_*p*_*θ*_(x|z)*p*(z) dz cannot be evaluated in closed form. Hence, we perform *variational inference* to approximate the true posterior distribution with a simpler distribution.

Here, we approximate the posterior distribution using a multivariate Gaussian distribution with diagonal covariance, 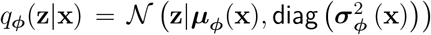. The Kullback-Leibler (KL) divergence is used to quantify the difference between the true posterior distribution *p*_*θ*_(z|x) parameterized by *θ* and the multivariate Gaussian approximation *q*_*ϕ*_(z|x) parameterized by *ϕ*. The objective function, also known as the *evidence lower bound (ELBO)*, aims to maximize the negative KL divergence:

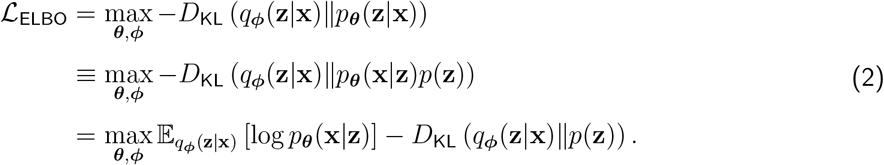

Note that the KL divergence is non-negative, i.e. *D*_KL_(·) ≥ 0. Thus, the ELBO is the lower bound to the log likelihood of generating the observed data, i.e. ℒ_ELBO_ ≤ log *p*_*θ*_(x). An encoder parameterized by the variational parameters *ϕ* (i.e. *q*_*ϕ*_(z|x)) and a decoder parameterized by the generative parameters *θ* (i.e. *p*_*θ*_(x|z)) are trained to maximize the ELBO, thereby simultaneously maximizing the log likelihood of generating the observed data samples and minimizing the KL divergence between the approximate posterior and the true posterior. Maximizing the ELBO is equivalent to minimizing the objective function as follows (see Appendix C for more details):

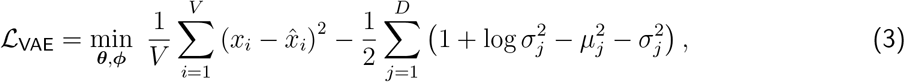

where *x*_*i*_ and 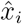 are the *i*-th element of the observed data x and the reconstructed data 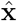, respectively; *µ*_*j*_ and *σ*_*j*_ are the *j*-th element of *µ*_*ϕ*_(x) and *σ*_*ϕ*_(x), respectively. To train the model using gradient-based optimization, the *reparameterization trick* (Kingma & Welling, 2014) is used: z = *µ*_*ϕ*_(x) + *σ*_*ϕ*_(x) ⊙ *ϵ*, where ⊙ represents the element-wise product, and *ϵ* ∈ ℝ^*D*^ is a noise variable *ϵ* ∼ 𝒩 (0, I). By optimizing the loss, the VAE is capable of learning the underlying factors in the dataset and generating continuous synthetic data by interpolating between latent samples using the learned factors.

A multilayer perceptron (MLP) was implemented as the encoder and decoder in the VAE. We performed a hyperparameter search to select the optimal model architecture that yielded the best training performance from four different architectures for each dataset and each data type separately (see Appendix F.1). The model was trained using the Adam optimizer (Kingma & Ba, 2017) with an initial learning rate of 0.001. We implemented a learning rate scheduler to reduce the learning rate by a factor of 0.1 if the loss value did not improve for 10 epochs. Each model was trained for 1000 epochs with a batch size of 16 and 512 for the sFNC and dFNC data, respectively. We implemented an early stopping mechanism – if the loss value did not decrease after 20 epochs, the training would stop. All training processes stopped within the predefined number of epochs. For each model, we initialized the model weights with 10 different random seeds and recorded the results across 10 runs. The model was implemented in Python using the PyTorch framework (Paszke et al., 2019) and trained with NVIDIA A40 GPUs.

### 2.3 Model comparison

To comprehensively evaluate model performance, we compared the VAE with a linear baseline method – probabilistic principal component analysis (PPCA) (Tipping & Bishop, 1999) and a semi-supervised alternative – identifiable variational autoencoder (iVAE) (Khemakhem et al., 2020). We review the generative model and optimization approach used in PPCA in Appendix D, and describe the generative model and objective function for iVAE in Appendix E.

#### 2.3.1 Probabilistic principal component analysis

PPCA is a probabilistic extension of PCA. Unlike conventional PCA, PPCA can estimate missing values in the dataset and generate samples from the learned distribution. Recent work has shown that the principal components learned by PPCA can be fully recovered by linear VAEs (Lucas et al., 2019). Thus, PPCA can be viewed as equivalent to linear VAEs. We implemented the expectation-maximization (EM) algorithm to estimate the model parameters, as described in Tipping and Bishop, 1999. We randomly initialized the model parameters with 10 seeds and reported the results across 10 runs.

#### 2.3.2 Identifiable variational autoencoders

Although a VAE is trained to learn latent distributions, there is no guarantee of identifiability^1^ of the model parameters in general. The iVAE has been developed to address the identifiability problem by conditioning on an auxiliary variable such as class label or time index. It has been shown that the iVAE recovers conditionally identifiable latent variables up to permutations and pointwise nuonlinear transformations (Khemakhem et al., 2020). Since our main interest lies in psychiatric characteristics, we used the one-hot encoded diagnostic label as the auxiliary variable in the iVAE. We performed a similar hyperparameter search experiment to select the optimal iVAE architecture (Appendix F.2). Each iVAE model was initialized using 10 different random seeds.

### 2.4 Functional network connectivity interpolation framework

We propose a functional network connectivity interpolation framework based upon the VAE (Figure 1). For each dataset, a VAE was trained on the sFNC data or the dFNC data separately. The vectorized upper triangle of each sFNC or windowed dFNC matrix was used as the input to the VAE. The VAE was trained to learn a latent variable z by minimizing the reconstruction loss between the input and the reconstructed output, while the KL divergence regularized the difference between the prior distribution *p*(z) and the approximate posterior distribution *q*_*ϕ*_(z|x). After the training stage, we performed sFNC interpolation and dFNC interpolation separately, as described below.

**Figure 1.**
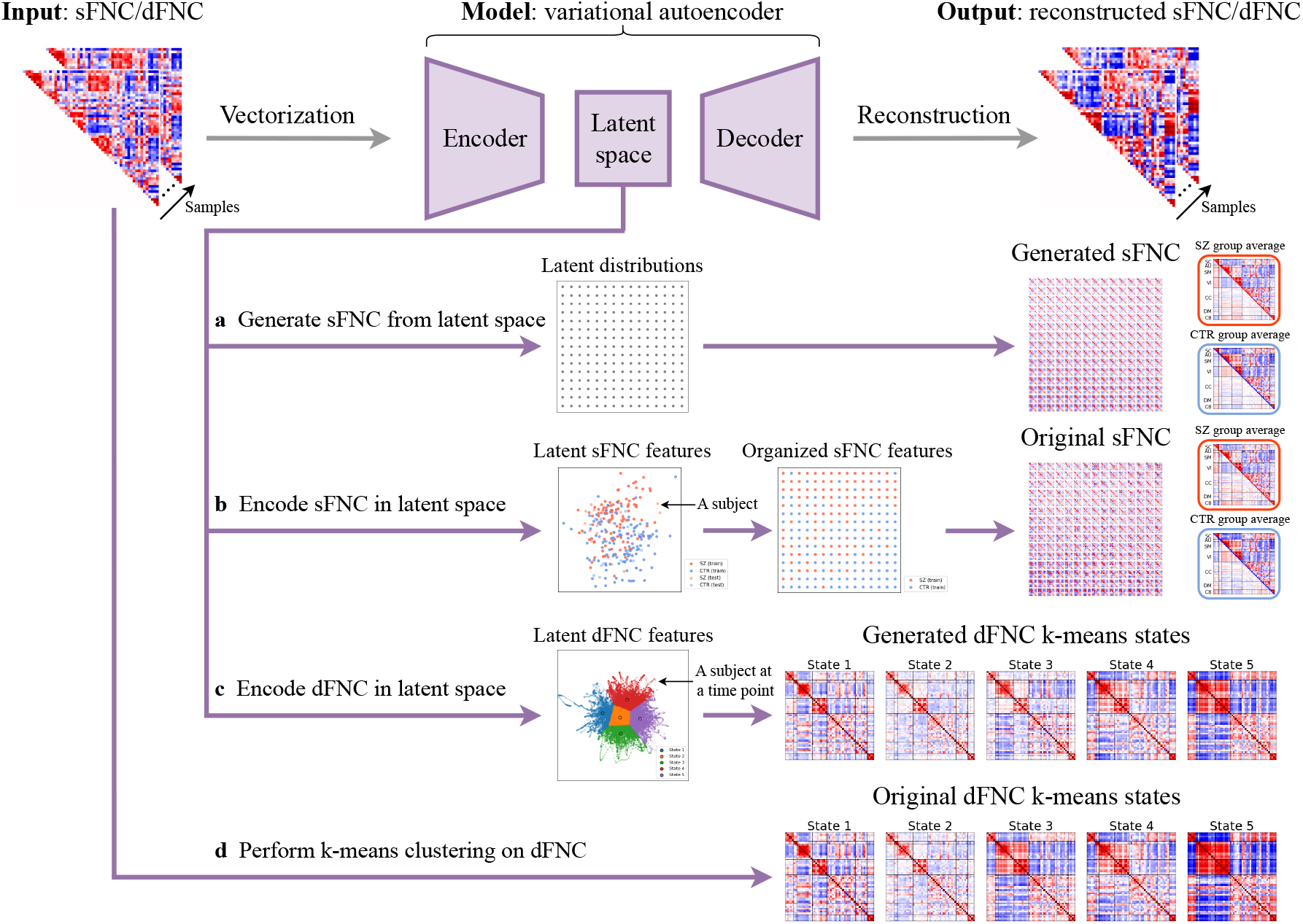
Overview of functional network connectivity interpolation framework. The upper triangle of each sFNC or windowed dFNC matrix was vectorized and used as the input to the VAE. During the training stage (gray arrows), a VAE was trained to learn a 2D latent variable z by minimizing the reconstruction loss between the input and the reconstructed output, while the KL-divergence regularized the difference between the prior distribution *p*(z) and the approximate posterior distribution *q*_*ϕ*_(z|x). During inference time, four pathways (purple arrows) are described as follows: (a) The sFNC matrices were generated by uniformly sampling from the prior distribution *p*(z) in the latent space and then forward propagating through the trained decoder. (b) The original sFNC matrices were projected onto the latent space by the trained encoder, the corresponding latent features were then mapped to a 2D grid using the JV algorithm, and the corresponding original sFNC matrices were then displayed on the 2D grid. (c) The dFNC matrices were projected onto the latent space by the trained encoder. The *k*-means clustering algorithm was then performed on the dFNC latent features. The dFNC matrices were generated by sampling from the posterior distributions *q*_*ϕ*_(z|x). The generated dFNC state was then computed as the element-wise median of the generated dFNC matrices in each cluster. (d) The *k*-means clustering algorithm was performed on the dFNC matrices directly. The original dFNC state was then computed as the element-wise median of the original dFNC matrices in each cluster.

#### 2.4.1 Static functional network connectivity interpolation

We first fed the original training or test data to the trained encoder and extracted the corresponding two-dimensional features in the latent space. To generate synthetic data, we defined a 15 *×* 15 grid of evenly spaced 2D coordinates in the latent space (as shown in Figure 1a), with the range determined by the 80th percentile of the absolute values of latent features from the training set. To display all training subjects’ sFNC matrices on a 2D grid, we select the grid dimension such that the total number of grid points (i.e., the square of the dimension) approximates the number of training samples. These coordinates (values of latent variables) were then forward propagated through the trained decoder to generate continuous synthetic sFNC matrices, which were visualized on the 2D grid. To examine the corresponding original data, the Jonker-Volgenant (JV) algorithm (Jonker & Volgenant, 2005) for the linear sum assignment problem was subsequently used to map these latent features to the nodes on the 2D grid by minimizing the pairwise Euclidean distance between the positions of the latent features and the 2D grid nodes. We then displayed the original sFNC matrices on the 2D grid correspondingly. This allows us to visualize the generated sFNC matrices and the original sFNC matrices on the 2D grid, respectively.

#### 2.4.2 Dynamic functional network connectivity interpolation

We projected the windowed dFNC matrices onto the VAE latent space and extracted their corresponding latent features. Subsequently, the *k*-means clustering algorithm was applied to these dFNC latent features, using a fixed random seed and varying the number of states *k* from 2 to 9. The dFNC matrices were generated by sampling from the learned posterior distributions *q*_*ϕ*_(z|x). The group-specific generated dFNC state was then computed as the element-wise median of the generated dFNC matrices in each cluster for each group. For comparison, the *k*-means clustering algorithm was performed on the original dFNC matrices directly with the same search window of *k*, and then the optimal *k* value was identified using the same elbow criterion. The group-specific original dFNC state was then computed as the element-wise median of the original dFNC matrices in each cluster for each group.

To further quantify the properties of the dFNC states, we calculated the group-specific dwell time (DT) for each state *α*, defined as the number of FNC matrices per state 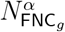 divided by the number of subjects per state 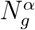 for each group *g*:

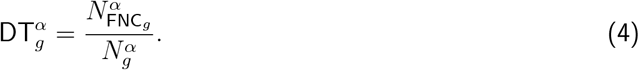

We also calculated the group-specific transition probability matrix P_*g*_ ∈ ℝ^*k×k*^to evaluate how likely a state changes to another between two consecutive time points. Specifically, we first counted the number of transitions from a state *α* to a different state *β* through the entire time course *T* for each group *g*:

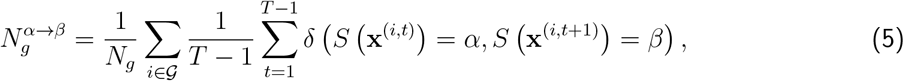

where *δ*(·) = 1 when the transition happens from the state *α* at time *t* to the state *β* at time *t* + 1 for the *i*-th subject, otherwise *δ*(·) = 0. *S*(·) ∈ {1, …, *k*} indicates the state of a sample. 𝒢 is the set of subject indices for a group and *N*_*g*_ is the corresponding number of subjects for that group.

Then, the off-diagonal entry of the group-specific transition probability matrix P_*g*_(*α, β*) (*α*≠ *β*) is defined as the number of transition times from the state *α* to the state *β* divided by the total number of transition times from the state *α* to all the other states:

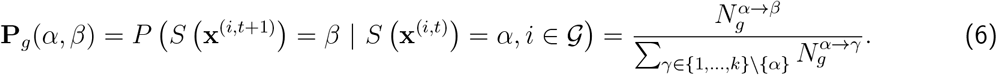

## 3 Results

### 3.1 sFNC interpolation captures psychosis continuum and heterogeneity

We first compared VAE performance with two alternative methods, probabilistic principal component analysis (PPCA) and identifiable variational autoencoders (iVAEs). We hypothesized that nonlinear VAEs that capture more complex relationships from observed data would achieve better performance than linear PPCA. As expected, the Pearson correlations between the generated and original sFNC matrices from VAEs are significantly higher than those from PPCA (Figure 2, *p <* 0.0001 for both FBIRN and ABIDE I, Wilcoxon signed-rank test, Bonferroni correction). More specifically, the average correlations for VAEs are 0.765 and 0.724, while those for PPCA are 0.711 and 0.694 for FBIRN and ABIDE I data, respectively. Surprisingly, VAEs also significantly outperformed iVAEs in this interpolation task (Figure 2, *p <* 0.05 for FBIRN, *p <* 0.01 for ABIDE I, Wilcoxon signed-rank test, Bonferroni correction). The average correlations for iVAEs are 0.741 for the FBIRN dataset and 0.708 for the ABIDE I dataset, both slightly lower than those for VAEs. By design, we consider VAEs better suited for learning low-dimensional continuous representations in this case due to their *nonlinear* and *unsupervised* nature. Our empirical results support this theoretical advantage, demonstrating that VAEs significantly outperformed both a linear baseline (PPCA) and a semi-supervised alternative (iVAE). Therefore, we focused on VAEs for all subsequent experiments in our study.

**Figure 2.**
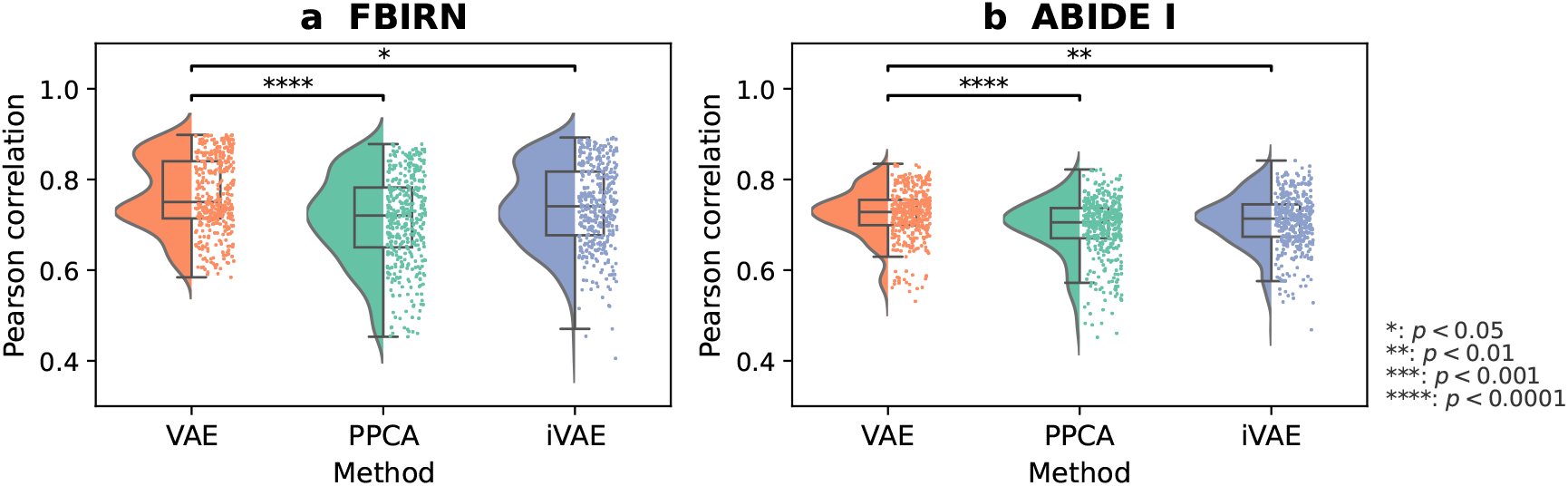
Pearson correlations between generated and original sFNC matrices. (a) FBIRN test data. (b) ABIDE I test data. The scatter-box-violin plot shows the data distribution, median and interquartile range of Pearson correlations between the original and generated sFNC matrices in the FBIRN and ABIDE I test sets for VAE, PPCA, and iVAE, across 10 random seeds. In both datasets, VAEs significantly outperformed iVAEs and PPCA in the interpolation task (Wilcoxon signed-rank test, Bonferroni correction).

We confirmed that the VAE representations were robust to potential confounding factors, including age, sex, and collection site (Appendix G), and subsequently performed sFNC interpolation as described in Section 2.4.1. By comparing the generated sFNC matrices and the original sFNC matrices, we observe a high degree of correspondence between the generated and original results for both datasets (Figures 3, 4; zoom-in version in Appendix H). Group-specific patterns, group pattern alterations, and individual differences can be visualized on the 2D grid layout. By examining the group-specific patterns in both the generated and original sFNC matrices, we observe focal modularity in the top half dominated by the patient group, and highly modular and polar patterns in the bottom half dominated by the control group. Specifically, the SZ patient group shows weak functional connectivity among the auditory (AU), sensorimotor (SM), and visual (VI) domains (Figure 3). Similarly, the ASD patient group also shows weak connectivity in these sensory networks (Figure 4). In contrast, the control group shows strong functional connection within the sensory domains (AU, SM, VI), as well as between the subcortical (SC) and cerebellar (CB) domains. The average cell-wise standard deviations of the patient group is slightly higher than that of the control group, suggesting slightly higher variability in the patient group (Appendix I). In addition, the generated sFNC matrices reveal continuous pattern alterations between two groups. The continuous sFNC spectrum from the SZ patient group to the control group demonstrates two key connectivity patterns: (1) stronger positive correlations within the sensory domains (AU, SM, VI) and between the SC and CB domains, and (2) stronger negative correlations between the SC domain and the sensory networks, as well as between the CB domain and these sensory regions. We observe a similar continuum of sFNC changes from ASD patients to controls, but the group differences are less pronounced compared to SZ, suggesting that functional connectivity alterations in ASD may not be as substantial as those observed in SZ. Overall, the generated sFNC matrices show representative patterns within a group and continuous alterations between groups, while the original sFNC matrices help to further examine individual variability.

**Figure 3.**
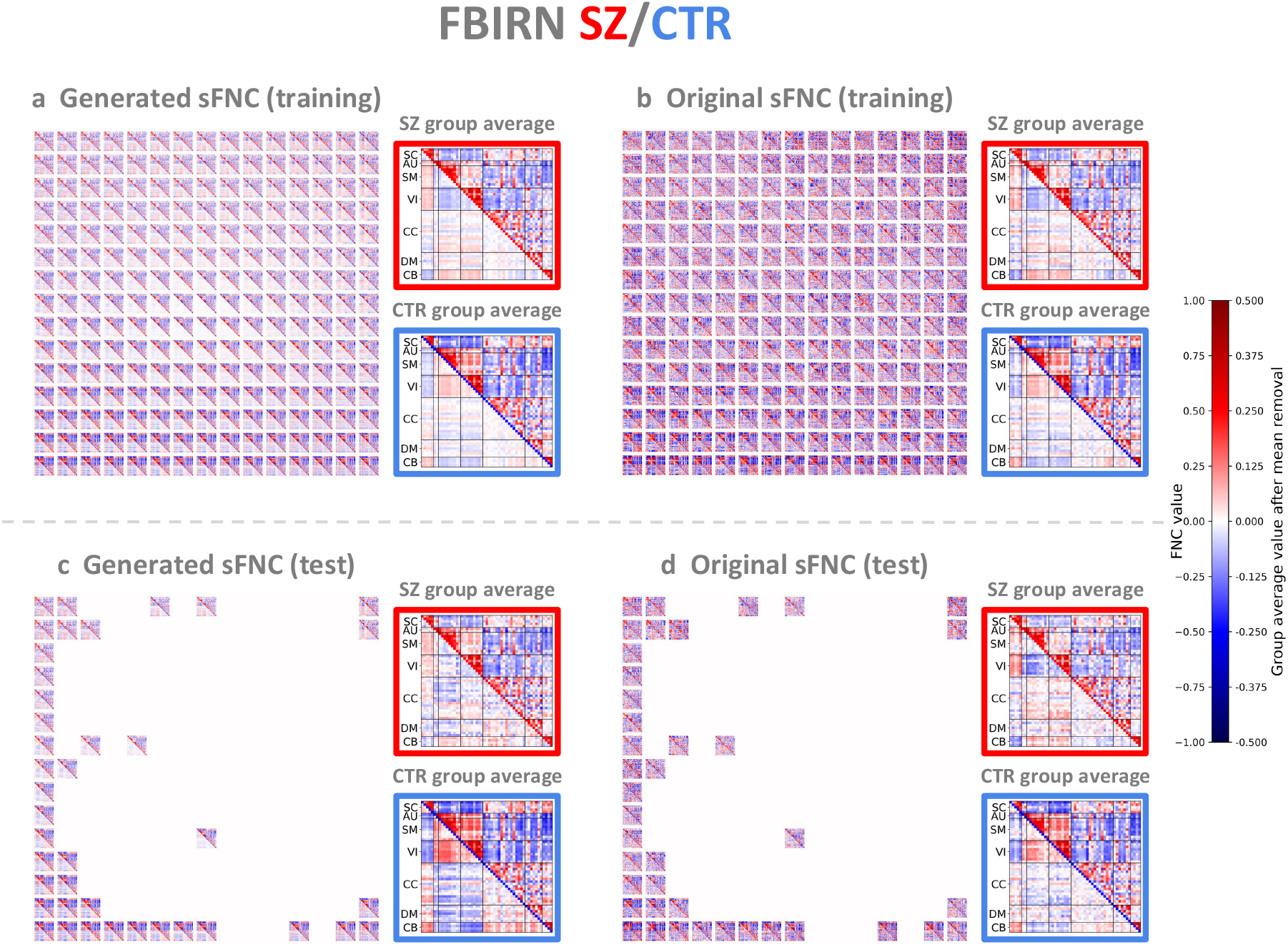
Generated and original FBIRN sFNC matrices. (a) Generated sFNC matrices in the training set. (b) Original sFNC matrices in the training set. (c) Generated sFNC matrices in the test set. (d) Original sFNC matrices in the test set. For each sFNC matrix, we retained the upper triangle, removed the element-wise mean across all subjects in the lower triangle to investigate how each subject deviates from the population mean, and color-coded the diagonal based on the diagnosis groups (red: patients; blue: controls). In each red or blue box, the SZ patient or control group average pattern is shown in the upper triangle. The corresponding pattern after removing the element-wise mean across all subjects is shown in the lower triangle (values were multiplied by 2 for better visualization). There is a high degree of correspondence between generated and original sFNC matrices. The generated sFNC matrices capture a continuous spectrum of pattern alterations between groups, while the original sFNC matrices show individual variability across subjects.

**Figure 4.**
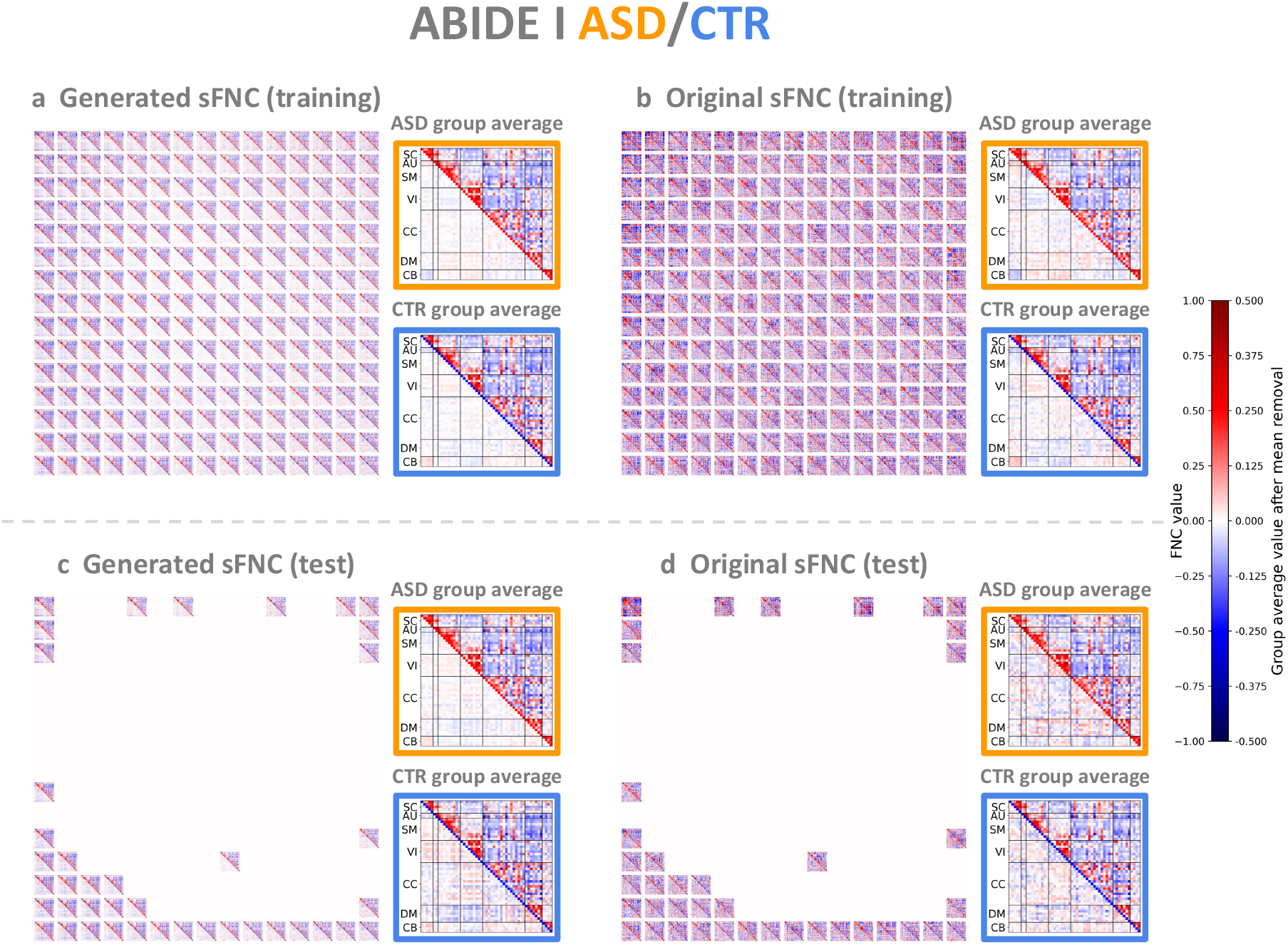
Generated and original ABIDE I sFNC matrices. (a) Generated sFNC matrices in the training set. (b) Original sFNC matrices in the training set. (c) Generated sFNC matrices in the test set. (d) Original sFNC matrices in the test set. In each orange or blue box, the ASD or control group average pattern is shown in the upper triangle and the respective group average pattern after removing the element-wise mean across all subjects is shown in the lower triangle. The group average values after mean removal (the lower triangles) in the colored boxes were multiplied by 2 for better visualization.

Furthermore, we displayed multiple FBIRN subject measures, including clinical assessment scores, normalized cognitive scores (van Erp et al., 2015), age and gender information, on the 2D grid accordingly (Figure 5), providing interpretable visualization of how each subject measure contributes to the unsupervised learning process. Crucially, we observe that diagnosis is a key factor distinguishing subjects in the latent space, with most patients occupying the upper half of the 2D grid and most controls occupying the lower half. In addition, age appears to be another important factor, as younger subjects tend to cluster in the lower triangle, while older subjects are mostly in the upper triangle. Interestingly, SZ patients located in the lower/left/lower triangular half of the 2D grid have relatively higher (better) cognitive scores than those located in the upper/right/upper triangular half (Table 2). In particular, the speed of processing, reasoning/problem solving, and composite scores in the lower half are significantly higher than those in the upper half, and in addition, composite scores in the lower triangle are significantly higher than those in the upper triangle (*p <* 0.05, Mann–Whitney U test), implying that our framework can group patients in the latent space based on their cognitive ability and further identify subgroups in SZ. To support this application, we applied *k*-means clustering to the latent representations to identify subgroups, examined their associations with subject measures, and characterized subgroup-specific connectivity patterns (Appendix J). To examine how the dimensionality of the VAE latent space affects downstream subgroup identification, we compared the relationships between *k*-means subgroups and cognitive scores in both 2D and three-dimensional (3D) latent space, and found that the 3D latent space does not yield additional meaningful subgroups (Appendix J). Moreover, we investigated sFNC patterns across cognitive performance levels and identified both common and distinct connectivity changes across cognitive measures (Appendix K). ABIDE I subject measures (clinical assessment scores, age, and gender) are shown in Figure A.1, Appendix A.

**Table 2:** Cognitive measure statistics of the FBIRN patient cohort. We partitioned the 2D grid into six regions: the upper 7 rows (upper), the lower 8 rows (lower); the left 7 columns (left), the right 8 columns (right); the upper triangle with the diagonal (upper triangle), and the lower triangle without the diagonal (lower triangle). In the Table, the first row shows the number of SZ patients in each region. The second to eighth rows show the mean (standard deviation) of each cognitive measure for SZ patients across different regions. A higher (better) cognitive score is highlighted in bold for each row. The star (∗) indicates that cognitive scores in one half are significantly higher than those in the other half (*p <* 0.05, Mann‐Whitney U test). Overall, the SZ patients located in the lower/left/lower triangular half of the 2D grid have consistently *higher* cognitive scores compared to those located in the upper/right/upper triangular half, implying that our framework can group SZ patients in the latent space based on their cognitive performance.

| Cognitive measure | Upper | Lower | Left | Right | Upper triangle | Lower triangle |
| --- | --- | --- | --- | --- | --- | --- |
| Diagnosis | 69 | 36 | 57 | 48 | 67 | 38 |
| Speed of processing | −1.35(1.0) | **−0.92(0.8)\*** | **−1.13(0.9)** | −1.29(1.0) | −1.34(1.0) | **−0.96(0.8)** |
| Attention/Vigilance | −1.41(1.4) | **−1.05(1.2)** | **−1.06(1.3)** | −1.55(1.4) | −1.43(1.4) | **−1.05(1.2)** |
| Working memory | −1.10(1.0) | **−0.79(0.9)** | **−0.91(0.9)** | −1.10(1.0) | −1.14(1.0) | **−0.74(0.9)** |
| Verbal learning | −1.38(1.2) | **−1.11(0.9)** | **−1.22(1.0)** | −1.37(1.2) | −1.40(1.2) | **−1.10(1.0)** |
| Visual learning | −1.09(1.1) | **−0.81(1.1)** | **−0.92(1.0)** | −1.08(1.2) | −1.11(1.1) | **−0.78(1.0)** |
| Reasoning | −0.87(1.1) | **−0.40(0.9)\*** | **−0.66(1.1)** | −0.76(1.0) | −0.83(1.1) | **−0.49(0.9)** |
| Composite score | −1.70(1.2) | **−1.21(1.0)\*** | **−1.39(1.1)** | −1.71(1.3) | −1.71(1.3) | **−1.22(0.9)\*** |

**Figure 5.**
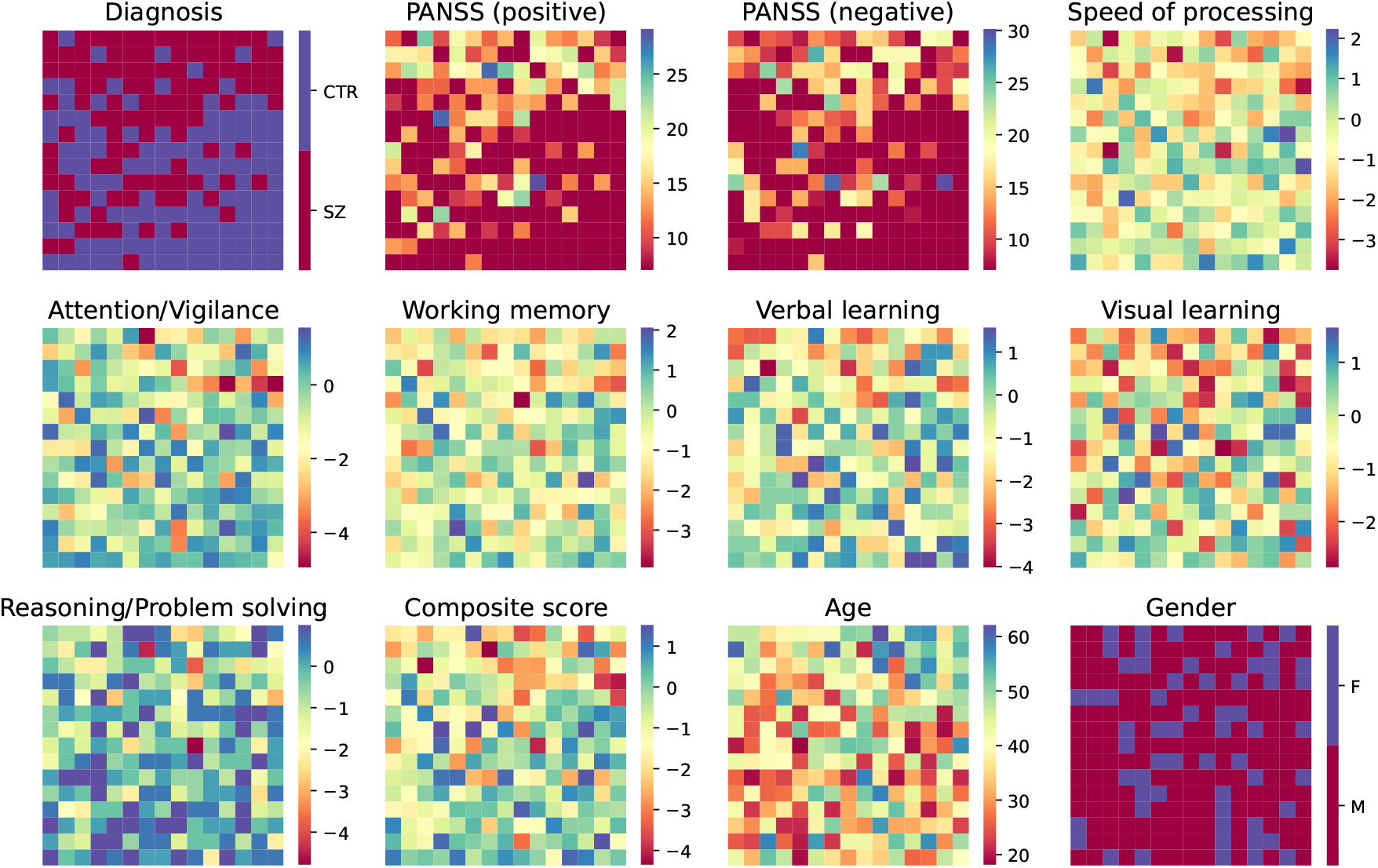
FBIRN subject measures displayed on the 2D grid. Based on the individualized mapping between latent features and 2D grid nodes (as in Figure 3), we displayed diagnosis labels, PANSS positive and negative subscale scores, CMINDS cognitive scores (including speed of processing, attention/vigilance, working memory, verbal learning, visual learning, reasoning/problem solving, and composite scores), age and gender information of FBIRN subjects on the 2D grid correspondingly.

In addition, we measured Pearson correlations between the generated and original sFNC matrices to quantify the correspondence (Figure 6). For FBIRN, all 225 matrices (100.0%) used for training have correlations greater than 0.6 and 106 out of 225 matrices (47.1%) have correlations greater than 0.8, with the median correlations of 0.819 and 0.768 for controls and SZ patients, respectively (Figure 6a). For ABIDE I, 224 out of 225 matrices (99.6%) have correlations greater than 0.6 and 54 out of 225 matrices (24.0%) have correlations greater than 0.8, with the median correlations of 0.784 and 0.762 for the control and ASD patient groups, respectively (Figure 6b). We also observe high correlations from the unseen test sets (Figure 6c,d). In the FBIRN test set, the median correlations for the control group and the patient group are 0.792 and 0.747, respectively, while in the ABIDE I test set, they are 0.740 and 0.711, respectively. Mean squared errors (MSE) between the generated and original sFNC matrices also demonstrate their similarity, with the median MSEs of 0.031, 0.034, 0.029, and 0.034 for the FBIRN training set, FBIRN test set, ABIDE I training set, and ABIDE I test set, respectively (Appendix L). These comparisons indicate that the generated sFNC matrices well aligned with the original ones, especially for controls, who show more homogeneity in their functional connectivity. Thus, the VAE latent space can be subsequently used to generate continuous samples by interpolating along pathological trajectories.

**Figure 6.**
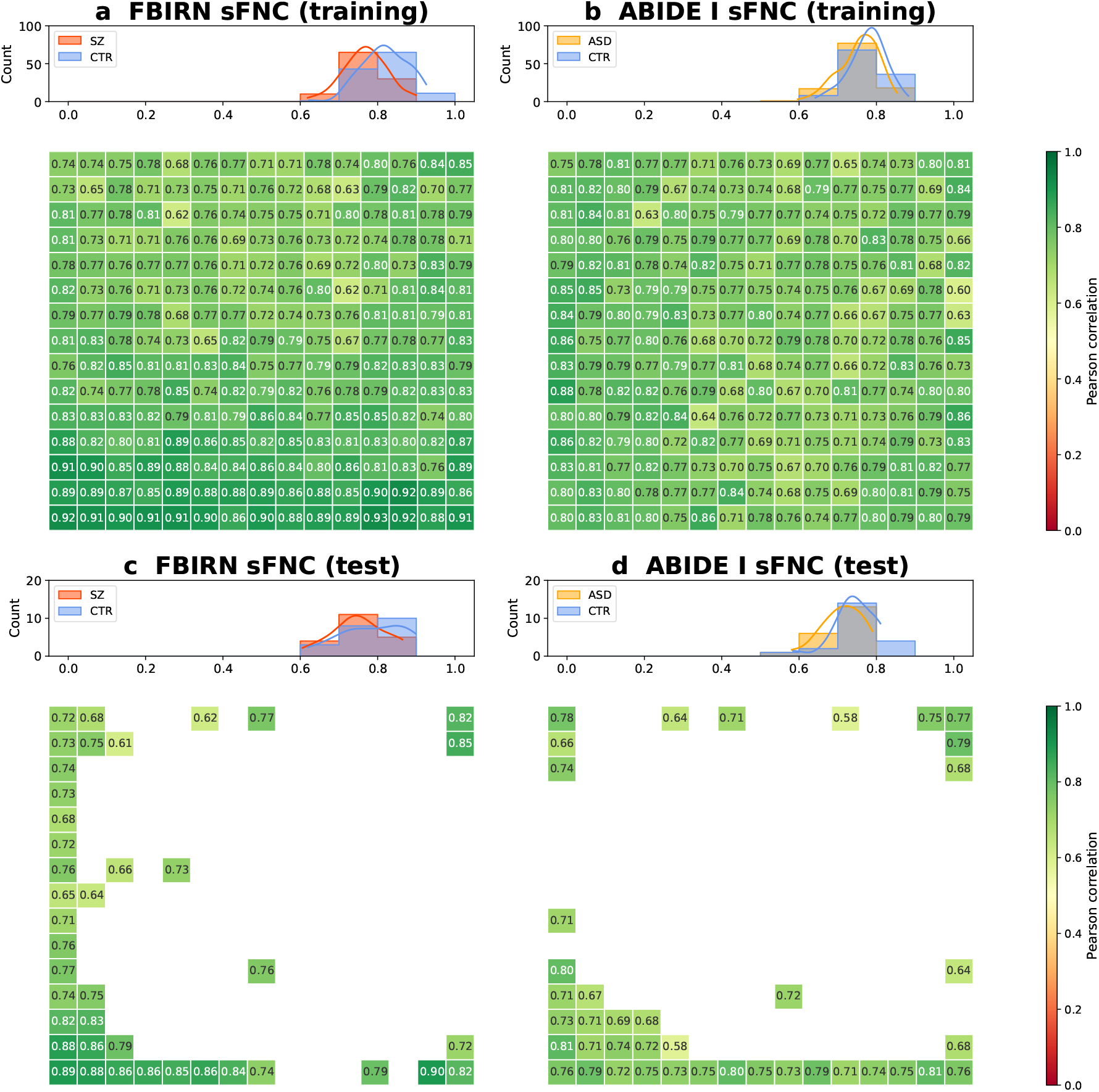
Pearson correlations between generated and original sFNC matrices. (a) FBIRN training set. (b) ABIDE I training set. (c) FBIRN test set. (d) ABIDE I test set. Each panel shows Pearson correlations between individual generated and original sFNC matrices on the 2D grid and the corresponding group-specific histogram distributions. High correlations across all panels suggest that the generated sFNC matrices well aligned with the original sFNC matrices ordered by the JV algorithm, especially for controls who show less variability in functional connectivity.

### 3.2 dFNC interpolation reveals group-specific dynamic states

To further evaluate how these group-specific patterns change over time, we performed a dynamic state analysis using the *k*-means clustering algorithm on the dFNC data (Section 2.4.2). We searched *k* from 2 to 9, and considered that five clusters were optimal for both FBIRN and ABIDE I datasets based upon the elbow criterion on the *L*_2_ distance (Appendix M). The group-specific dFNC states and group differences for FBIRN and ABIDE I datasets are presented in Figures 7 and 8, respectively.

**Figure 7.**
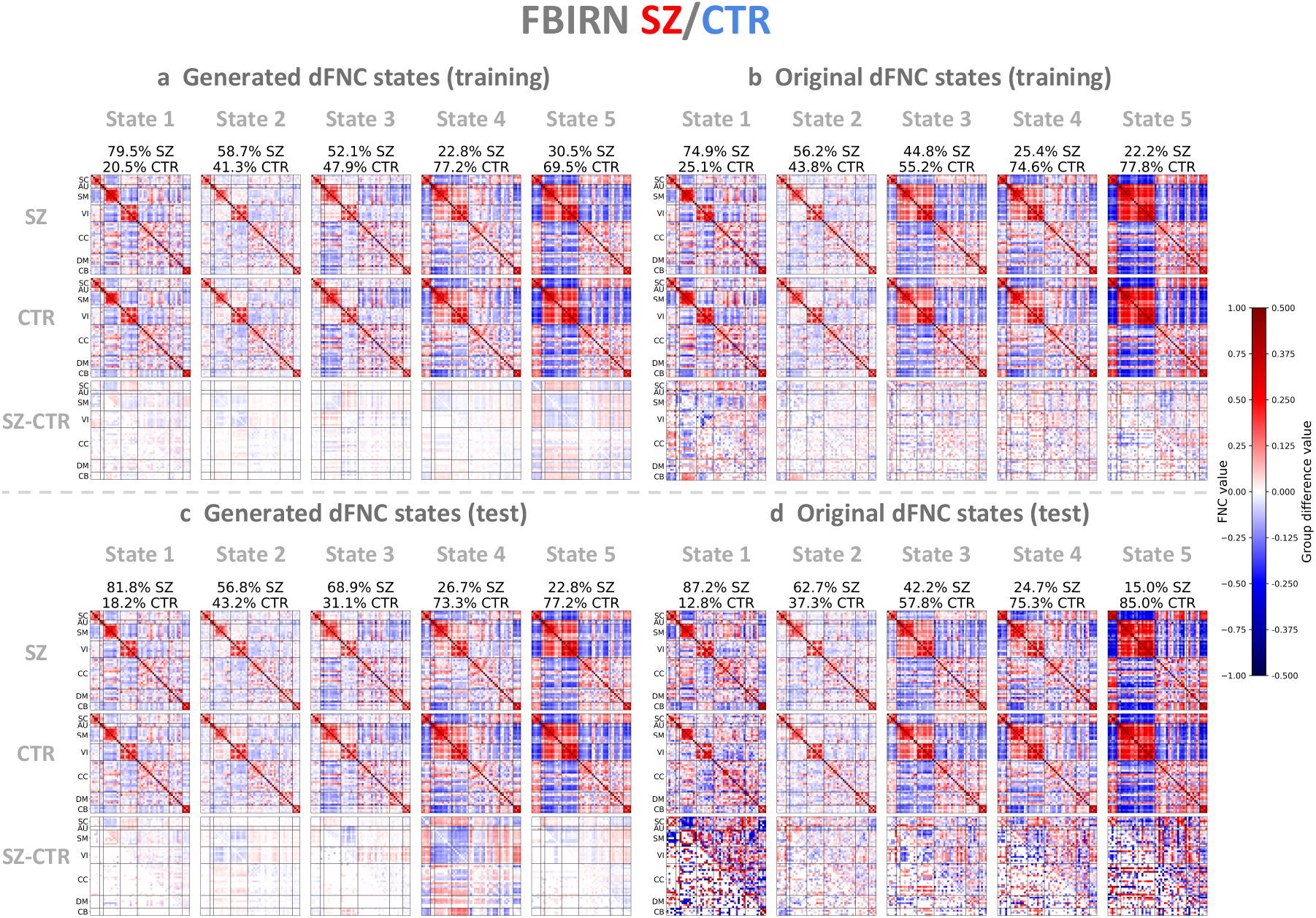
Generated and original FBIRN dFNC states. (a) Five generated dFNC states of SZ patients, controls and group differences in the training set. (b) Five original dFNC states in the training set. (c) Five generated dFNC states in the test set. (d) Five original dFNC states in the test set. The generated and original dFNC states represent the element-wise medians of dFNC patterns, obtained by applying *k*-means clustering to the VAE latent features and the dFNC matrices, respectively. The original dFNC states were sorted by the increasing percentage of control windows relative to total windows per state, and the generated states were ordered based on their similarity to the original states. Group differences between patient and control states are shown in the upper triangle, and significant differences are shown in the lower triangle (*p <* 0.05, two-sample *t*-test, Bonferroni correction), with all values multiplied by 2 for better visualization. The generated dFNC states revealed state-based group differences, similar to the original ones. Moreover, the VAE effectively captured group-specific representations that are generalizable to the unseen test set.

**Figure 8:**
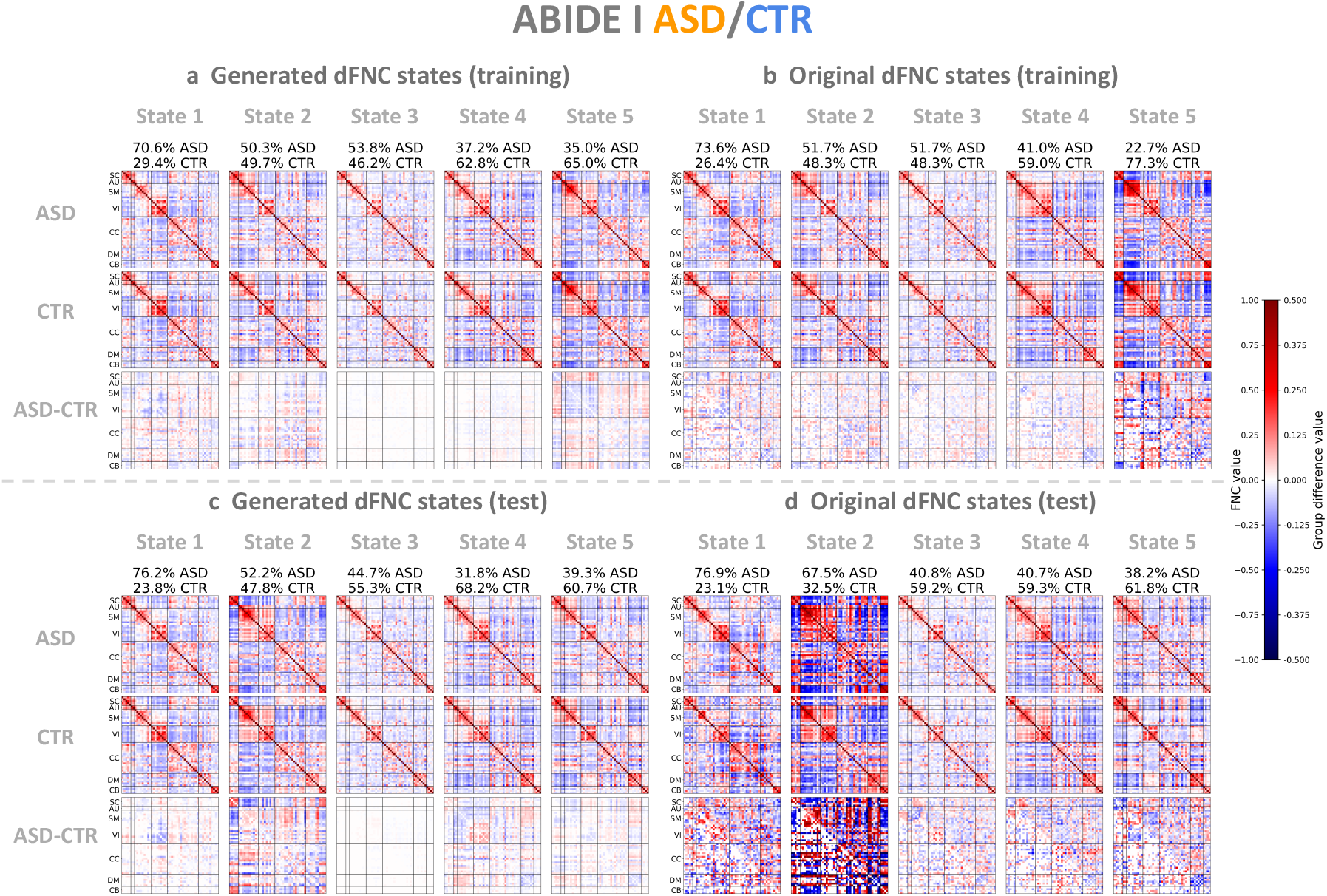
Generated and original ABIDE I dFNC states. (a) Five generated dFNC states of ASD patients, controls and group differences in the training set. (b) Five original dFNC states in the training set. (c) Five generated dFNC states in the test set. (d) Five original dFNC states in the test set. The generated and original dFNC states represent the element-wise medians of dFNC patterns, obtained by applying *k*-means clustering to the VAE latent features and the dFNC matrices, respectively. The original dFNC states were sorted by the increasing percentage of control windows relative to total windows per state, and the generated states were ordered based on their similarity to the original states. Group differences between patient and control states are shown in the upper triangle, and significant differences are shown in the lower triangle (*p <* 0.05, two-sample *t*-test, Bonferroni correction), with all values multiplied by 2 for better visualization.

Similar to the sFNC results, we demonstrate high correlations between the generated dFNC states and the original ones. Specifically, for the FBIRN training set, we first applied *k*-means clustering to the dFNC latent features from all subjects (including SZ patients and controls) to derive five shared states. We then computed the Pearson correlations between the generated and original dFNC states (1 to 5) separately for each group. The resulting correlations are 0.990, 0.990, 0.882, 0.983, 0.967 for the SZ patient group, and 0.958, 0.982, 0.872, 0.994, 0.974 for the control group, respectively (Figure 7a,b). For the FBIRN test set, the correlations from states 1 to 5 are 0.902, 0.968, 0.891, 0.935, 0.908 for the patient group and 0.820, 0.955, 0.882, 0.967, 0.955 for the control group, respectively (Figure 7c,d). In state 1, SZ patients, compared with controls, exhibit stronger negative correlations between the subcortical (SC) and sensorimotor (SM) domains and between the SM and visual (VI) domains. Conversely, they show increased positive correlations between the SC and VI domains and between the SM and cerebellar (CB) domains. In state 5, SZ patients show stronger negative correlations between the SM and VI domains and between the SC and CB domains, and stronger positive correlations between the SC domain and both the SM and VI domains, and between the CB domain and these two sensory areas. When evaluated on the unseen test set, the original group difference patterns appear noisy due to the small sample size (Figure 7d). Nonetheless, the generated group difference patterns (Figure 7c) closely resemble those in the training set, particularly between test state 2 and training state 3 (Pearson correlation coefficient *r* = 0.811), and test state 4 and training state 4 (*r* = 0.793). This finding suggests that the VAE can effectively capture *generalizable* group-specific representations.

For the ABIDE I training set, the correlations between the generated and original dFNC states 1 to 5 are 0.985, 0.994, 0.995, 0.982, 0.924 for the ASD patient group, and 0.971, 0.990, 0.993, 0.984, 0.947 for the control group, respectively (Figure 8a,b). For the ABIDE I test set, the correlations from states 1 to 5 are 0.957, 0.820, 0.974, 0.947, 0.961 for the patient group, and 0.885, 0.867, 0.980, 0.969, 0.960 for the control group, respectively (Figure 8c,d). In state 1 which is mainly occupied by ASD patients, we observe significant group differences, including negative correlations between the SM and VI domains and between the SC and SM domains, as well as positive correlations between the VI domain and the SC, CB and DM domains. These patterns partially overlap with those in the FBIRN state 1, indicating that ASD and SZ may share similar neurobiological deficits.

Subsequently, we evaluated two group-specific state-based dynamic measures – dwell time and transition probability matrix – to further quantitatively compare the generated and original dFNC states (Figure 9). The group-specific dwell time measures the average amount of time that a group lingers in a state. The dwell time statistics of the generated states are largely consistent with those of the original states in both datasets. For the FBIRN training set (Figure 9a,b), states 1, 2, 4 and 5 show significant group differences in both the generated and original states (*p <* 0.01, two-sample *t*-test, FDR correction). In particular, SZ patients stay in weakly connected states 1 and 2 significantly longer, while controls stay in densely connected states 4 and 5 significantly longer. For the FBIRN test set (Figure 9c,d), we note that the generated results show significant group differences in state 5 (*p <* 0.05, two-sample *t*-test, FDR correction), whereas the original results do not show any significant differences across five states.

**Figure 9.**
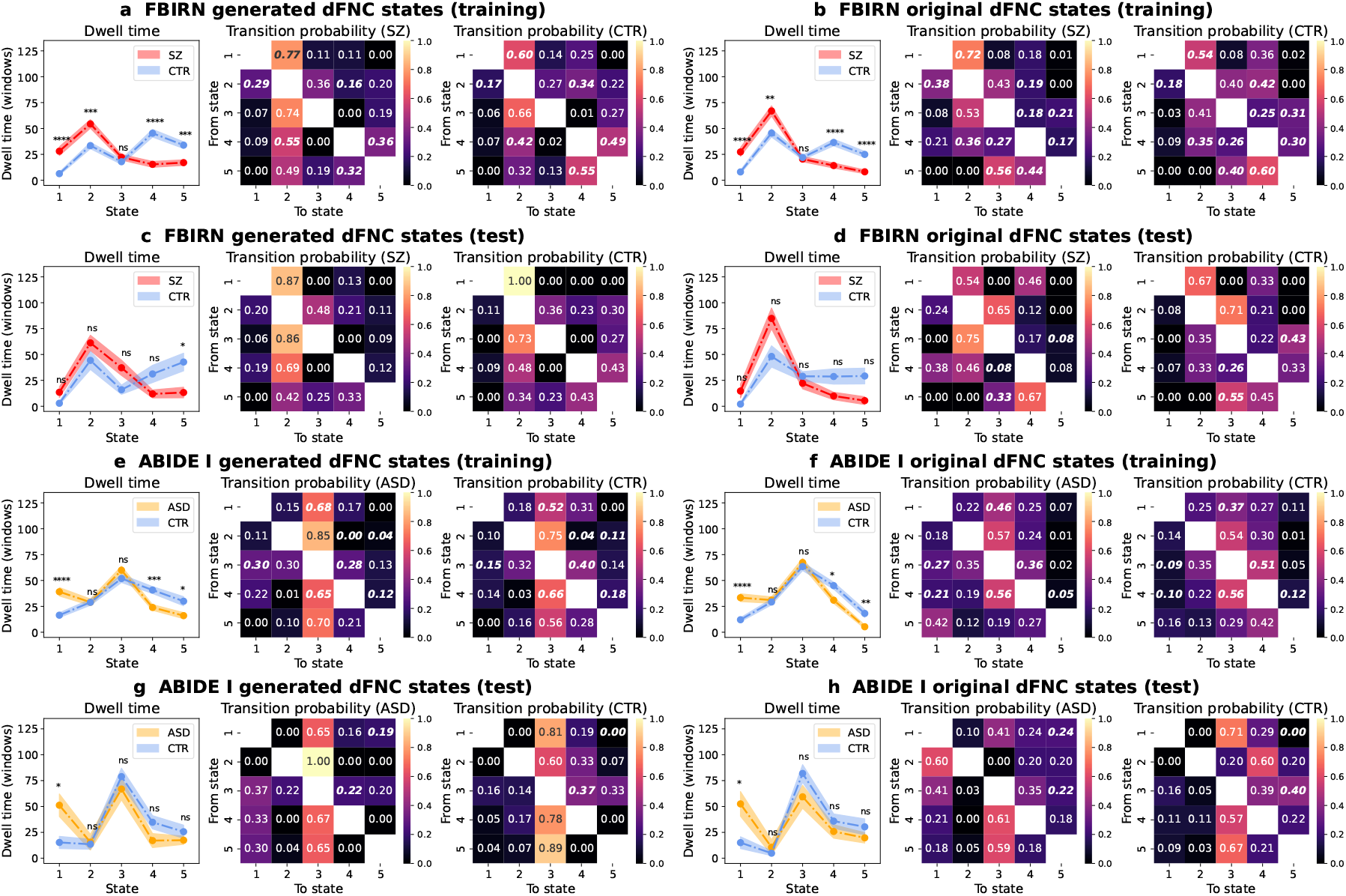
Group-specific dwell time and transition probability matrices of generated and original dFNC states. Parts a,b,c,d show the results from the FBIRN training and test sets, while parts e,f,g,h show the results from the ABIDE I training and test sets. Parts a,c,e,g show the metrics derived from the generated data and parts b,d,f,h present the results from the original data. In each part, the left panel shows the mean *±* standard error of dwell time measured by windows for different groups (red: SZ patients; orange: ASD patients; blue: controls). Two-sample *t*-tests with the false discovery rate (FDR) correction for five states were used to assess group differences (state labels are consistent with Figures 7 and 8). The number of asterisks (1, 2, 3, 4) indicates the level of significance (*p <* 0.05, *p <* 0.01, *p <* 0.001, *p <* 0.0001, FDR correction). The abbreviation “ns” stands for “not significant”. The middle and right panels show the transition probabilities between every two states for patients and controls, respectively. The transition probability is calculated by counting the number of transition times from one state to another state divided by the number of subjects from each group, and then normalizing the sum of each row to 1. Significant transition count differences between patients and controls are highlighted in bold italics (*p <* 0.05, two-sample *t*-test). The generated states exhibited group-specific properties, such as dwell time and transition probabilities, that closely matched the original states. These metrics indicate that VAEs effectively captured state-based dynamic properties.

For the ABIDE I training set (Figure 9e,f), the dwell time for the ASD group is significantly longer than that for the control group in both the generated and original state 1 (*p <* 0.0001, two-sample *t*-test, FDR correction). In contrast, the dwell time for the control group is significantly longer than that for the ASD group in both the generated and original states 4 and 5 (*p <* 0.05, two-sample *t*-test, FDR correction). No significant group differences are observed in states 2 and 3. For the ABIDE I test set (Figure 9g,h), the dwell time statistics in the generated results are also consistent with the original ones. Only the patient-dominant state 1 shows significant group differences (*p <* 0.05, two-sample *t*-test, FDR correction).

The group-specific transition probability matrix shows the probability that a group switches from one state to another. For the FBIRN training set (Figure 9a,b), we note that SZ patients are more likely to transition between loosely connected states 1 and 2 (state 1 to state 2: *P*_generated_ = 0.77, *P*_original_ = 0.72; state 2 to state 1: *P*_generated_ = 0.29, *P*_original_ = 0.38). In contrast, controls show significantly lower transition probabilities when moving between state 1 and state 2 (state 1 to state 2: *P*_generated_ = 0.60, *P*_original_ = 0.54; state 2 to state 1: *P*_generated_ = 0.17, *P*_original_ = 0.18). By comparison, controls are more likely to switch between densely connected states 4 and 5 (state 4 to state 5: *P*_generated_ = 0.49, *P*_original_ = 0.30; state 5 to state 4: *P*_generated_ = 0.55, *P*_original_ = 0.60), whereas patients with SZ have relatively lower probabilities of switching between states 4 and 5 (state 4 to state 5: *P*_generated_ = 0.36, *P*_original_ = 0.17; state 5 to state 4: *P*_generated_ = 0.32, *P*_original_ = 0.44).

Interestingly, for the ABIDE I training and test data (Figure 9e-h), both the ASD patient and control groups tend to switch from other states to state 3, suggesting that it may act as a hub for dynamic state transitions. In general, controls exhibit significantly higher transition probabilities from other states to state 4, while autistic patients are more likely to transition to state 1.

Taken together, the generated states show highly similar group-specific properties, such as dwell time and transition probabilities, as the original ones. These results indicate that VAEs effectively learn state-based dynamic properties without explicit incorporation of time information during the training stage.

### 3.3 A low-dimensional manifold to interpret and interpolate FNC

In our proposed interpolation framework, the VAE latent space serves as a low-dimensional manifold to interpret and interpolate sFNC or dFNC matrices, providing novel insights into how FNC patterns change along a trajectory of interest. Here, we show two potentially useful applications of the interpolation framework: (1) sFNC interpolation between patient and control groups to estimate a psychosis continuum (Figure 10) and (2) dFNC interpolation between two states to investigate state transition (Figure 11).

**Figure 10.**
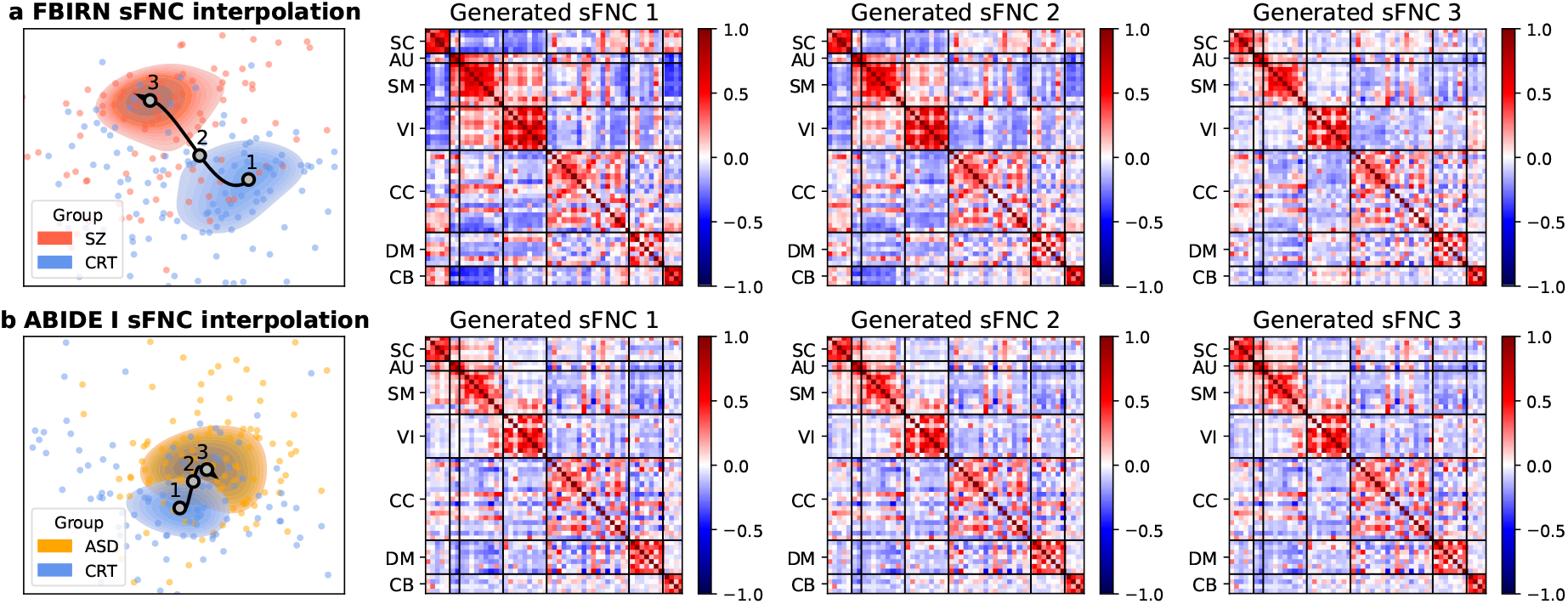
sFNC interpolation between diagnostic groups in the VAE latent space. (a) FBIRN training data. (b) ABIDE I training data. In each row, the scatter plot shows latent features colored by the diagnostic label (red: SZ patients; orange: ASD patients; blue: controls), overlaid with contour plots of two clusters connected by a trajectory. For each group, we generated three representative sFNC patterns by uniformly sampling from the derived trajectory.

**Figure 11.**
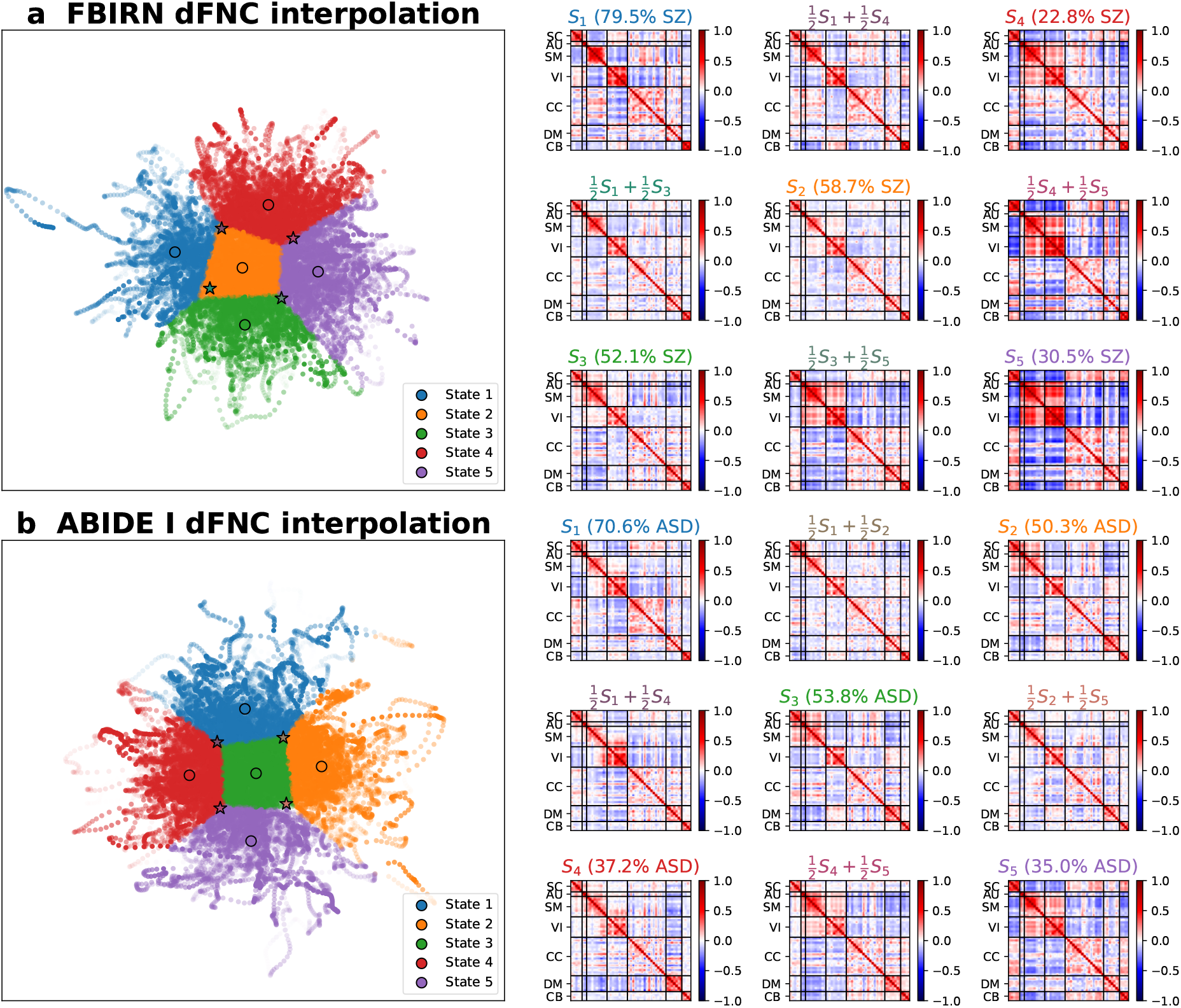
dFNC interpolation between two dynamic states in the VAE latent space. (a) FBIRN training data. (b) ABIDE I training data. Left, we show the dFNC latent features color-coded by *k*-means states. We performed interpolation at the midpoint between two state centroids to examine the transition patterns. Stars (⋆) represent the relative positions in the latent space where the samples were generated. Right, we show the generated samples from each of five state centroids and each of four midpoints between two state centroids.

To visualize how sFNC continuously changes between the control group and the patient group, we interpolated along a trajectory from the control center to the patient center (Figure 10). When interpolating along the trajectory from the control center to the SZ patient center, we observe significant connectivity changes across multiple domains (*p <* 0.0001, Wilcoxon signed-rank test, Bonferroni correction). Specifically, anti-correlations become significantly weaker in the SC–SM, SC–VI, and SM–CB domains, whereas positive correlations become significantly weaker within the SM, VI, and CC domains. In addition, positive correlations gradually shift to negative correlations in the SC–CB and SM–VI domains, while negative correlations shift to positive correlations in the VI–CB domain (Figure 10a). When interpolating along the trajectory from the control center to the ASD patient center, we observe partially overlapping but distinct connectivity patterns. Similar to SZ, negative correlations are significantly stronger in the SC–CB domain but weaker in the VI-CB domain. In contrast, ASD shows unique alterations, including significantly weaker negative correlations in the SM–DM and VI–DM domains, stronger negative correlations in the SC-VI and VI-CC domains, decreased positive correlations within the DM domain, and increased positive correlations within the CC domain (Figure 10b).

To evaluate how a dynamic state switches to another, we generated an FNC matrix by sampling from the distribution at the midpoint between two state centroids in the latent space (Figure 11). When an SZ-dominant state (e.g., state 1) changes to a control-dominant state (e.g., state 5), we observe weaker negative correlations between the SM and VI domains and between the VI and CC domains, and stronger negative correlations between the CB domain and the AU and SM domains (Figure 11a). When an ASD-dominant state (e.g., state 1) switches to a state occupied by more controls (e.g., state 2), we notice a connectivity shift from positive correlations between the SM and VI domains to positive correlations among the SC, AU, and SM domains (Figure 11b).

## 4 Discussion

This work presents a functional network connectivity interpolation framework using a VAE to investigate neuropsychiatric continuum and heterogeneity from static and dynamic FNC data. Our framework provides additional advantages that complement existing diagnostic and analytical methods. To identify neurobiological markers of SZ and ASD, we leveraged functional network connectivity derived from fMRI – an objective measure of brain abnormalities under pathological conditions. To address the limitations of diagnostic labels and supervised prediction, we implemented an *unsupervised generative* model to learn hidden relationships in a fully data-driven manner and generate continuous FNC patterns along the latent disorder spectrum. The correlations between the generated and original sFNC matrices demonstrate that the VAE captured representative and generalizable latent distributions in both datasets, outperforming a linear baseline (PPCA) and a semi-supervised alternative (iVAE) (Figures 2, 6). Notably, the unsupervised VAE slightly outperformed the semi-supervised iVAE that conditioned on the diagnostic label. One reason could be that the iVAE is trained to reduce the total correlation of the latent variable given the auxiliary variable except for optimizing reconstruction (see Khemakhem et al., 2020 Appendix F), and thus its reconstruction performance might not be optimal. In addition, we chose the diagnostic label as the auxiliary variable, yet the most appropriate auxiliary information in the neuroimaging data remains unknown. Regardless, we consider VAEs to be the most suitable for our interpolation task because of their ability to learn nonlinear latent relationships in an unsupervised manner, as supported by empirical evidence.

Unlike prior studies that focus on group-level comparisons, our framework provides a generative approach to model a functional connectivity continuum between groups. Specifically, the interpolated continua from controls to patients in both disorders show two common functional network connectivity gradients: (1) weaker positive correlations within sensory networks (auditory, sensorimotor, visual) and between the subcortical and cerebellar domains; (2) weaker negative correlations between the subcortical domain and the sensory domains, as well as between the cerebellar domain and these sensory regions (Figures 3, 4). Consistent with these findings, previous studies using NeuroMark sFNC in SZ and ASD have reported overlapping group-level alterations in both disorders, particularly between the subcortical domain and the cerebellar, auditory, and sensorimotor domains (Du et al., 2020; Jensen et al., 2024; Yan et al., 2024). We also observe disorder-specific patterns. For example, SZ is characterized by reduced functional connectivity in the visual domain, whereas ASD shows decreased functional connectivity within the default mode network, aligning with prior work (Du et al., 2020).

Another key benefit of our interpolation framework is to visualize and characterize individual differences. Instead of solely relying on group averages, we designed a 2D grid display to organize sFNC matrices, supporting examination of subject-level variability and inter-subject relationships (Figures 3, 4). Apart from sFNC matrices, we projected multiple subject measures, including clinical and cognitive scores, on the same 2D grid (Figure 5). Notably, SZ patients grouped in the lower/left/lower triangular half of the 2D grid consistently show higher cognitive scores, indicating that these patients have better cognitive functions (Table 2).

Moreover, we identified subgroups among SZ patients by applying the *k*-means clustering algorithm on the learned latent features and then examined subgroup differences in subject measures and functional network connectivity patterns (Appendix J). Changes related to working memory were observed across multiple network connections, including sensorimotor–default mode, sensorimotor–cerebellar, and cognitive control–cerebellar networks. Previous studies have similarly linked working memory deficits in SZ to weaker connectivity between the fronto-parietal network and cerebellar network (Repovs et al., 2011), as well as stronger within-network connectivity in the fronto-parietal and default mode networks (Unschuld et al., 2014). The default mode network was also associated with speed of processing, consistent with previous reports (Mwansisya et al., 2013; Wang et al., 2014). In addition, attention performance was associated with connectivity changes between the default mode and cognitive control networks and each of the sensorimotor, cerebellar, and auditory domains. This finding is in line with prior work proposing the insula as a key hub that modulates activity between the default mode network and task-positive networks (Fox et al., 2005; Z. He et al., 2013; Moran et al., 2013).

We further examined how sFNC patterns vary across cognitive performance levels by interpolating along a trajectory from the higher-score group to the lower-score group (Appendix K). Existing literature suggests that cognitive deficits in SZ are associated with reduced resting-state functional connectivity between cortical and subcortical regions (Sheffield & Barch, 2016). Consistent with this evidence, we found that, compared with the higher-score group, the lower-score group exhibited significantly attenuated negative correlations in the subcortical–sensorimotor, subcortical–visual, and sensorimotor–cerebellar domains, as well as reduced positive correlations within the sensorimotor, visual, and cognitive control domains, supporting the role of disrupted cortical–subcortical integration in cognitive impairment.

In addition to static brain connectivity, we evaluated whether the VAE can capture dynamic properties from windowed dFNC data. We observe highly similar *k*-means states between the generated and original results (Figures 7, 8). These generated states exhibit consistent group-specific properties, measured by dwell time and transition probability, as the original states (Figure 9). In particular, the dynamic state analysis in both the generated and original states suggests that SZ and ASD patients tend to stay longer in sparsely connected states while controls tend to stay longer in densely connected states, consistent with prior literature (Damaraju et al., 2014; Fu et al., 2019, 2021).

A VAE is designed to learn latent distributions and capture hidden relationships from observed data in an unsupervised manner. By leveraging the VAE latent space, we performed interpolation along a pathological trajectory to visualize continuous sFNC alterations from the control center to the patient center (Figure 10). Similarly, we generated samples between two dFNC state centroids to examine how one state gradually switches to another (Figure 11). Overall, we demonstrate the ability of the unsupervised generative model to capture representative group-specific functional network connectivity profiles and generate continuous pattern alterations between different subjects, groups, dynamic states, and time points.

Despite the promising potential of our interpolation framework, we acknowledge limitations in this work. First, we implemented a vanilla VAE with a multivariate Gaussian prior assumption, but it might not be fully capable of capturing the heterogeneity or potential hierarchical structures in the SZ and ASD datasets. Second, in order to generate continuous sFNC data, we sampled from linearly spaced coordinates in the latent space, rather than the posterior distributions, so the generated sFNC matrices and the original sFNC matrices ordered by the JV algorithm were not one-to-one matched due to uniform sampling. Lastly, we made an assumption that the differences in FNC between groups, which reflect the underlying neurobiological changes in psychiatric disorders, are continuous and can be interpolated. However, there has been debate about whether psychosis can be viewed as continua or categories (David, 2010; DeRosse & Karlsgodt, 2015; Lawrie et al., 2010). More recently, there is growing evidence that suggests the shift from the concept of schizophrenia to a broader psychotic spectrum (Guloksuz & van Os, 2018) that captures a continuum of psychosis proneness from normal to abnormal (Van Os et al., 2009). Hence, it is worth further quantifying the associations between the interpolated FNC patterns and the subject measures, including clinical symptoms and cognitive functions.

In the future, we aim to extend the framework to further characterize psychotic syndromes using hierarchical clustering or mixture model priors in VAEs, and to automatically capture dynamic states using dynamical VAEs (Girin et al., 2021). We will also more systematically quantify the relationships between functional network connectivity patterns and cognitive functions across patient subgroups. To improve generalizability and establish more reliable continuum prediction, we will analyze other large-scale longitudinal datasets with multiple mental disorders, such as the Adolescent Brain and Cognitive Development (ABCD) dataset (Barch et al., 2018; Casey et al., 2018; Garavan et al., 2018).

## 5 Conclusions

We present a functional network connectivity interpolation framework using an unsupervised generative model to interpolate sFNC and dFNC data from controls and patients with SZ or ASD. We demonstrated a high degree of correspondence between the generated and original individual sFNC patterns, as well as the group-specific dFNC states. We showed that our interpolation framework captured individual variability within a group, sFNC pattern alterations between groups, and dFNC dynamic states between groups. In addition, we provided examples of interpolating along trajectories of interest in the latent manifold to visualize the corresponding continuous FNC alterations in the original space. Our approach offers new insights into the neuropsychiatric continuum and heterogeneity, marking a promising step toward personalized characterization of mental disorders.

## Ethics statement

The authors have no conflict of interest to declare. Large language models such as Claude and ChatGPT were used to correct grammar mistakes at the sentence level.

## Data and code availability

The FBIRN dataset can be accessed at https://www.nitrc.org/projects/fbirn/. The ABIDE I dataset can be accessed at https://fcon_1000.projects.nitrc.org/indi/abide/. The NeuroMark network templates are available at http://trendscenter.org/software. The analysis and visualization code is publicly available at https://github.com/XinhuiLi/interpolation.git.

## Author contributions

**Xinhui Li:** Conceptualization; formal analysis; investigation; methodology; software; visualization; writing - original draft; writing - review and editing. **Eloy Geenjaar:** Investigation; methodology; writing - review and editing. **Zening Fu:** Data curation; investigation; writing - review and editing. **Godfrey D. Pearlson:** Investigation; writing - review and editing. **Vince D. Calhoun:** Conceptualization; funding acquisition; investigation; methodology; project administration; resources; supervision; writing - review and editing.

## Funding

This work was supported by the National Institutes of Health (NIH) grants (R01MH118695 and R01EB006841) and the National Science Foundation (NSF) grant (NSF2112455). Additionally, X.L. and E.G. were supported by the Georgia Tech/Emory NIH/NIBIB Training Program in Computational Neural-engineering (T32EB025816).

## Declaration of competing interests

The authors have no competing interest to declare.

## Acknowledgments

We thank the FBIRN team for collecting and sharing the dataset, including Adrian Preda, Aysenil Belger, Bryon A. Mueller, Daniel H. Mathalon, Daniel S. O’Leary, Jessica A. Turner, Juan R. Bustillo, Judith M. Ford, Kelvin O. Lim, Steven G. Potkin, and Theo G.M. van Erp. We also thank the ABIDE team for collecting, organizing and sharing the dataset. We acknowledge the NeuroImaging Tools and Resources Collaboratory (NITRC) team for providing the data sharing platform.

## Appendix A ABIDE I data information

In the ABIDE I dataset, we used 266 subjects from the following three sites: NYU Langone Medical Center (NYU), University of California, Los Angeles: Sample 1 (UCLA), and University of Utah School of Medicine (USM). Site-specific demographic information is shown in Table A.1. fMRI acquisition parameters for each site are shown in Table A.2.

Based on the mapping of latent features to 2D grid nodes (as in Figure 4), we show diagnosis labels, autism diagnostic observation schedule (ADOS) scores, age and gender information of ABIDE I subjects on the 2D grid (Figure A.1). We note that the diagnosis label plays a key role in grouping subjects, with most ASD patients clustered in the upper triangle and most controls in the lower triangle.

**Table A.1:** ABIDE I site-specific demographic information. *N*: number of total subjects; *N*_male_: number of male subjects; *N*_female_: number of female subjects; s.d.: standard deviation.

| Site | $N$ | $N_{\text{male}}$ | $N_{\text{female}}$ | Age mean ± s.d. (years) | Age range (years) |
| --- | --- | --- | --- | --- | --- |
| ASD |  |  |  |  |  |
| NYU | 64 | 55 | 9 | $15.72 \pm 7.11$ | 7.15 – 39.10 |
| UCLA | 41 | 36 | 5 | $13.54 \pm 2.25$ | 9.26 – 17.94 |
| USM | 28 | 28 | 0 | $25.05 \pm 6.11$ | 15.93 – 37.78 |
| CTR |  |  |  |  |  |
| NYU | 95 | 70 | 25 | $16.45 \pm 6.12$ | 7.29 – 31.78 |
| UCLA | 29 | 25 | 4 | $13.35 \pm 1.96$ | 9.50 – 17.79 |
| USM | 9 | 9 | 0 | $20.57 \pm 4.65$ | 13.81 – 27.60 |

**Table A.2:**
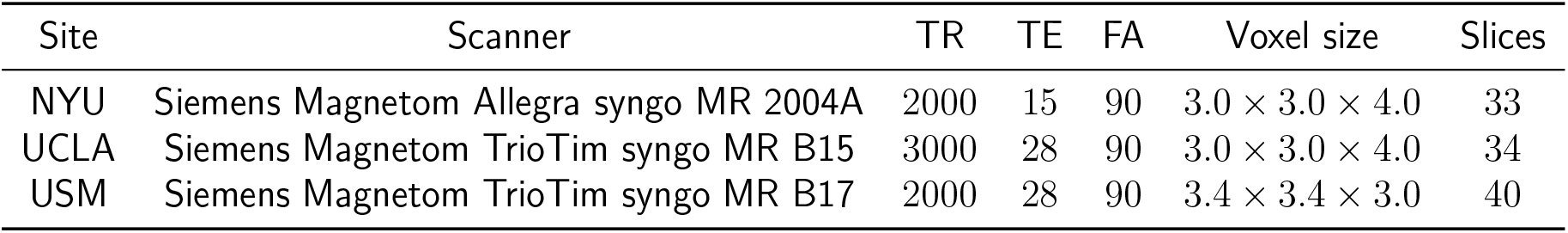
ABIDE I site-specific fMRI acquisition parameters. Parameters (units) are shown as follows: TR (ms), TE (ms), FA (degree), voxel size (mm).

| Site | Scanner | TR | TE | FA | Voxel size | Slices |
| --- | --- | --- | --- | --- | --- | --- |
| NYU | Siemens Magnetom Allegra syngo MR 2004A | 2000 | 15 | 90 | $3.0 \times 3.0 \times 4.0$ | 33 |
| UCLA | Siemens Magnetom TrioTim syngo MR B15 | 3000 | 28 | 90 | $3.0 \times 3.0 \times 4.0$ | 34 |
| USM | Siemens Magnetom TrioTim syngo MR B17 | 2000 | 28 | 90 | $3.4 \times 3.4 \times 3.0$ | 40 |

**Figure A.1:**
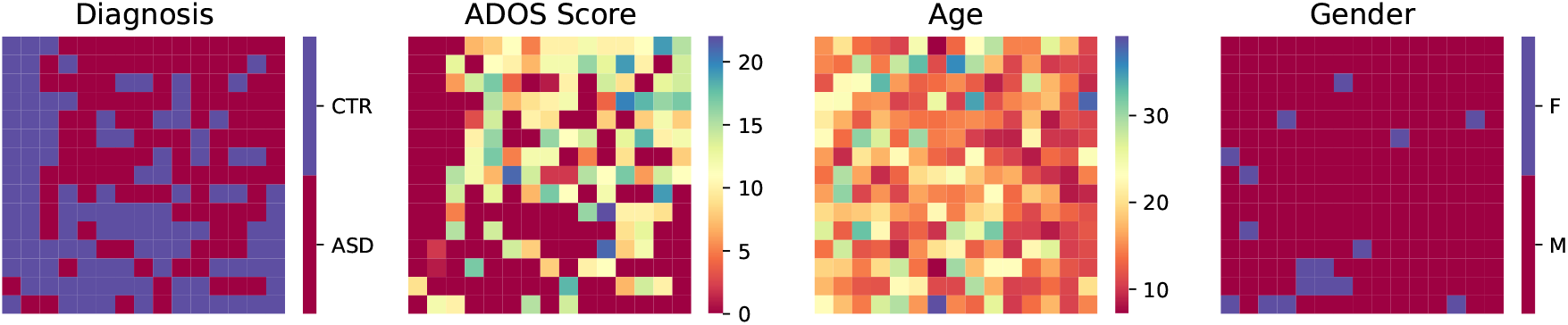
ABIDE I subject measures displayed on the 2D grid. Diagnosis labels, autism diagnostic observation schedule (ADOS) scores, age and gender information of ABIDE I subjects were displayed on the 2D grid correspondingly.

## Appendix B Statistics of subject measures

Table B.1 summarizes the group-level mean, standard deviation, and range for each subject measure (diagnostic or cognitive score) in each dataset.

**Table B.1:** Statistics of subject measures. PT: patient; CTR: control.

| Subject measure | PT mean ± s.d. | PT range | CTR mean ± s.d. | CTR range |
| --- | --- | --- | --- | --- |
| FBIRN |  |  |  |  |
| PANSS (positive) | $14.976 \pm 4.75$ | 7 – 29 | $7.000 \pm 0.00$ | 7 – 7 |
| PANSS (negative) | $13.928 \pm 5.00$ | 7 – 30 | $7.000 \pm 0.00$ | 7 – 7 |
| Speed of processing | $-1.217 \pm 0.98$ | -3.682 – 1.165 | $0.038 \pm 1.00$ | -3.764 – 2.195 |
| Attention/Vigilance | $-1.277 \pm 1.36$ | -4.965 – 1.390 | $0.026 \pm 0.98$ | -4.965 – 1.542 |
| Working memory | $-1.018 \pm 0.97$ | -3.947 – 2.053 | $0.020 \pm 0.96$ | -3.019 – 2.053 |
| Verbal learning | $-1.313 \pm 1.17$ | -4.034 – 1.561 | $0.042 \pm 0.99$ | -3.101 – 1.561 |
| Visual learning | $-1.002 \pm 1.13$ | -2.873 – 1.152 | $0.026 \pm 0.99$ | -2.873 – 1.646 |
| Reasoning/Problem solving | $-0.756 \pm 1.11$ | -4.443 – 0.981 | $0.022 \pm 0.97$ | -4.805 – 0.981 |
| Composite score | $-1.562 \pm 1.22$ | -4.502 – 1.496 | $0.044 \pm 0.95$ | -4.023 – 1.550 |
| ABIDE I |  |  |  |  |
| ADOS score | $11.594 \pm 4.00$ | 2 – 22 | $0.090 \pm 0.47$ | 0 – 4 |

## Appendix C Variational autoencoders

Here, we provide the step-by-step derivation of the VAE objective function. We refer readers to Kingma and Welling, 2014 for the original theory and to Tiao, 2017 for a detailed tutorial.

Let x ∈ ℝ^*V*^ be an observed data variable and z ∈ ℝ^*D*^ be its corresponding latent variable. We assume that the latent variable z is sampled from a prior distribution *p*(z) and the observed variable x is sampled from the conditional likelihood distribution *p*_*θ*_(x|z) parameterized by *θ*. We consider a latent variable model with the following joint distribution:

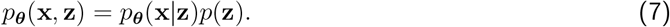

We would like to estimate the posterior distribution in order to perform inference on the latent variable. According to Bayes’ theorem, the posterior can be computed as:

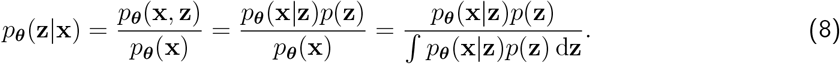

The denominator *p*_*θ*(x) = ∫_*p*_*θ*_(x|z)*p*(z) dz is the marginal distribution of the observed data, also known as the *evidence*. It is analytically intractable because there is no closed form for the integral term.

To overcome this intractability, we leverage *variational inference*, which assumes a simpler distribution to approximate the true posterior. We want to estimate the optimal variational parameters *ϕ*^∗^by minimizing the Kullback-Leibler (KL) divergence^2^ between the approximate posterior *q*_*ϕ*_(z|x) and the true posterior *p*_*θ*_(z|x):

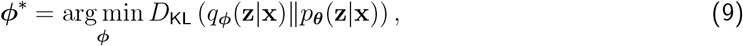

where *D*_KL_ (*q*_*ϕ*_(z|x)∥*p*_*θ*_(z|x)) = 0 if and only if *q*_*ϕ*_(z|x) = *p*_*θ*_(z|x). This is again intractable due to *p*_*θ*_(z|x). However, note that the true posterior is proportional to the joint distribution, *p*_*θ*_(z|x) ∝ *p*_*θ*_(x|z)*p*(z). Thus, we can minimize the following KL divergence instead:

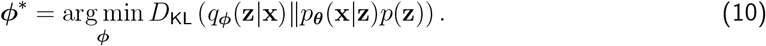

The *evidence lower bound* (ELBO) is defined to maximize the negative KL divergence in Equation 10 by optimizing the generative parameters *θ* and the variational parameters *ϕ*:

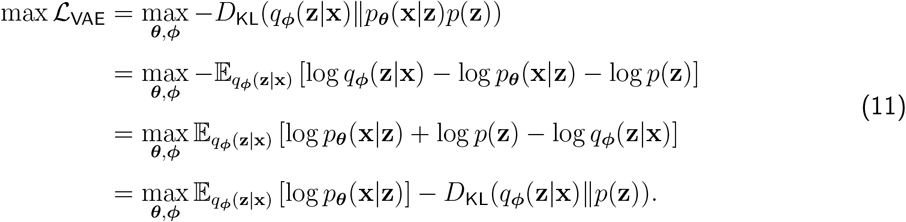

Crucially, maximizing the ELBO with respect to the generative parameters *θ* approximately maximizes the log marginal likelihood of the observed data, while maximizing it with respect to the variational parameters *ϕ* equivalently minimizes the KL divergence. Also, note that the KL divergence is non-negative, i.e. *D*_KL_(·) ≥ 0. Thus, the ELBO is the lower bound to the log likelihood of generating the observed data, i.e. ℒ_ELBO_ ≤ log *p*_*θ*_(x).

In this work, we assume that both the variational posterior and the prior follow multivariate Gaussian distributions:

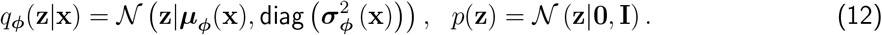

Then the KL term in Equation 11 can be rewritten as:

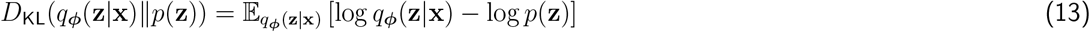

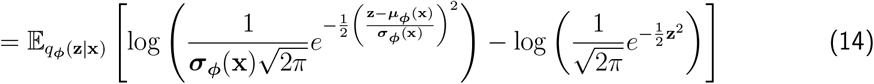

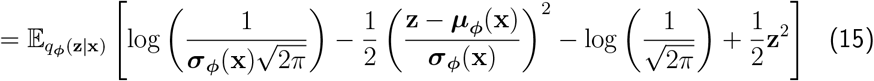

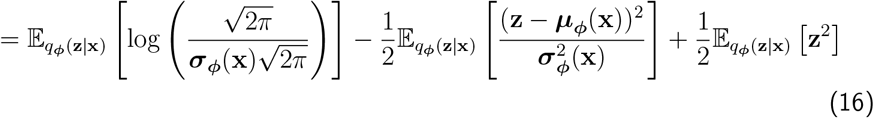

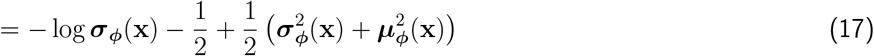

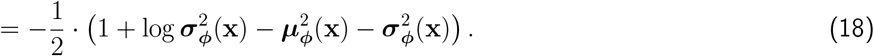

To simplify Equation 16 to Equation 17, we leverage the definition of variance,

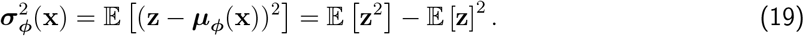

Thus, 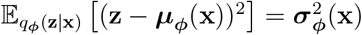 and 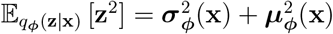.

In order to use gradient descent, the VAE objective function is defined to minimize the reconstruction error measured by the mean squared error (MSE) and the KL divergence in Equation 18, which equivalently minimizes the negative ELBO in Equation 11:

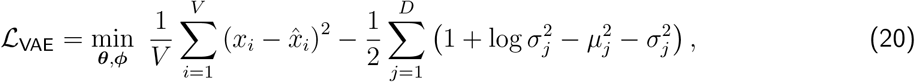

where *x*_*i*_ and 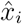 are the *i*-th element of the observed data x and the reconstructed data 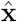, respectively; *µ*_*j*_ and *σ*_*j*_ are the *j*-th element of *µ*_*ϕ*_(x) and *σ*_*ϕ*_(x), respectively.

To train the ELBO with gradient-based optimization, we use the *reparameterization trick* which expresses the random variable z as a deterministic transformation of the input x and a Gaussian random variable with zero mean and unit variance *ϵ* ∈ ℝ^*D*^:

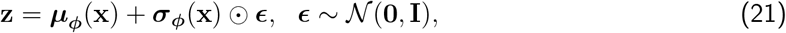

where ⊙ represents the element-wise product.

## Appendix D Probabilistic principal component analysis

In PPCA, we assume the prior distribution over the latent variable z ∈ ℝ^*D*^ and the conditional distribution of the observed variable x ∈ ℝ^*V*^ given z follow Gaussian distributions:

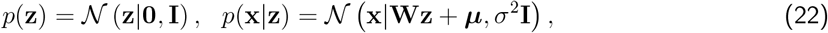

where the matrix W ∈ ℝ^*V ×D*^ and the vector *µ* ∈ ℝ^*V*^ define the mean of x, and the scalar *σ*^2^ ∈ R determines the variance of the conditional distribution.

The marginal distribution of the observed data is also a Gaussian:

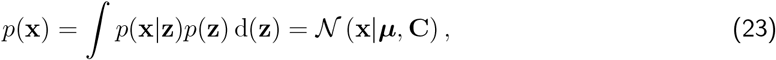

where the covariance matrix C ∈ ℝ^*V ×V*^is defined by C = WW^⊤^ + *σ*^2^I. Note that x = Wz + *µ* + *ϵ, ϵ* ∼ 𝒩 (0, *σ*^2^I). Thus, 𝔼 [x] = 𝔼 [Wz + *µ* + *ϵ*] = *µ* and cov[x] = 𝔼 [(Wz + *ϵ*)(Wz + *ϵ*)^⊤^] = 𝔼 [Wzz^⊤^W^⊤^] + 𝔼 [*ϵϵ*^⊤^] = WW^⊤^ + *σ*^2^I.

Using Bayes’ rule, the posterior distribution can be derived as:

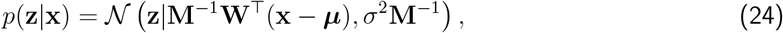

where the matrix M ∈ 𝔼^*D×D*^ is given by M = W^⊤^W + *σ*^2^I.

_Given a dataset of observed samples 𝒟 =_ x^(1)^, …, x^(*i*)^, …, x^(*N*)^, the corresponding log likelihood function can be written as:

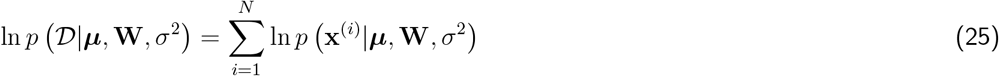

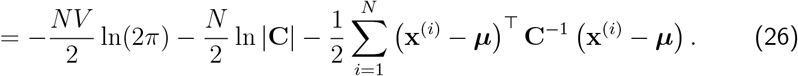

Using maximum likelihood estimation, we obtain the model parameter 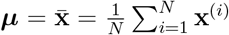 by setting the derivative of the log likelihood with respect to *µ* equal to zero.

We implement the expectation-maximization (EM) algorithm to estimate the model parameters W and *σ*^2^. In the E step, we use the old parameter values to evaluate the expectation of the complete-data log likelihood with respect to the posterior distribution:

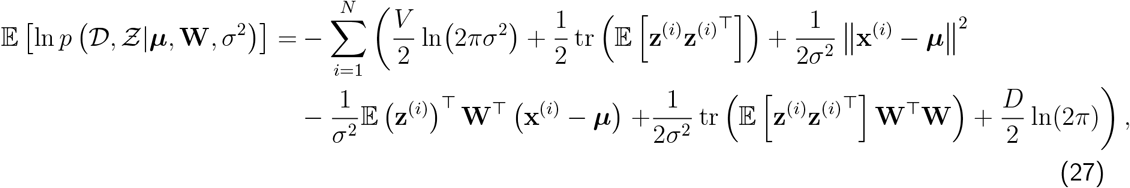

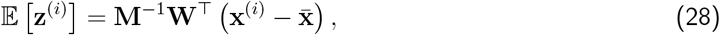

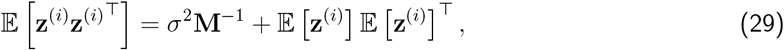

where z^(*i*)^_is the *i*-th element in the set of latent variables *Ƶ* = {_z^(1)^, …, z^(*i*)^, …, z^(*N*)^}.

In the M step, we maximize the the expected complete-data log likelihood with respect to W and *σ*^2^, yielding the new parameter values:

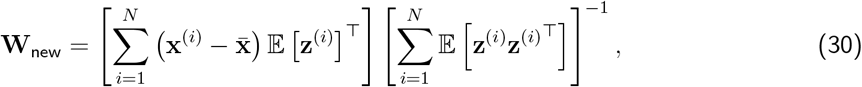

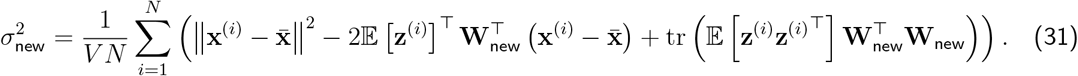

We iteratively alternate between the E step and the M step, until the parameter values converge. For more details, we refer readers to Tipping and Bishop, 1999 or Chapter 16, Bishop and Bishop, 2023.

## Appendix E Identifiable variational autoencoders

In iVAE, we consider the following conditional generative model:

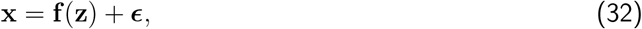

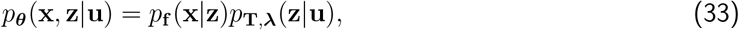

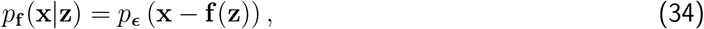

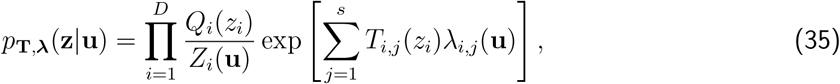

where x ∈ ℝ^*V*^is the observed data variable, u ∈ ℝ^*U*^is the observed auxiliary variable, z ∈ ℝ^*D*^ is the latent variable, *ϵ* ∈ ℝ^*V*^is a noise variable with probability density function *p*_*ϵ*_(*ϵ*), *θ* = (f, T,*λ*) is a set of parameters of the generative model, and f : ℝ^*D*^ → ℝ^*V*^is a nonlinear mixing function. Since our main interest lies in psychiatric characteristics, we use the one-hot encoded diagnostic label as the auxiliary variable u in the iVAE (*U* = 2). We assume that the prior on the latent variable *p*_T,*λ*_(z|u) is conditionally independent, and each *z*_*i*_follows a univariate exponential family distribution given the auxiliary variable u, where *Q*_*i*_is the base measure, *Z*_*i*_(u) is the normalizing constant, T_*i*_ = (*T*_*i*,1_, …, *T*_*i,s*_) are the sufficient statistics, *λ*_*i*_(u) = (*λ*_*i*,1_(u), …, *λ*_*i,s*_(u)) are the parameters depending on u, and *s* is the dimension of each sufficient statistic.

Similar to VAE, the iVAE objective function aims to learn the parameters (*θ, ϕ*) that maximize the data generation likelihood by maximizing the ELBO:

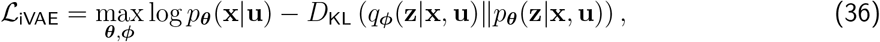

where *p*_*θ*(x|u) =∫_*p*_*θ*_(x, z|u) dz is the conditional marginal distribution of the observed data, and *q*_*ϕ*_(z|x, u) is the variational posterior of its true posterior *p*_*θ*_(z|x, u).

For more information, please refer to Khemakhem et al., 2020.

## Appendix F Model architecture search

### F.1 VAE architecture search

We evaluated four VAE architectures based on multilayer perceptrons (MLPs). Each architecture has a different set of output units in the VAE encoder and the corresponding symmetric output units in the VAE decoder. The number of output units in each encoder layer is listed in Table F.1. In each layer, we used a Leaky Rectified Linear Unit (Leaky ReLU) activation function (Maas et al., 2013) with a negative slope of 0.5.

For each dataset (FBIRN or ABIDE I) and each data type (sFNC or dFNC), we evaluated each of four potential model architectures separately. For each architecture, we used 10 different random seeds to initialize model weights. As shown in Figure F.1, we find that smaller models with 2 or 3 layers achieved optimal performance for sFNC data, while larger models with 5 or 7 layers worked better for dFNC data. Based on the training performance across 10 runs, we selected the following VAE architectures as shown in Table F.2.

**Table F.1:**
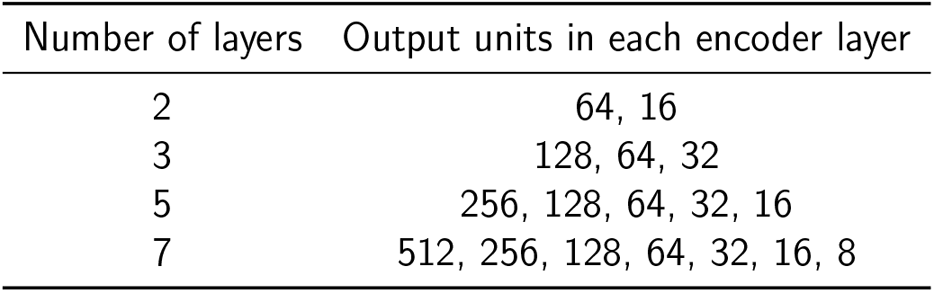
Evaluated VAE architectures.

**Table F.2:** Selected VAE architectures.

| Dataset | Data type | Number of layers | Output units in each encoder layer |
| --- | --- | --- | --- |
| FBIRN | sFNC | 3 | 128, 64, 32 |
| FBIRN | dFNC | 7 | 512, 256, 128, 64, 32, 16, 8 |
| ABIDE I | sFNC | 2 | 64, 16 |
| ABIDE I | dFNC | 5 | 256, 128, 64, 32, 16 |

### F.2 IVAE architecture search

**Figure F.1:**
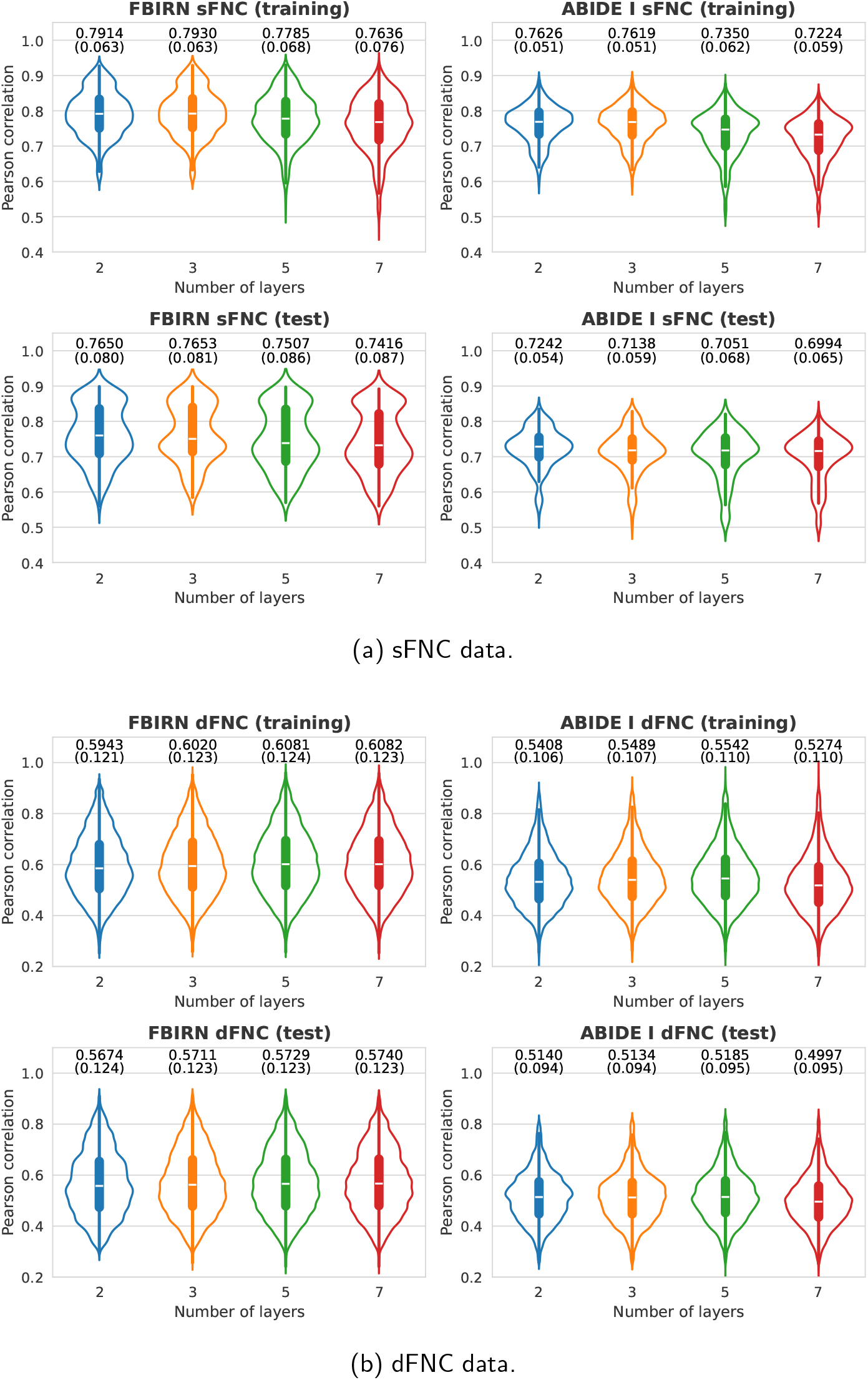
VAE architecture search. Each violin plot shows a symmetric kernel density estimate of Pearson correlations between original and generated sFNC matrices across 10 random seeds, along with the quartiles represented by a box plot.

We evaluated 12 MLP-based encoder-decoder architectures, varying the number of layers (2, 3, 4) and hidden dimensions (16, 32, 64, 96). According to the training performance (Figure F.2), we selected the iVAE model with 3 layers and 16 hidden units for FBIRN, and the model with 2 layers and 32 hidden units for ABIDE I.

**Figure F.2:**
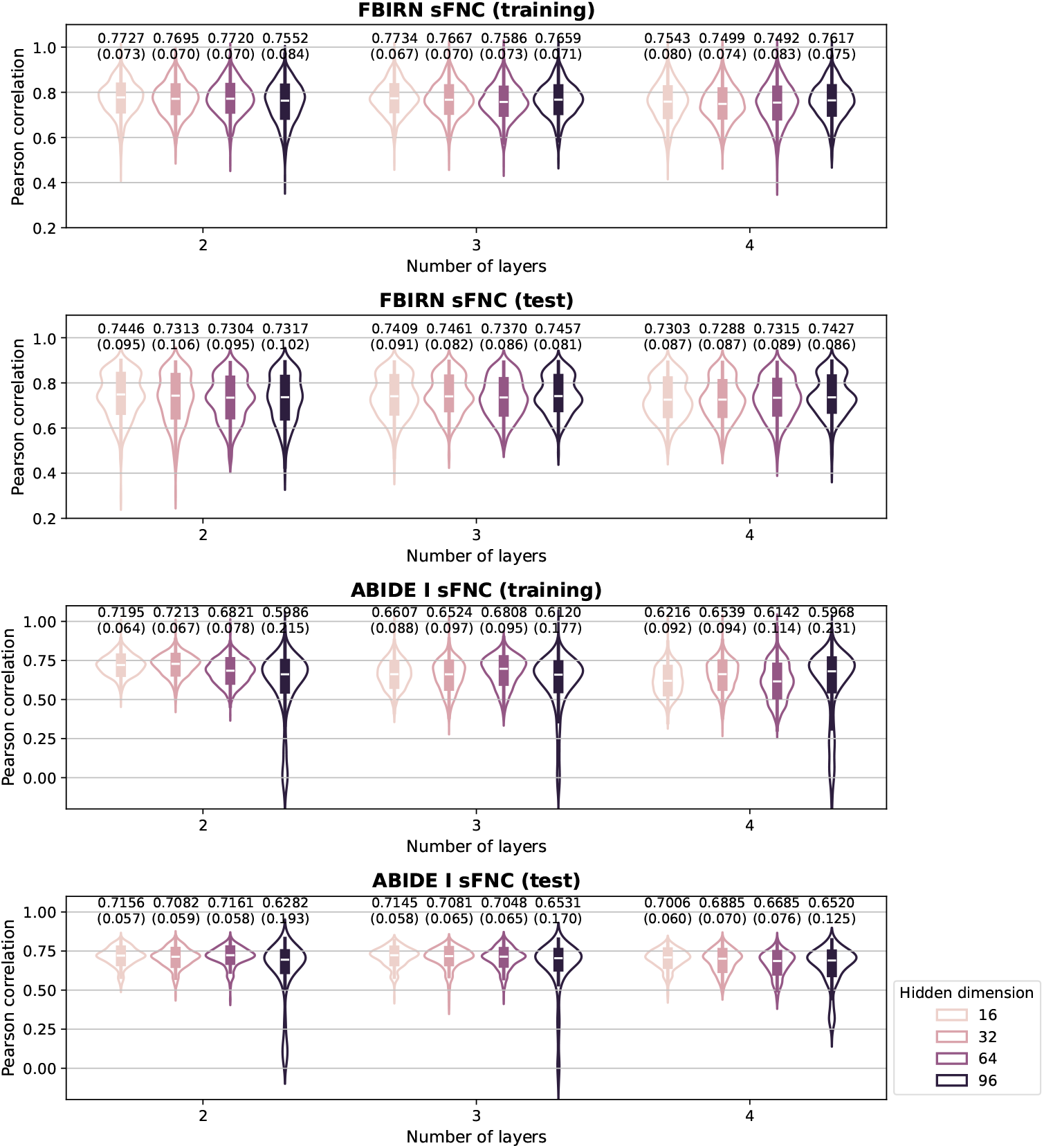
IVAE architecture search for sFNC data. Each violin plot shows a symmetric kernel density estimate of Pearson correlations between original and generated sFNC matrices across 10 random seeds, along with the quartiles represented by a box plot.

## Appendix G Confound effects on latent representations

To assess the robustness of the learned representations to potential confounding variables (e.g., age, sex, and collection site), we regressed these variables out of the latent representation (z) using linear regression. For the FBIRN dataset, we constructed a confound matrix C = [1, c_age_, c_sex_] ∈ ℝ^*N×*3^, where 1 ∈ ℝ^*N*^ is a column vector of ones, c_age_ ∈ ℝ^*N*^ is the demeaned linear age vector, and c_sex_ ∈ ℝ^*N*^ is the binary sex vector (0: female; 1: male). For the multi-site ABIDE I dataset, we also included the site information in the confound matrix, C = [1, c_age_, c_sex_, c_site_] ∈ ℝ^*N×*5^, where c_site_ ∈ ℝ^*N×*2^encodes the site information ([0, 0]: NYU; [1, 0]: UCLA; [0, 1]: USM). The latent representation after regressing out confounding variables (z^′^) is calculated as follows:

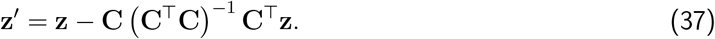

Next, we calculated the randomized dependence coefficient (RDC) (Lopez-Paz et al., 2013), a measure of nonlinear dependence, between the representations before and after regression. An RDC value is bounded between 0 and 1, with higher values indicating greater similarity.

As shown in Figure G.1, the RDC values along the diagonal were very high (FBIRN: 0.989, 0.970; ABIDE I: 0.976, 0.955), indicating high similarity between the representations before and after regression. These results suggest that the learned representations appear to be largely insensitive to the examined confounding factors.

**Figure G.1:**
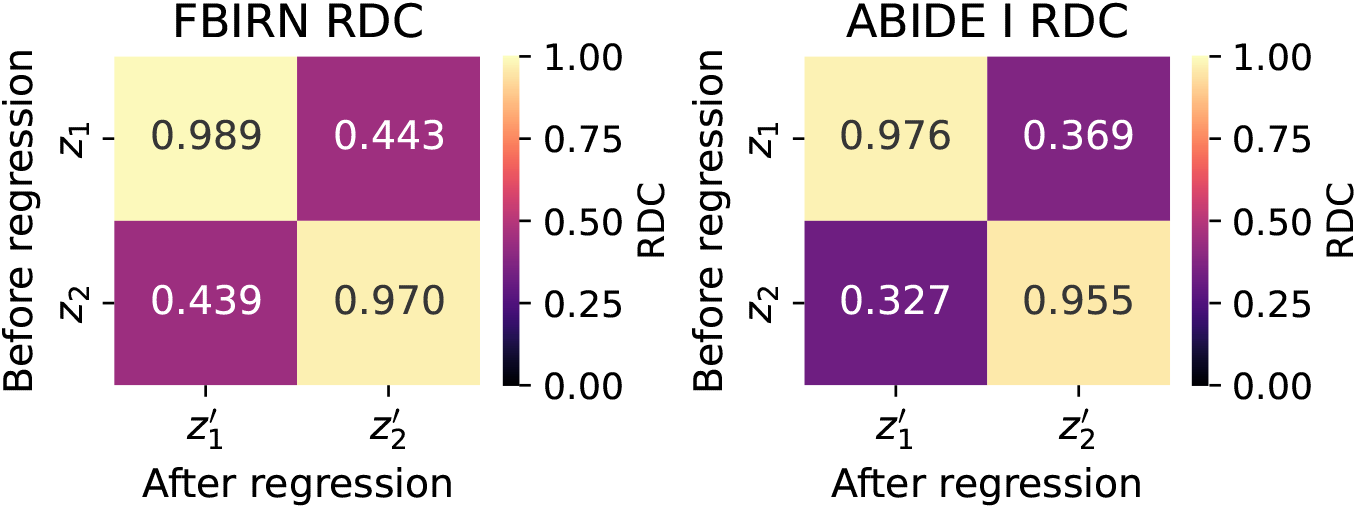
Randomized dependence coefficient (RDC) between the latent representations before and after regressing out confounding variables. High diagonal RDC values indicate strong similarity between representations before and after regression, suggesting that the learned representations are largely unaffected by the examined confounding factors.

## Appendix H sFNC visualization

### H.1 FBIRN

**Figure H.1:**
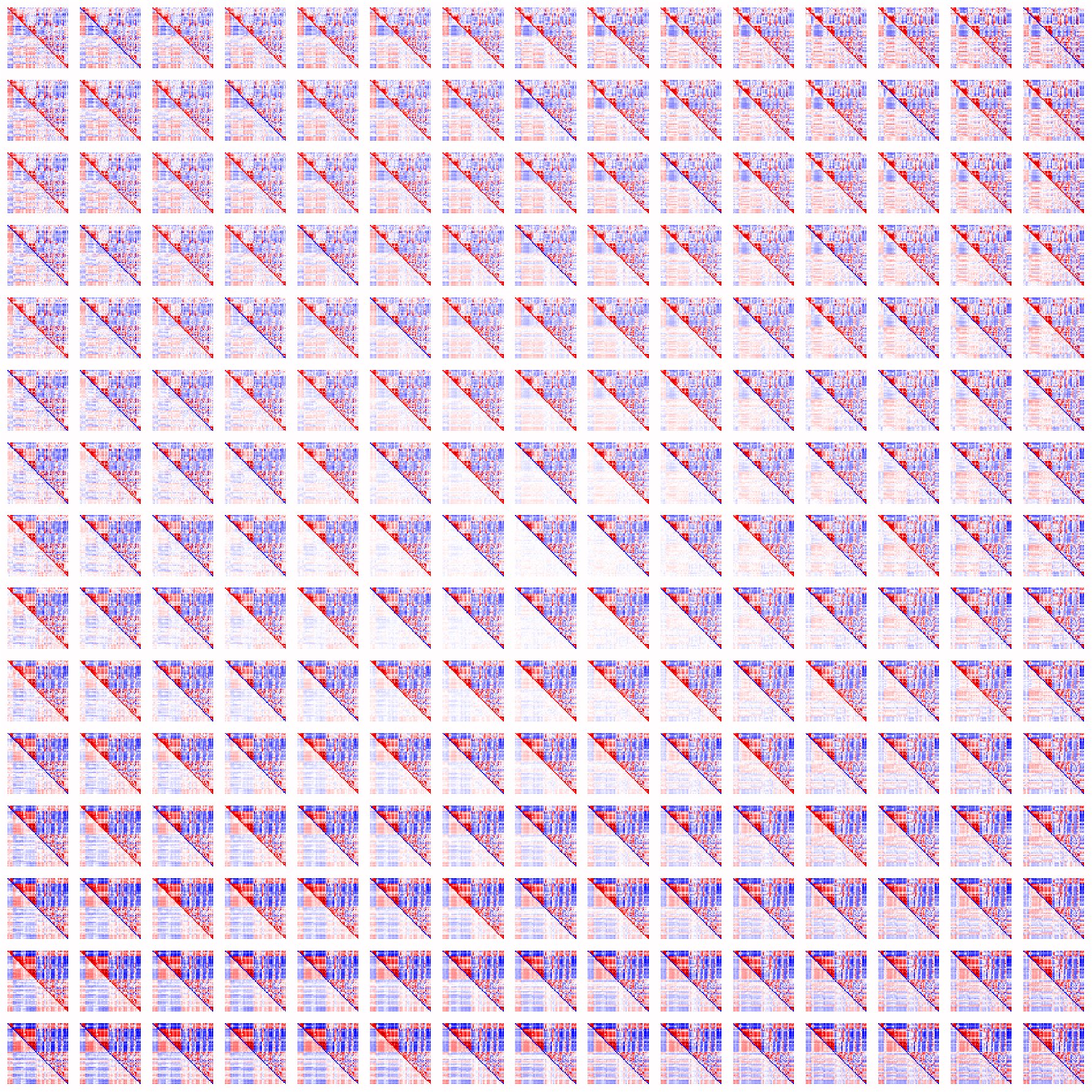
FBIRN generated sFNC (training).

**Figure H.2:**
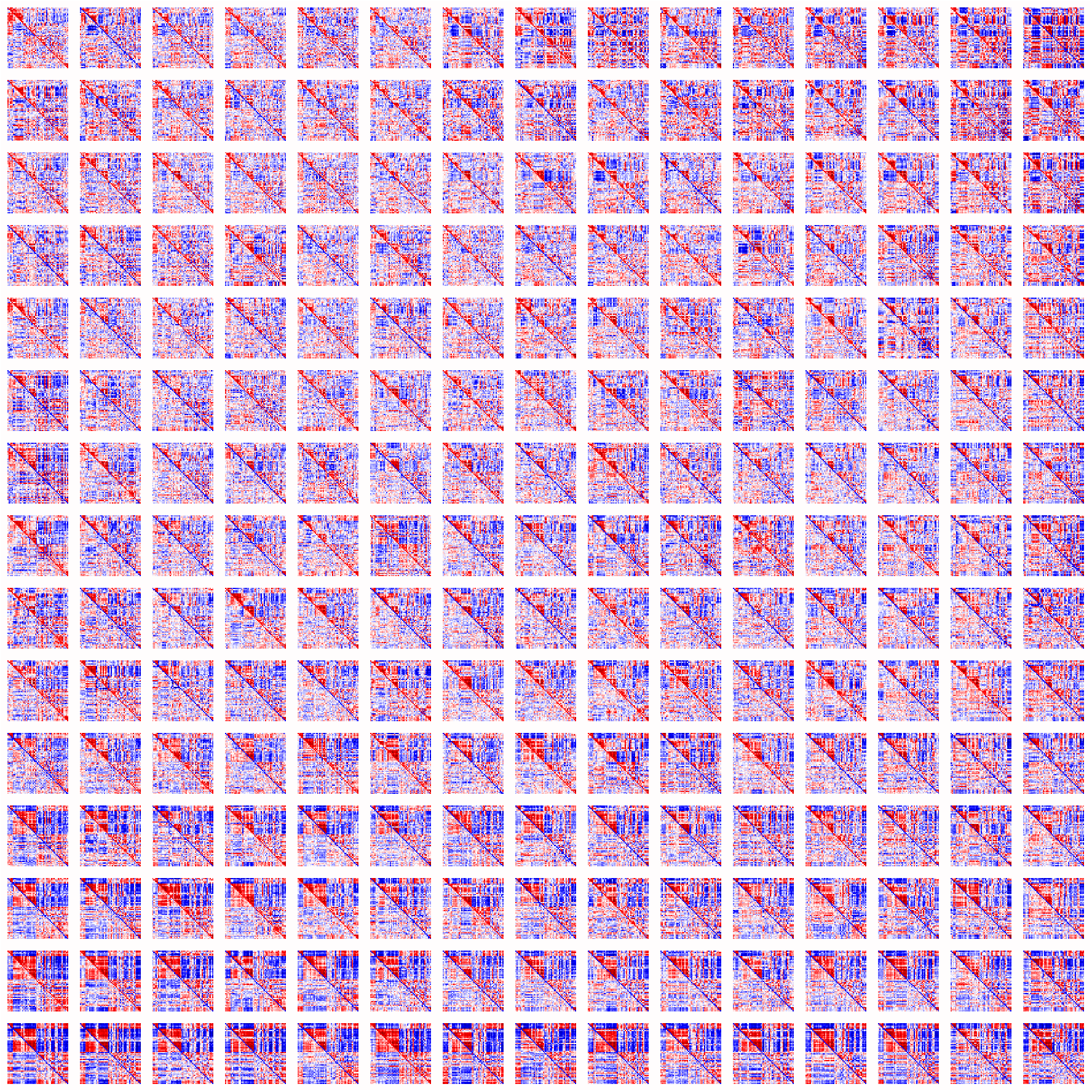
FBIRN original sFNC (training).

**Figure H.3:**
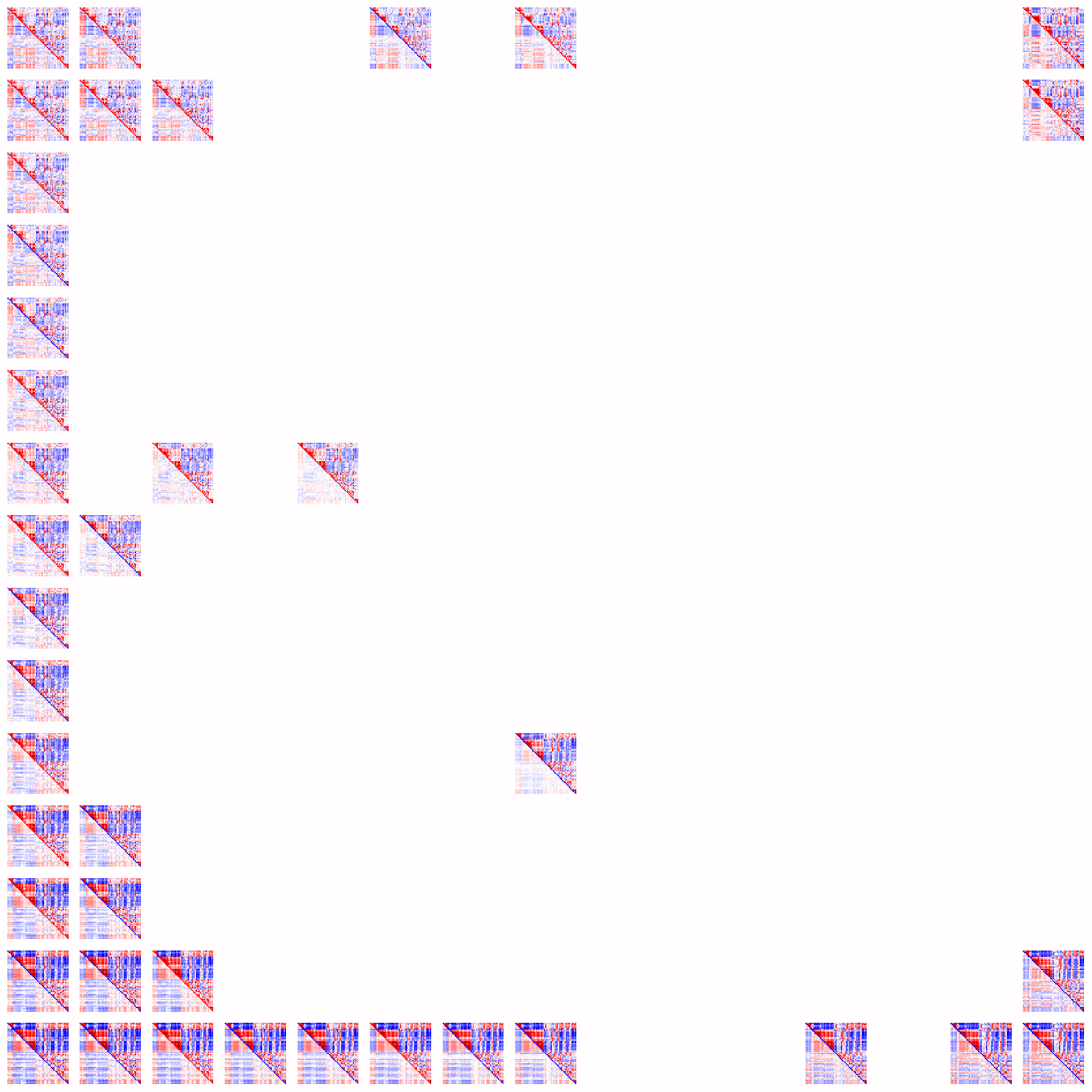
FBIRN generated sFNC (test).

**Figure H.4:**
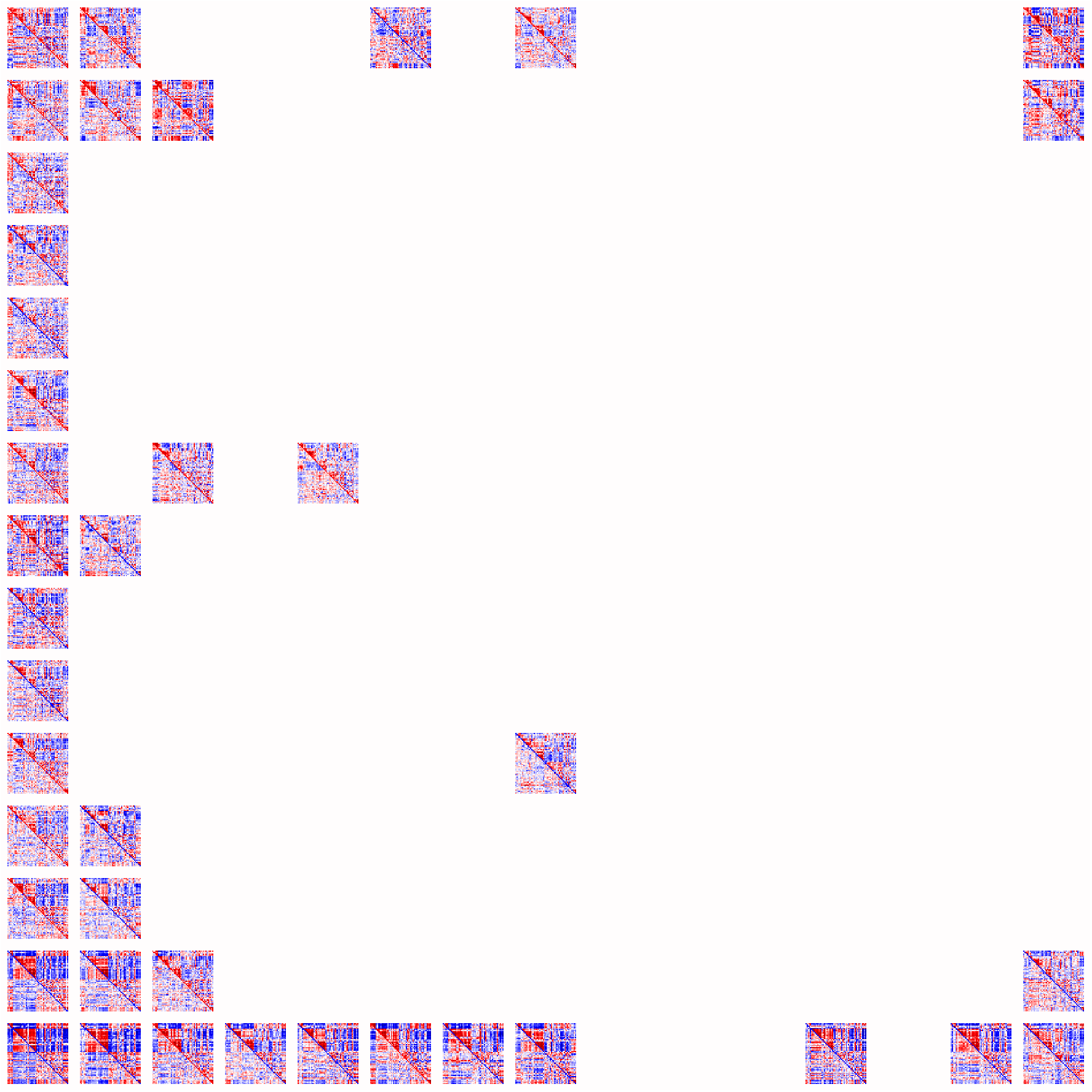
FBIRN original sFNC (test).

### H.2 ABIDE I

**Figure H.5:**
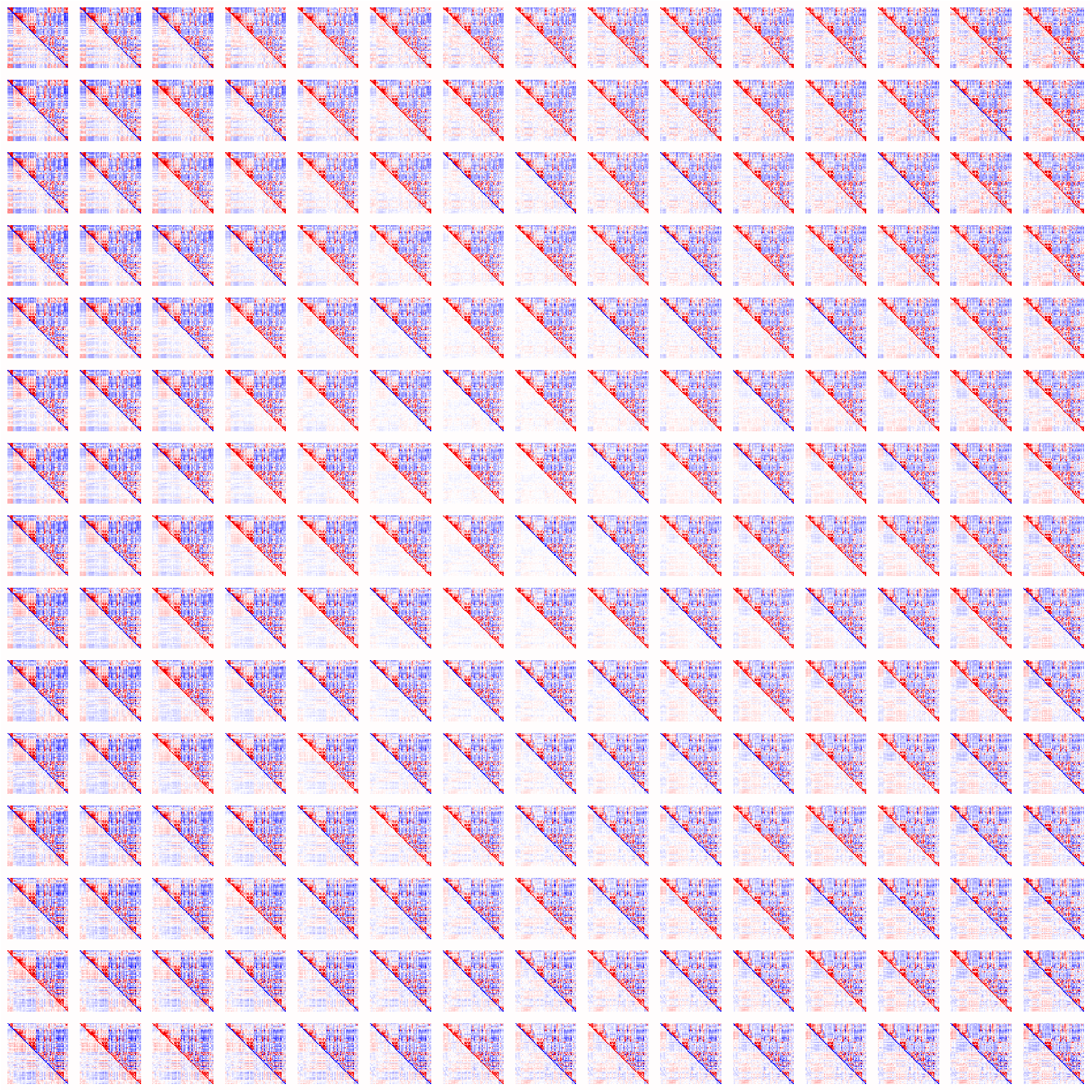
ABIDE I generated sFNC (training).

**Figure H.6:**
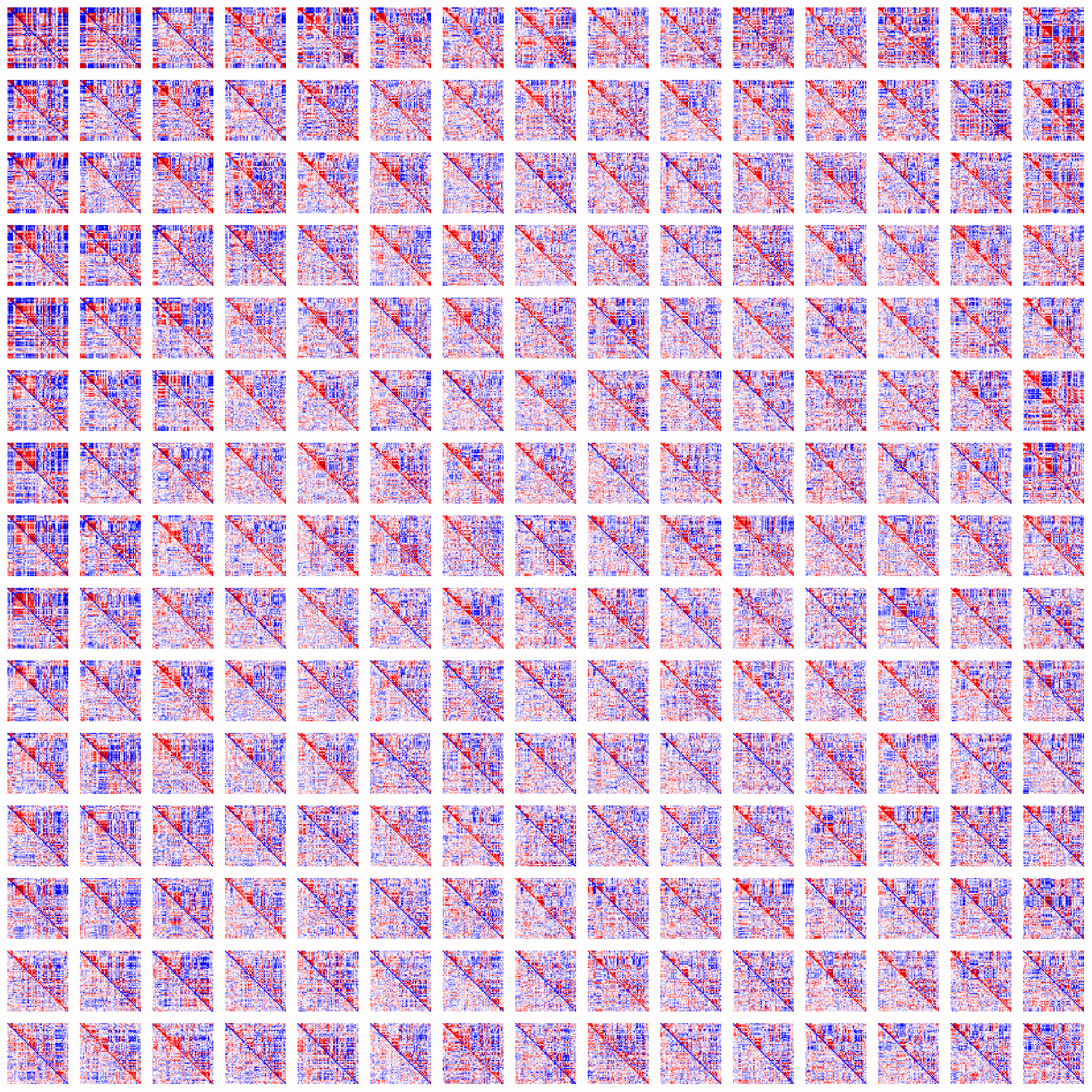
ABIDE I original sFNC (training).

**Figure H.7:**
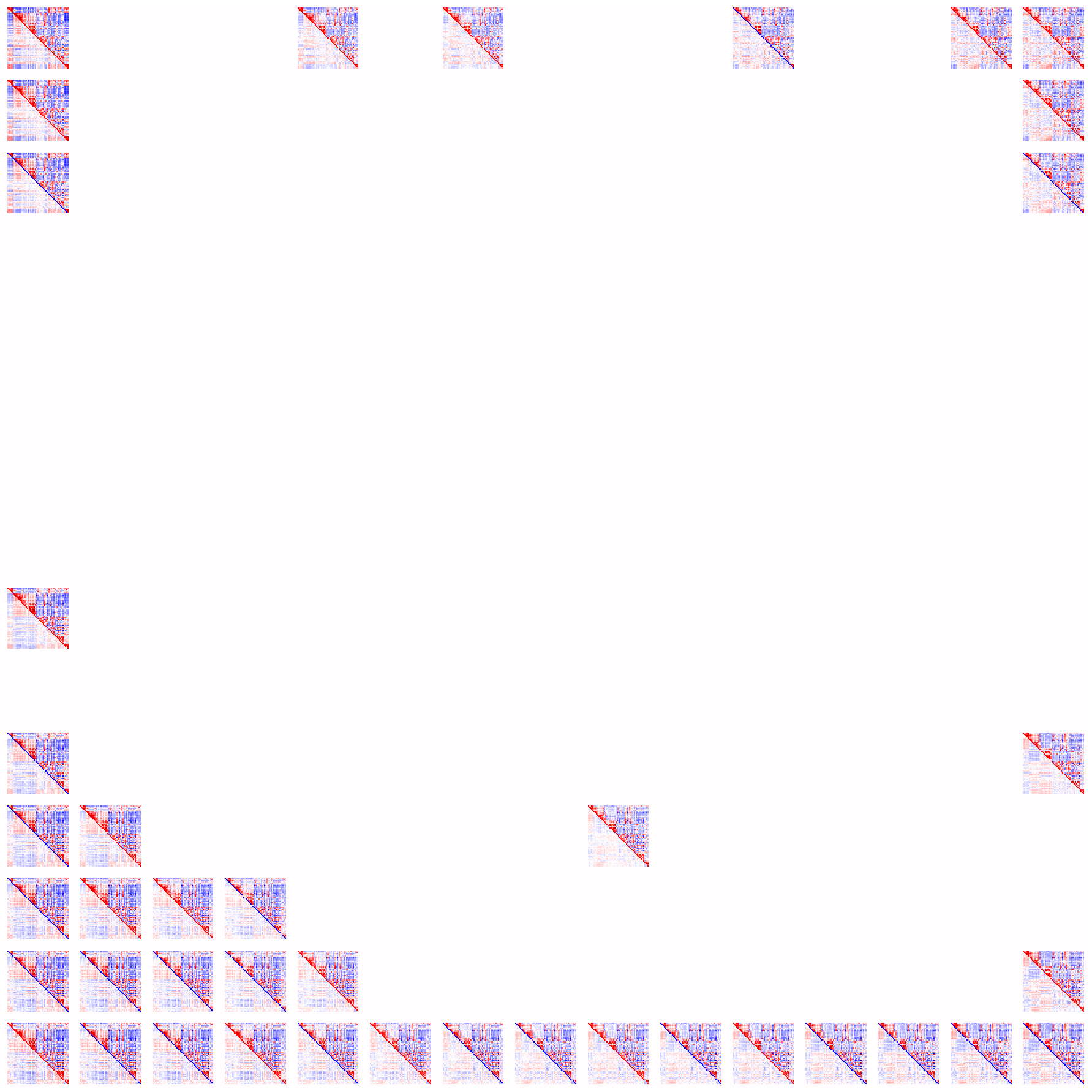
ABIDE I generated sFNC (test).

**Figure H.8:**
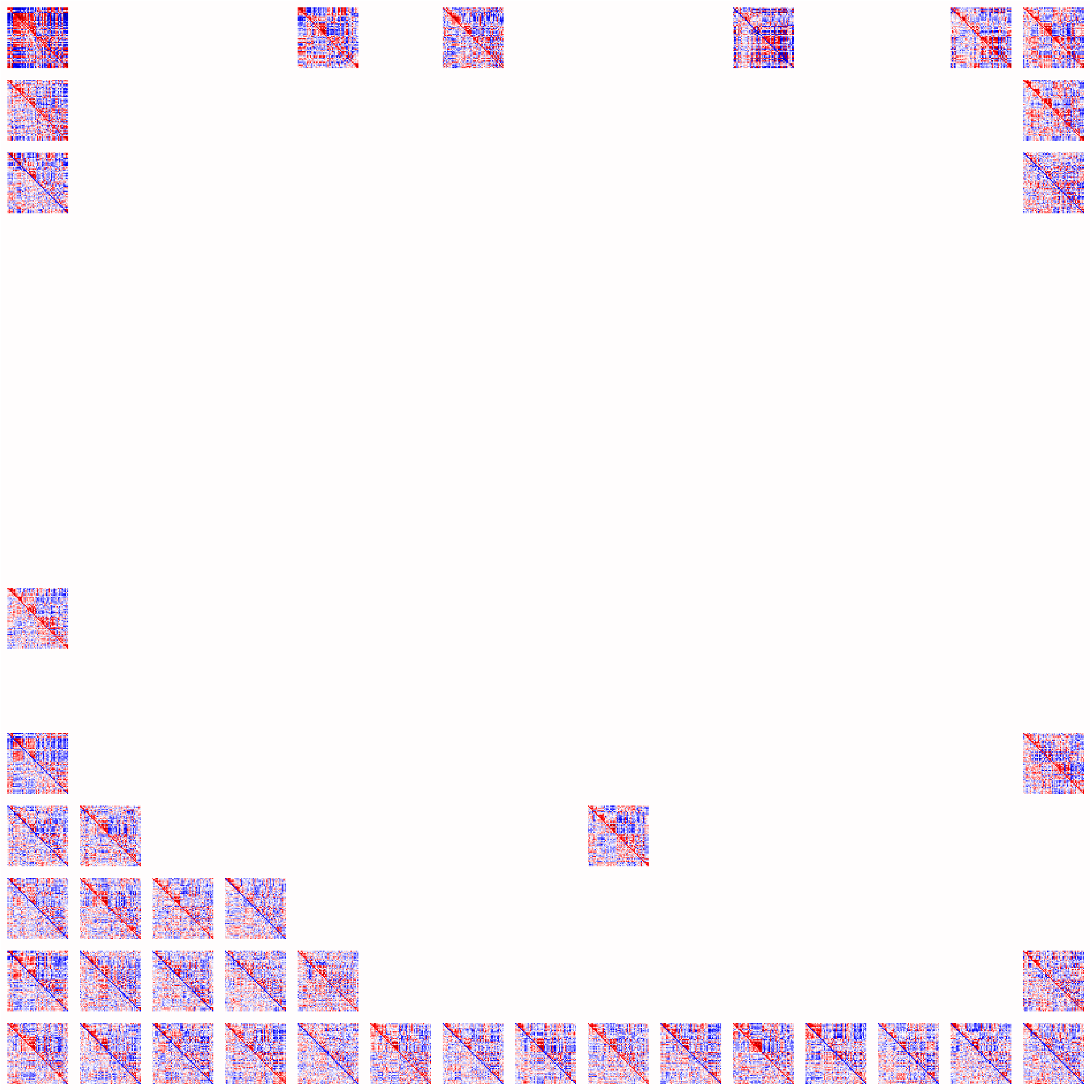
ABIDE I original sFNC (test).

## Appendix I Group differences in sFNC variability

To quantify heterogeneity in sFNC, we computed the cell-wise standard deviation of sFNC for each group and each dataset (Figure I.1). In the FBIRN dataset, the average standard deviation was slightly higher in the patient group (0.201) than in the control group (0.199). A similar pattern was observed in ABIDE I, where the patient group also showed a marginally higher average standard deviation (0.185) compared with controls (0.183). These results indicate slightly greater sFNC variability in the patient group across both datasets.

**Figure I.1:**
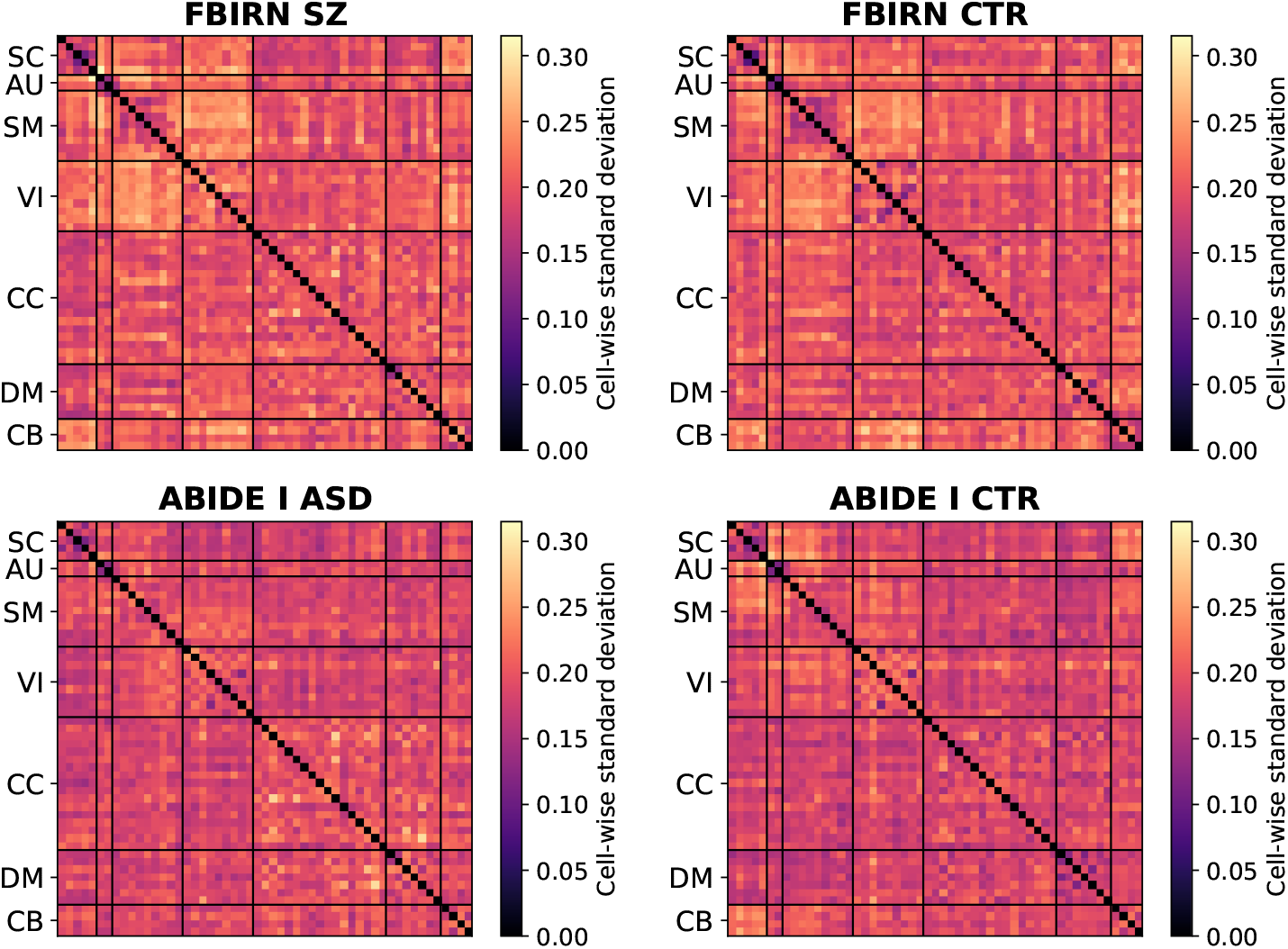
sFNC cell-wise standard deviation. In both FBIRN and ABIDE I datasets, the average cell-wise standard deviation in the patient group is slightly higher than that in the control group, indicating slightly greater sFNC variability in the patient group.

## Appendix J Subgroup analysis using two- and three-dimensional latent representations

We examined how the dimensionality of the VAE latent space affects downstream subgroup identification within the SZ patient cohort. First, we trained two VAEs to learn latent representations, one with a two-dimensional (2D) latent space and one with a three-dimensional (3D) latent space. Second, for each set of latent representations, we applied *k*-means clustering, varying the number of clusters *k* from 2 to 9. Third, we selected the optimal number of clusters using the elbow criterion based on the *L*_2_ loss. Finally, we assessed whether pairs of clusters differed significantly in subject measures, including PANSS positive and negative scores, CMINDS cognitive scores, age, and gender.

Based on the elbow criterion, four clusters were identified for the 2D representations and six clusters for the 3D representations (Figure J.1). Figures J.2 and J.5 show the resulting subgroup (cluster) assignments and the corresponding average sFNC pattern for each subgroup within each diagnostic group. Figures J.3 and J.6 illustrate the corresponding p-values for subject measures between subgroup pairs. Figures J.4 and J.7 then display distributions of subject measures across subgroups. Statistical relationships among subgroup pairs derived from the 2D latent space are largely consistent with those reported in Table 2. For both 2D and 3D representations, fifteen subgroup pairs exhibited statistically significant differences (*p <* 0.05, Mann–Whitney U test), suggesting that the higher-dimensional (3D) latent space does not yield additional meaningful subgroups compared with the 2D latent space.

Furthermore, we examined FNC patterns across four subgroups derived from 2D latent representations (Figure J.2). Table J.1 summarizes pairwise subgroup differences in subject measures and functional network connectivity. Subgroups 1 and 2 differed significantly in age and attention scores (*p <* 0.05, Mann–Whitney U test), with associated FNC changes primarily involving sensorimotor and cognitive control networks, including SC-SM, AU-CC, SM-VI, SM-DM, and CC-CB domains (*p <* 0.0001, Wilcoxon signed-rank test with Bonferroni correction). Subgroups 1 and 3 showed significant differences in age, visual learning scores, and composite scores. Statistically significant differences were observed in both within-network alterations (SM, VI) and between-network interactions. Subgroups 1 and 4 differed significantly in visual learning scores, with FNC differences dominated by visual network connectivity (VI VI, VI-CB, VI-SM, VI-SC) and subcortical, sensorimotor, and cerebellar interactions (SC-SM, SC-CB, SM-CB). Subgroups 2 and 3 were significantly different in visual learning and reasoning scores, with FNC differences spanning within-network connectivity (SM, VI, and CC) and between-network interactions involving subcortical, sensorimotor, visual, and cerebellar-related connections. Subgroups 2 and 4 differed in working memory and visual learning scores, particularly among subcortical, sensorimotor, visual, and cerebellar domains. Subgroups 3 and 4 exhibited multiple cognitive score differences, including speed of processing, working memory, verbal learning, visual learning, and composite scores, with FNC differences mainly involving sensorimotor and visual networks. Table J.2 summarizes pairwise subgroup differences in subject measures and functional network connectivity patterns using 3D latent representations.

**Figure J.1:**
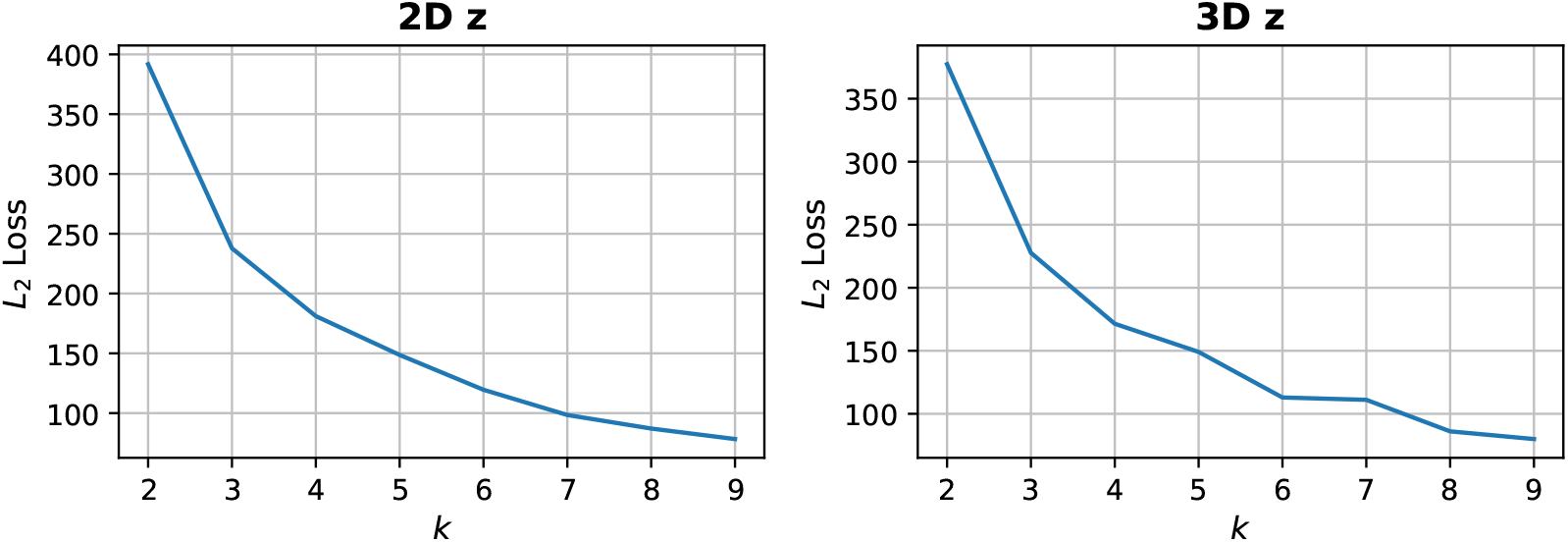
*k*-means clustering *L*_2_ loss with respect to different *k* values (2D vs 3D). According to the elbow criterion, four clusters were identified for the 2D representations and six clusters for the 3D representations.

**Figure J.2:**
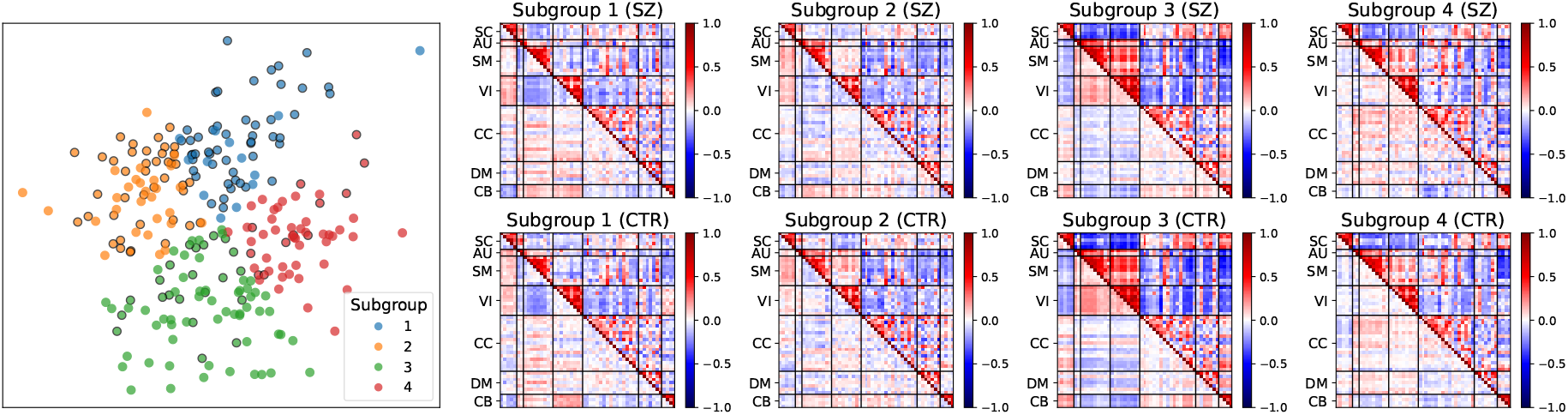
Subgroups based on 2D latent representations. In the scatter plot, SZ patients are highlighted in black circles, while controls are shown in their original colors. The average sFNC pattern is shown for each of the four *k*-means subgroups (clusters) and for each of the two diagnostic groups (SZ patients and controls). Subgroups were sorted by the increasing proportion of controls among all subjects.

**Figure J.3:**
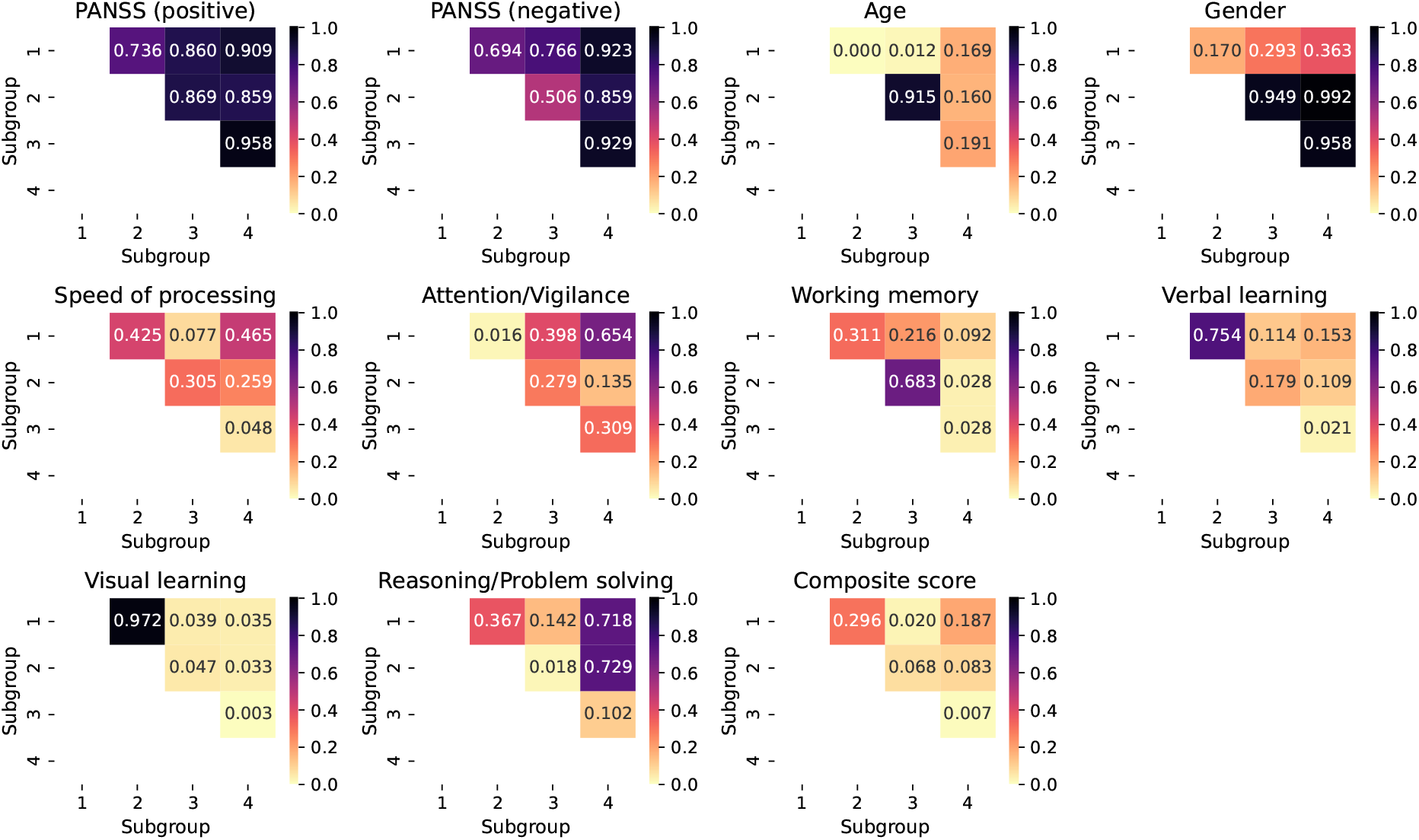
Statistical significance of each score type between two SZ subgroups (2D latent representations). The heatmaps show p-values for each score type between all pairs of subgroups (Mann–Whitney U test). Fifteen cluster pairs exhibit statistically significant differences (*p <* 0.05).

**Figure J.4:**
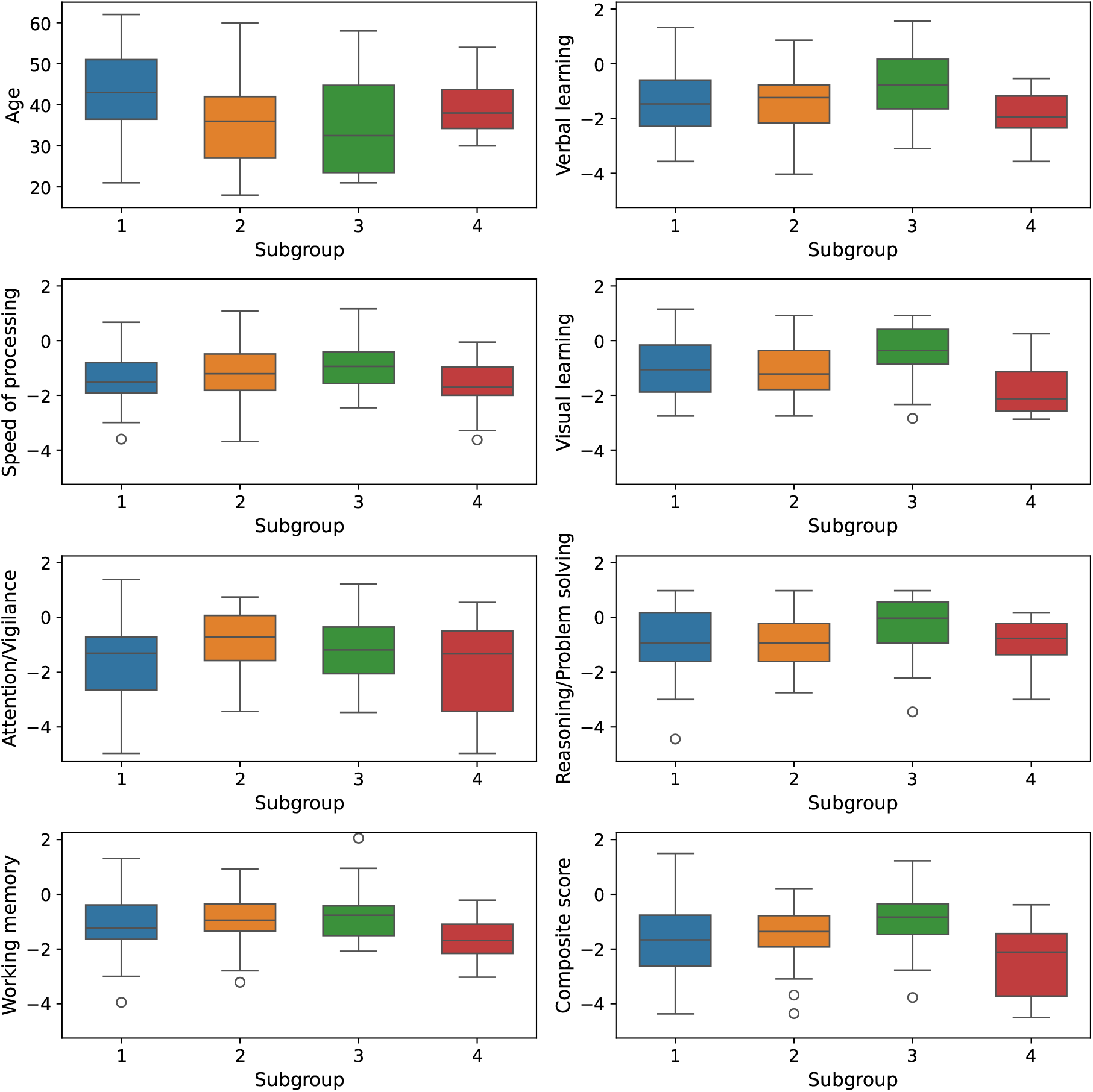
Subject measure distributions for SZ subgroups (2D latent representations). Box plots display subject measure (age, cognitive score) distributions (quartiles) for SZ subgroups.

**Figure J.5:**
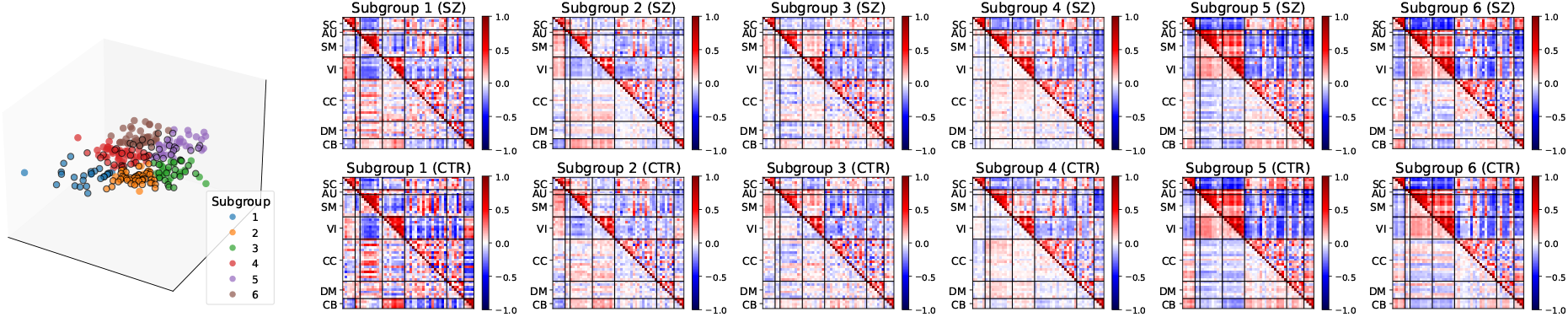
Subgroups based on 3D latent representations. In the scatter plot, SZ patients are highlighted in black circles, while controls are shown in their original colors. The average sFNC pattern is shown for each of the four *k*-means subgroups (clusters) and for each of the two diagnostic groups (SZ patients and controls). Subgroups were sorted by the increasing proportion of controls among all subjects.

**Figure J.6:**
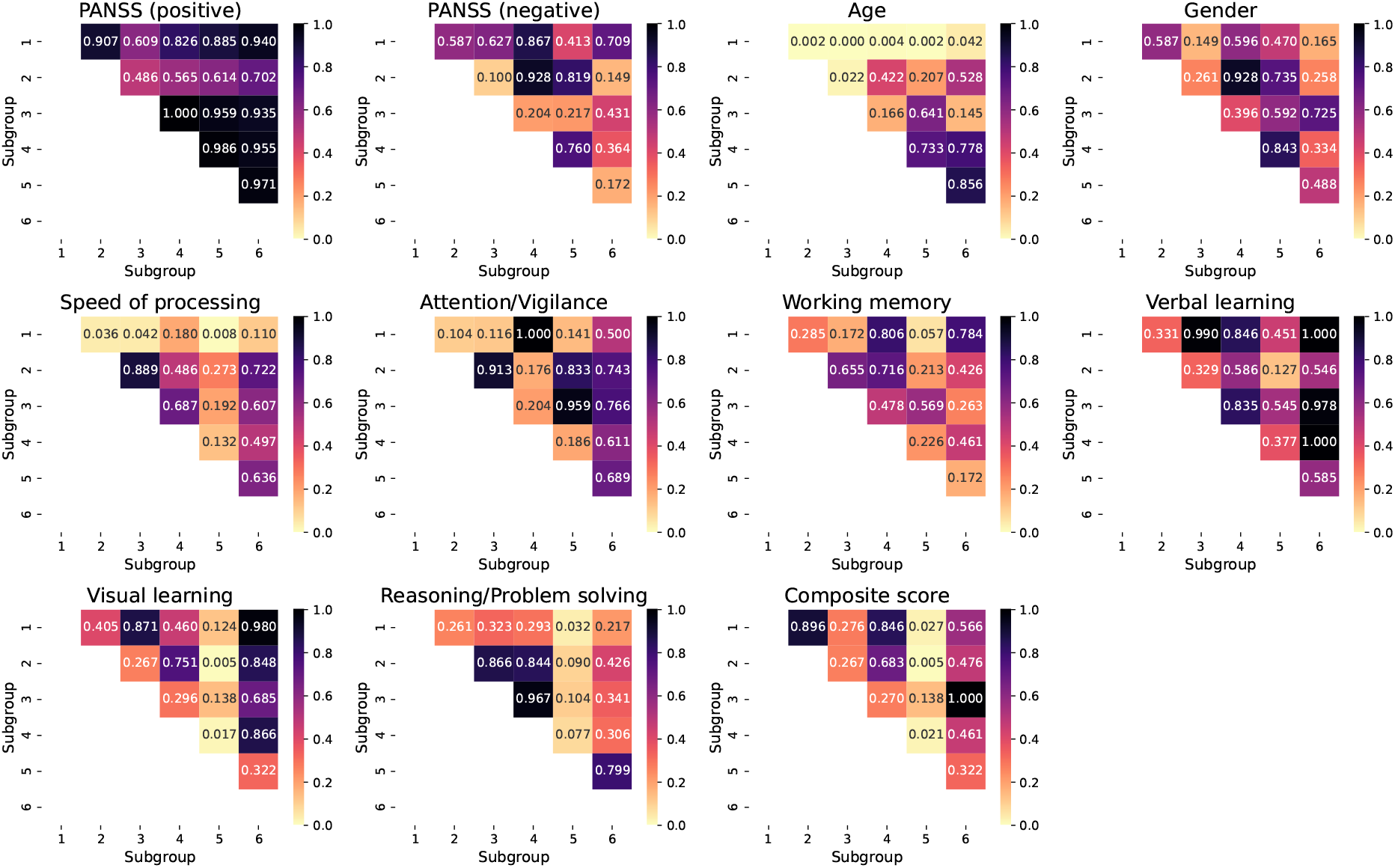
Statistical significance of each score type between two subgroups of 3D latent representations in the patient group. The heatmaps show p-values for each score type between all pairs of subgroups (Mann–Whitney U test). Fifteen cluster pairs exhibit statistically significant differences (*p <* 0.05).

**Figure J.7:**
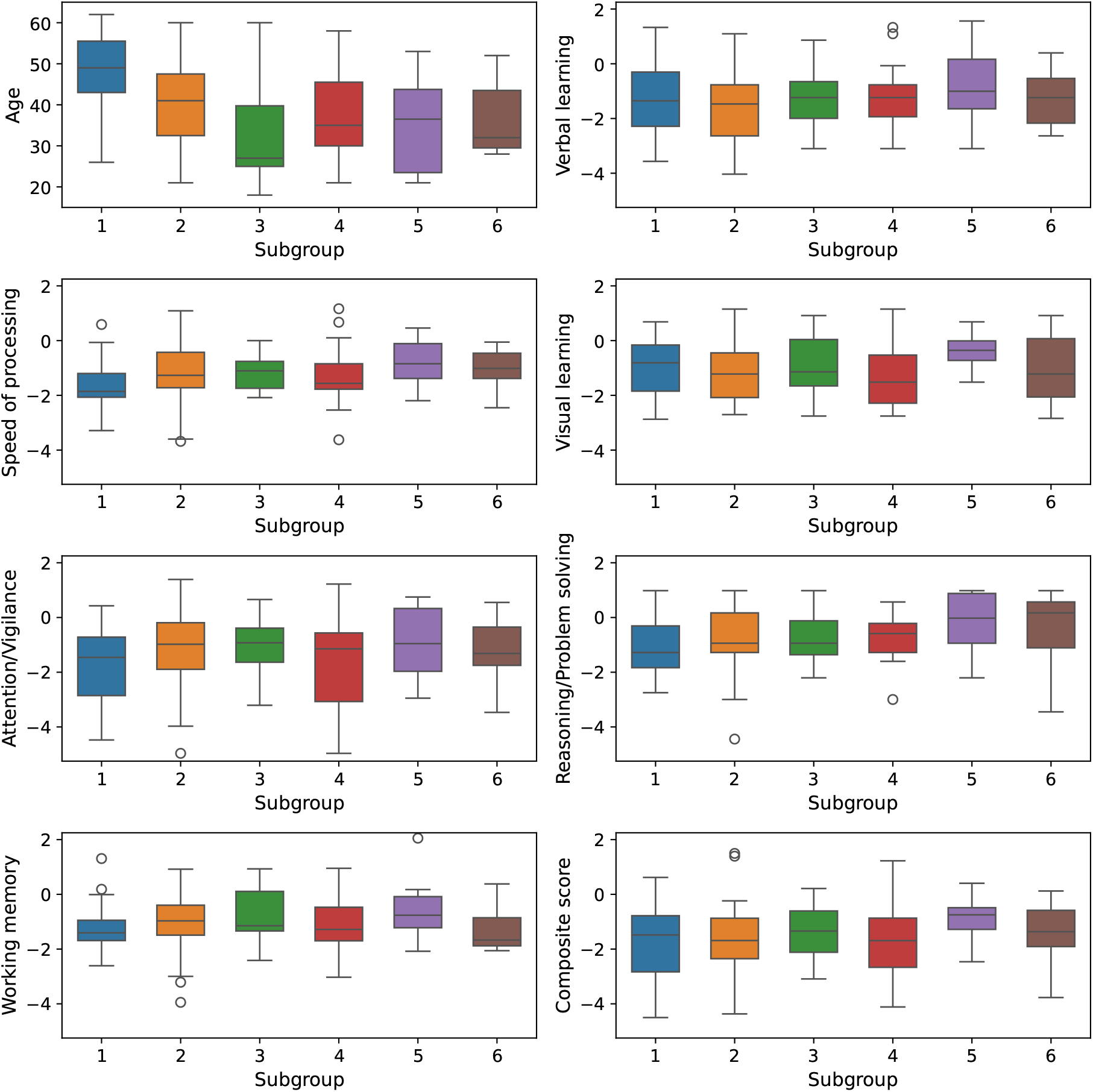
Subject measure distributions for SZ subgroups (3D latent representations). Box plots display subject measure (age, cognitive score) distributions (quartiles) for SZ subgroups.

**Table J.1:** Pairwise subgroup differences in subject measures and functional network connectivity (3D latent representations). Each row lists a subgroup pair, significantly different subject measures (*p <* 0.05, Mann–Whitney U test), and corresponding functional network connectivity patterns (*p <* 0.05, Wilcoxon signed-rank test with Bonferroni correction).

| Subgroup pairs | Subject measures | Functional network connectivity |
| --- | --- | --- |
| 1, 2 | Age<br>Attention | SC-SM, AU-CC, SM-VI, SM-DM, CC-CB ($p < 0.0001$);<br>AU-SM, SM-CC ($p < 0.001$);<br>SC-CB, VI-CB, DM-CB ($p < 0.01$);<br>SC-VI, AU-DM, CB-CB ($p < 0.05$) |
| 1, 3 | Age<br>Visual learning<br>Composite score | SC-SM, SC-VI, SC-CB, SM-SM, SM-VI,<br>SM-CC, SM-DM, SM-CB, VI-VI, VI-CC,<br>VI-CB, CC-CB, DM-CB ($p < 0.0001$);<br>AU-SM, AU-VI, AU-CC ($p < 0.001$);<br>SC-SC ($p < 0.01$);<br>SC-AU, SC-CC, CC-CC ($p < 0.05$) |
| 1, 4 | Visual learning | SC-SM, SC-VI, SC-CB, SM-VI, SM-CB,<br>VI-VI, VI-CB ($p < 0.0001$);<br>CC-CC, CB-CB ($p < 0.05$) |
| 2, 3 | Visual learning<br>Reasoning | SC-SM, SC-VI, SC-CB, SM-SM, SM-VI,<br>SM-CB, VI-VI, VI-CC, VI-CB, CC-CC ($p < 0.0001$);<br>SC-CC, AU-VI ($p < 0.001$);<br>SC-SC, AU-SM, SM-CC, CC-CB ($p < 0.01$);<br>SC-AU, CB-CB ($p < 0.05$) |
| 2, 4 | Working memory<br>Visual learning | SC-SM, SC-VI, SM-VI, SM-DM, SM-CB,<br>VI-VI, VI-CB, CC-CB ($p < 0.0001$);<br>AU-CC, SM-CC ($p < 0.001$);<br>SC-CB, SM-SM, CC-CC, DM-DM ($p < 0.01$);<br>SC-SC, AU-DM, CB-CB ($p < 0.05$) |
| 3, 4 | Speed of processing<br>Working memory<br>Verbal learning<br>Visual learning<br>Composite score | SC-SM, SC-VI, SC-CB, AU-CC, SM-SM,<br>SM-VI, SM-CC, SM-DM, SM-CB, VI-VI,<br>VI-CC, VI-CB, CC-CB, DM-CB ($p < 0.0001$);<br>SC-CC, AU-SM, AU-VI, DM-DM ($p < 0.001$);<br>AU-DM ($p < 0.01$);<br>SC-SC, SC-AU, VI-DM, CB-CB ($p < 0.05$) |

**Table J.2:** Pairwise subgroup differences in subject measures and functional network connectivity (2D latent representations). Each row lists a subgroup pair, significantly different subject measures (*p <* 0.05, Mann–Whitney U test), and corresponding functional network connectivity patterns (*p <* 0.05, Wilcoxon signed-rank test with Bonferroni correction).

| Subgroup pairs | Subject measures | Functional network connectivity |
| --- | --- | --- |
| 1, 2 | Age<br>Speed of processing | SC-SM, SC-VI, AU-CC, SM-VI, SM-CB, CC-CB ($p < 0.0001$);<br>AU-SM, VI-VI ($p < 0.001$);<br>SM-CC, SM-DM ($p < 0.01$);<br>SM-SM, CB-CB ($p < 0.05$) |
| 1, 3 | Age<br>Speed of processing | SC-SM, SC-VI, AU-CC, SM-VI, SM-DM, VI-CB,<br>CC-CB ($p < 0.0001$);<br>AU-SM, SM-CC ($p < 0.001$);<br>SC-CB, DM-DM, DM-CB ($p < 0.01$);<br>AU-DM, CB-CB ($p < 0.05$) |
| 1, 4 | Age | SC-VI, SM-VI, SM-CB, VI-VI, VI-CB, CC-CB ($p < 0.0001$);<br>AU-SM, DM-CB ($p < 0.001$);<br>SC-SC, SC-CB ($p < 0.01$);<br>AU-VI, SM-DM, CB-CB ($p < 0.05$) |
| 1, 5 | Age<br>Speed of processing<br>Reasoning<br>Composite score | SC-VI, SC-CB, AU-CC, SM-VI, SM-CC, SM-DM,<br>SM-CB, VI-VI, VI-CC, VI-CB, CC-CB, DM-CB ($p < 0.0001$);<br>AU-SM, AU-VI ($p < 0.001$);<br>SC-SC, SC-SM ($p < 0.01$);<br>SC-AU, CB-CB ($p < 0.05$) |
| 1, 6 | Age | SC-SM, SC-VI, SC-CB, SM-VI, SM-CC, SM-CB,<br>VI-VI, VI-CB, CC-CB, DM-CB ($p < 0.0001$);<br>AU-SM, AU-VI ($p < 0.001$);<br>SC-SC, VI-CC ($p < 0.01$) |
| 2, 3 | Age | SC-VI, SC-CB, SM-VI, SM-DM, SM-CB, VI-VI,<br>VI-DM, VI-CB, CC-CB, DM-DM, DM-CB ($p < 0.0001$);<br>AU-SM, AU-VI, SM-CC ($p < 0.001$);<br>SC-SC, SC-DM, AU-DM ($p < 0.01$);<br>SC-AU, SC-SM, CB-CB ($p < 0.05$) |
| 2, 5 | Visual learning<br>Composite score | SC-SM, SC-VI, SC-CB, SM-SM, SM-VI, SM-CC,<br>SM-CB, VI-VI, VI-CC, VI-CB, CC-CC, CC-CB ($p < 0.0001$);<br>SC-CC, AU-SM, AU-VI, AU-CC, DM-CB ($p < 0.001$);<br>SC-SC, SM-DM ($p < 0.01$);<br>SC-AU, SC-DM, CB-CB ($p < 0.05$) |
| 4, 5 | Visual learning<br>Composite score | SC-SM, SC-VI, SC-CC, SC-CB, AU-CC, SM-SM,<br>SM-VI, SM-CC, SM-DM, SM-CB, VI-VI, VI-CC,<br>VI-CB, CC-CB ($p < 0.0001$);<br>AU-SM, AU-VI, DM-CB ($p < 0.001$);<br>AU-DM ($p < 0.01$);<br>SC-AU, SC-DM, VI-DM, DM-DM ($p < 0.05$) |

## Appendix K sFNC interpolation along cognitive score trajectories

To examine how sFNC patterns vary across cognitive performance levels, we divided subjects in the FBIRN dataset into two groups based on the median cognitive score: a lower-score group (scores ≤ median) and a higher-score group (scores *>* median). Within each group, we derived a cluster in the latent space according to the distribution of cognitive scores. For each pair of clusters (higher- and lower-score groups), we constructed a trajectory connecting their centroids in the latent space. Finally, we generated one representative sFNC matrix from each cluster centroid to characterize connectivity patterns associated with different cognitive levels. Most cognitive scores show moderate to strong correlations (Figure K.1), therefore, the resulting clusters and trajectories are largely similar across different cognitive measures (Figure K.2).

**Figure K.1:**
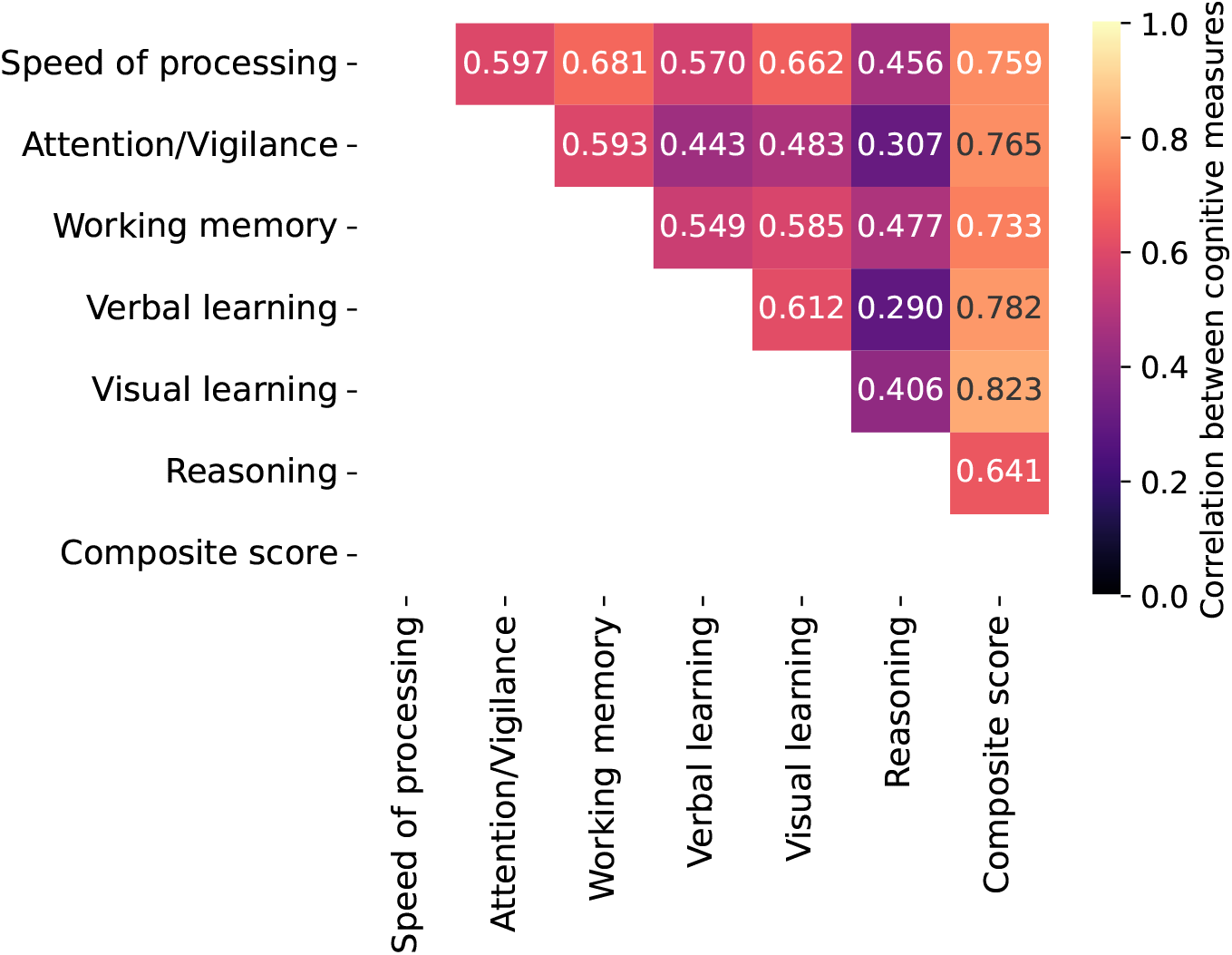
Correlations between cognitive measures. Cognitive scores exhibit varying degrees of similarity, with correlation coefficients ranging from 0.290 to 0.823. The composite score shows strong correlations with the other scores.

As shown in Figure K.2, clusters corresponding to different cognitive score groups occupy highly consistent spatial locations in the latent space, with the lower-score cluster located in the upper-left region and the higher-score cluster in the lower-right region. For each cluster, we generated one representative sFNC matrix by sampling from the cluster centroid. For each group, we also computed the average sFNC matrix across the original sFNC matrices. We demonstrate that, for each group, the generated sFNC matrices are highly similar to the corresponding original sFNC matrices (Figure K.3).

Across the two sampled states, we observe systematic and consistent changes in sFNC patterns associated with cognitive performance. From group 1 (with higher scores) to group 2 (with lower scores), negative correlations are significantly attenuated in the SC–SM, SC–VI, and SM–CB domains (*p <* 0.0001, Wilcoxon signed-rank test with Bonferroni correction). We also observe a transition from positive to negative correlations in the SC–CB and SM–VI domains (*p <* 0.0001), as well as a transition from negative to positive correlations in the VI–CB domain (*p <* 0.0001) and the SC–AU domain (*p <* 0.05). In addition, positive correlations within the SM, VI, and CC domains are significantly reduced (*p <* 0.0001), as are correlations within the SC domain and the AU–SM domain (*p <* 0.01), and within the CB domain (*p <* 0.05).

Several connectivity changes are specific to individual cognitive domains. For attention (*p <* 0.05), working memory (*p <* 0.05), and reasoning (*p <* 0.01), we observe a shift from positive to negative correlations in the AU–VI domain. Moreover, working memory (*p <* 0.001) and reasoning (*p <* 0.0001) scores are associated with significantly weakened negative correlations in the VI–CC domain.

**Figure K.2:**
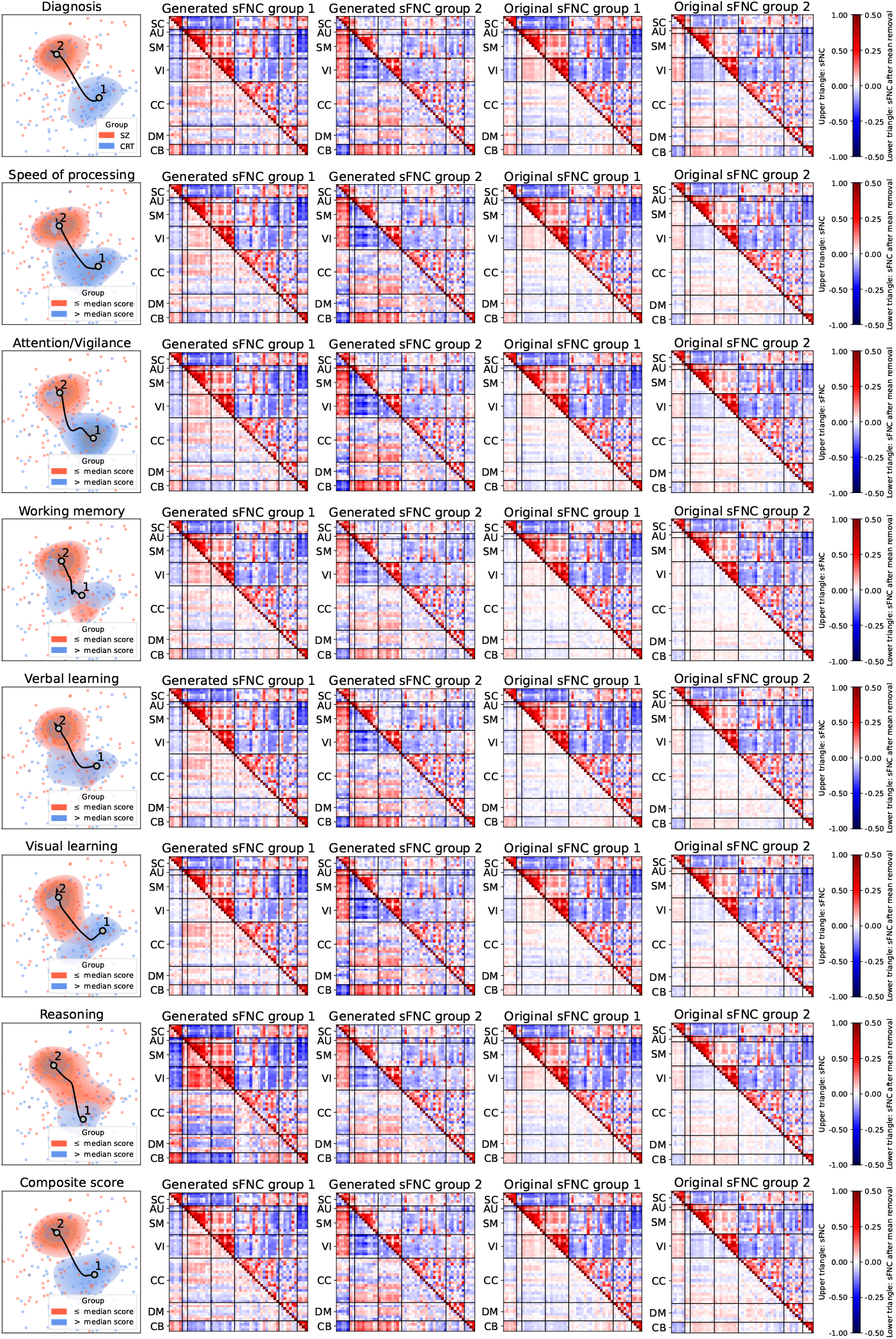
Interpolation along a trajectory derived from two clusters defined by cognitive scores. In each row, the scatter plot shows latent features colored by cognitive scores (red: ≤ median score; blue: *>* median score), overlaid with contour plots of two clusters connected by a trajectory. For each group, we generated one representative sFNC matrix by sampling from the cluster centroid. In addition, we computed the average sFNC matrix across the original sFNC matrices.

**Figure K.3:**
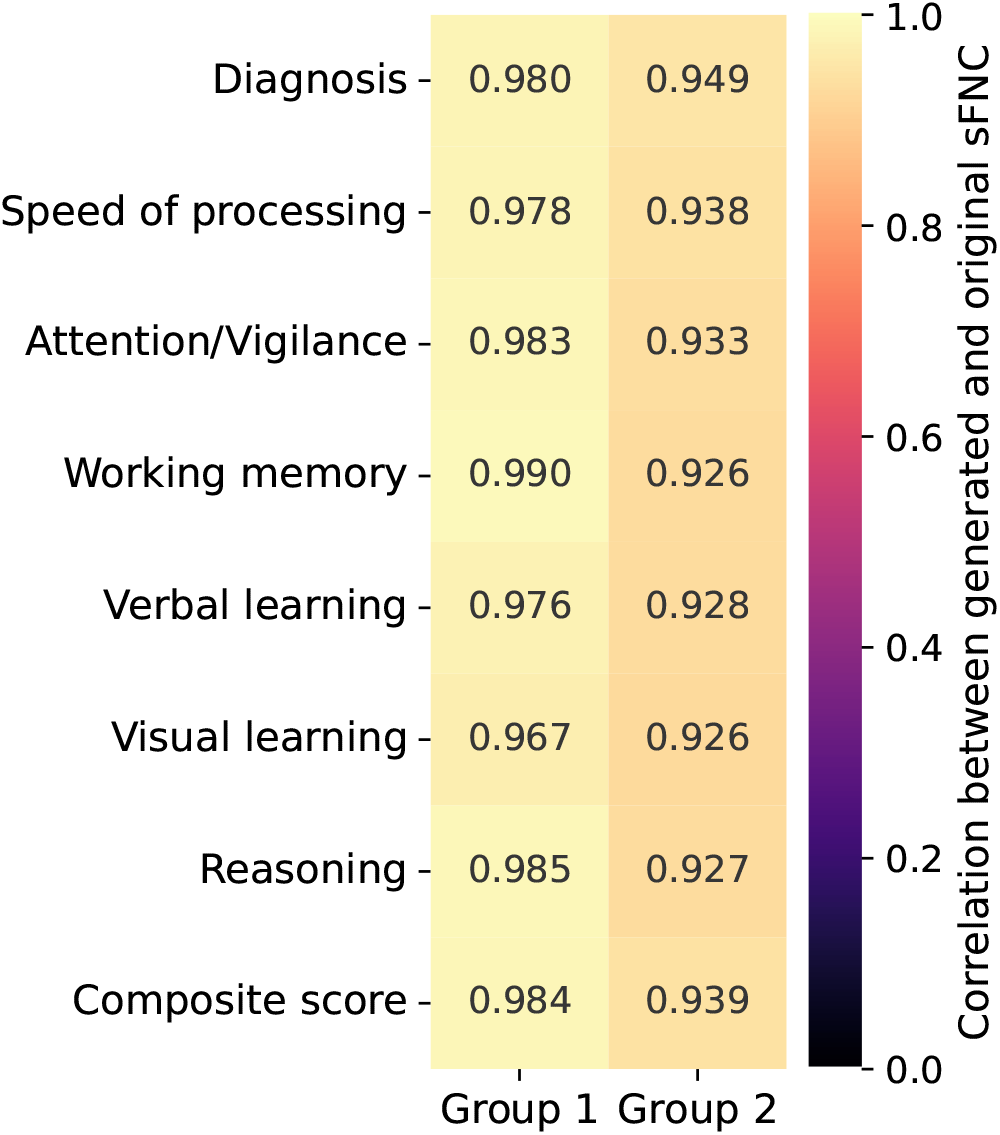
Correlations between generated and original sFNC. For each group, the generated sFNC matrices are highly similar to the corresponding original sFNC matrices.

## Appendix L Similarity measured by mean squared error

Mean squared error (MSE) was used to measure the similarity between the generated and original sFNC matrices. The median MSEs for the FBIRN training set, FBIRN test set, ABIDE I training set, and ABIDE I test set are 0.031, 0.034, 0.029, and 0.034, respectively. These results highlight that the generated sFNC matrices are highly similar to the original ones, as measured by MSE.

**Figure L.1:**
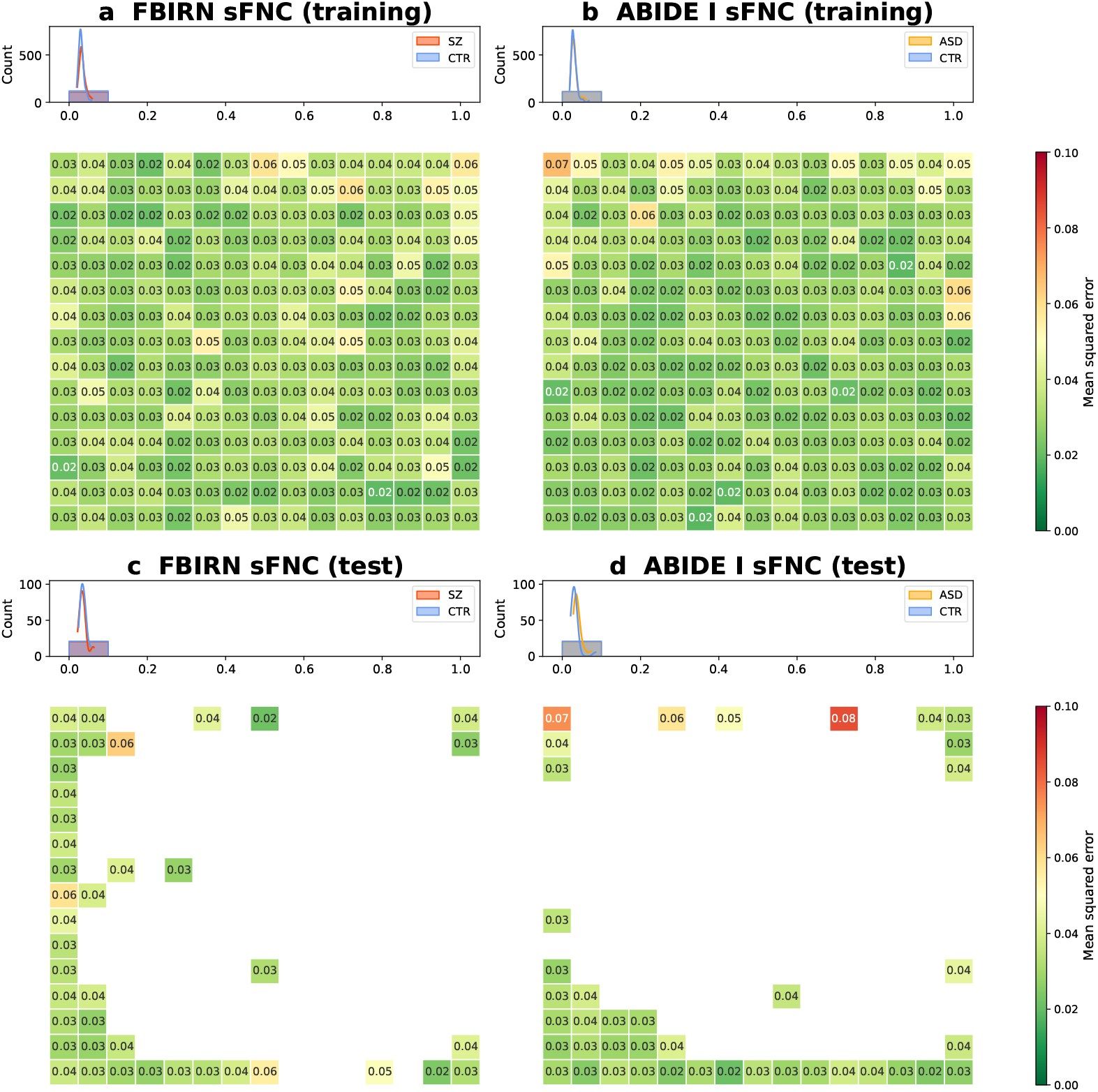
Mean squared errors between generated and original sFNC matrices. (a) FBIRN training set. (b) ABIDE I training set. (c) FBIRN test set. (d) ABIDE I test set. Each panel shows mean squared errors (MSEs) between individual generated and original sFNC matrices on the 2D grid and the corresponding group-specific histogram distributions. Low MSEs across all panels suggest a high degree of correspondence between generated and original sFNC matrices.

## Appendix M k-means clustering

The *k*-means clustering algorithm was separately applied to generated or original dFNC data in each dataset to identify dynamic states by searching the number of states *k* from 2 to 9. The optimal number of states was then identified using the elbow criterion based on the *L*_2_ loss, which measures the Euclidean distance between each sample and its assigned centroid. As shown in Figure M.1, the elbow point occurred at *k* = 5 in most cases (Figure M.1a,b,c). Thus, an optimal value *k* = 5 was used for both FBIRN and ABIDE I.

**Figure M.1:**
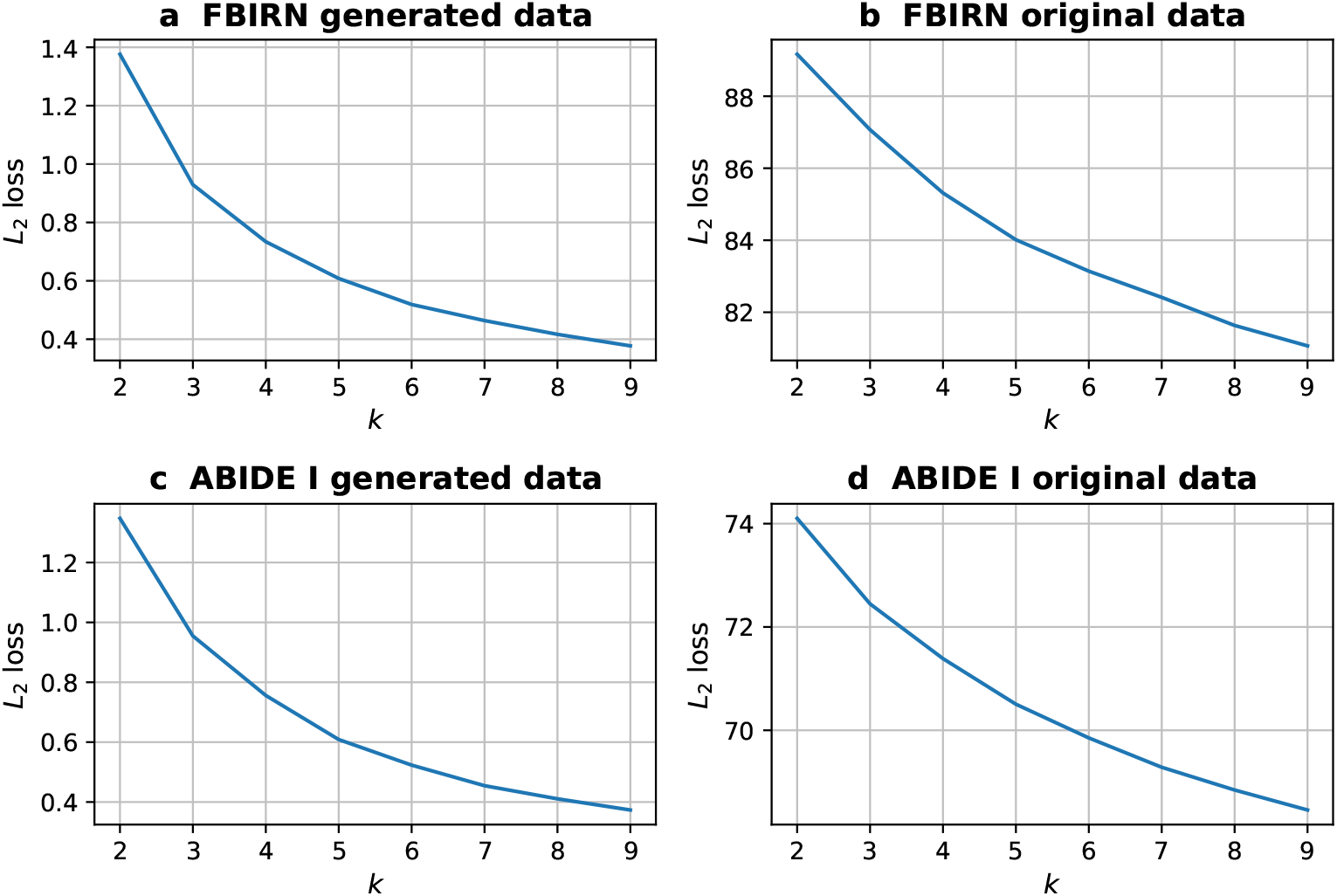
*k*-means clustering *L*_2_ loss with respect to different *k* values. (a) FBIRN generated data. (b) FBIRN original data. (c) ABIDE I generated data. (d) ABIDE I original data.

## Footnotes

1 A statistical model is identifiable if the mapping from each set of its parameters to the distribution of observed data is injective (Lehmann & Casella, 2006).

2 The KL divergence measures how a probability distribution differs from another probability distribution. For a discrete random variable, 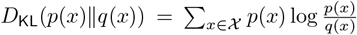; for a continuous random variable, 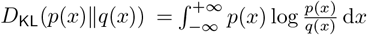.

## Notes

### Competing Interest Statement

The authors have declared no competing interest.

### Summary of Updates

In this revision, we added new appendix sections on subgroup analyses and cognitive score trajectory interpolation.

